# Tumor-associated macrophages enhance tumor innervation and spinal cord repair

**DOI:** 10.1101/2024.12.19.629374

**Authors:** Sissi Dolci, Loris Mannino, Alessandra Campanelli, Eros Rossi, Emanuela Bottani, Francesca Ciarpella, Isabel Karkossa, Elisa Setten, Benedetta Savino, Giulia Pruonto, Nicola Piazza, Stefano Gianoli, Alessia Amenta, Giuseppe Busetto, Alex Pezzotta, Marzia Di Chio, Alessandra Castagna, Nicolò Martinelli, Ilaria Barone, Federico Boschi, Adam Doherty, Maria Teresa Scupoli, Chiara Cavallini, Giorgio Malpeli, Zulkifal Malik, Ludovica Sagripanti, Vincenzo Silani, Patrizia Cristofori, Eugenio Scanziani, Marco Sandri, Anna Pistocchi, Patrizia Bossolasco, Marco Endrizzi, Kristin Schubert, Guido Francesco Fumagalli, Massimo Locati, Francesco Bifari, Ilaria Decimo

## Abstract

Tumor-associated macrophages (TAM) enhance cancer progression by promoting angiogenesis, extracellular matrix (ECM) remodeling, and immune suppression. Nerve infiltration is a hallmark of various cancers and is known to directly contribute to tumor growth. However, the role of TAM in promoting intratumoral nerve growth remains poorly understood. In this study, we demonstrate that TAM expressed a distinct “neural growth” gene signature. TAM actively enhance neural growth within tumors and directly promote neurites outgrowth. We identify secreted phosphoprotein 1 (Spp1) as a key mediator of TAM-driven neural growth activity, which triggers neuronal mTORC2 signaling.

Leveraging this new neural growth function, which added to the TAM wound healing properties, we explored TAM potential to repair central nervous system. Adoptive transfer of *in vitro*-generated TAM in a severe complete-compressive-contusive spinal cord injury (scSCI) model, not only repaired the damaged neural parenchyma by improving tissue oxygenation, ECM remodeling, and dampening chronic inflammation, but also resulted in neural regrowth and partial functional motor recovery. Proteomic analysis and subsequent functional validation confirmed that TAM-induced spinal cord regeneration is mediated through the activation of neural mTORC2 signaling pathways.

Collectively, our data unveil a previously unrecognized role of TAM in tumor innervation, neural growth, and neural tissue repair.

## Introduction

Intratumoral nerve infiltration supports cancer growth in several tumors, including pancreatic, breast, endometrial, and colon-rectal tumors (1). Intriguingly, many tumors promote nerve growth and infiltration, generating a positive feed-back loop where the cancer-dependent nerve growth further supports tumor progression (1). Several mediators and mechanisms have been proposed to regulate nerve-cancer interaction (2, 3). *In vitro* and *in vivo* experiments have identified different neurotrophins, growth factors, and axon guidance molecules produced by cancer cells that contribute to cancer-dependent nerve growth (4). Conversely, the contribution of different cells composing the tumor microenvironment to cancer-dependent nerve growth has been poorly investigated. In particular, tumor-associated macrophages (TAM) have been shown to play a crucial impact on tumor growth stimulation supporting angiogenesis, extracellular matrix (ECM) remodeling, cell proliferation, and immunosuppression (5, 6). Macrophages and other cells of the tumor microenvironment, including Schwann cells, have been shown to migrate through nerves, functioning as chemoattractants for cancer cells (7, 8). Nonetheless, the specific role of TAM in directly stimulating nerve growth remains unclear. Beyond their role in organogenesis, tissue regeneration (9–12), and wound healing (13), macrophages also contribute to peripheral nerve maintenance and regeneration (9, 14–16). Resident enteric microglia-like nerve-associated macrophages have been shown to play a role in developmental synaptic pruning and maintenance of mature neuronal circuitry (17–20). On the opposite, macrophage-mediated healing/regenerative functions have been reported to be defective in the injured adult CNS regeneration, characterized by the development of a maladaptive wound repair process supported by tissue hypoxia, chronic inflammation, and fibrotic cysts formation leading to neural tissue loss and axonal degeneration (10–12, 21). In this work, we investigated the neural growth properties of TAM and their relevance in tumor progression and neuroregeneration after central nervous system (CNS) injury, such as spinal cord injury (SCI).

## Results

### Tumor associated macrophages (TAM) express a “neural growth” gene signature

To assess whether TAM are endowed with properties related to neural growth potential, we first investigated the gene signature of TAM infiltrating human tumor in our single-cell RNA sequencing (scRNAseq) data obtained from human glioblastoma (GBM), astrocytoma, and adjacent healthy tissue (Fig. 1A, B; Fig. S1A, B). We further extend the analysis of TAM signature to publicly available scRNAseq datasets of human pancreatic (22), breast (23), and colorectal cancers (24) (Fig. 1C, D). Analysis and visualization by Uniform Manifold Approximation and Projection (UMAP) showed that single-cell transcriptomes clearly identified a cluster of TAM (Fig. 1A, D) characterized by the expression of TGFBI^+^/CD163^+^ /CD14^+^/PTPRC^+^/ (Fig. 1B, D) (25, 26). To highlight TAM-specific features, we extracted, across all the datasets, the differential expressed genes (DEGs) of the TAM cluster compared to the CD14^-^/TMEM119^-^ cells. Subsequently, we conducted gene ontology (GO) analysis on the identified DEGs to characterize their functional annotations and associated biological processes. As expected, the TAM cluster upregulated GOs related to *response to hypoxia, angiogenesis, ECM remodeling*, and *wound healing* (Fig. S1C) (5, 6). Of note, we consistently found in TAM cluster of all scRNAseq datasets of human GBM, pancreatic, breast, and colorectal cancer, the upregulation of GOs related to *neural development, synapse, response to axon injury*, and *neuron/axon regeneration* (Fig. 1C, Fig. S1D) (27). To describe this new finding, in accordance with previous studies (25, 28–30), we first included all the DEGs of the TAM cluster in the GBM datasets composing these GOs (Fig. S1D), resulting in a list of 152 genes (see Material and Methods). We then refined this list by including only genes related to the cell secretome and itemised in the SEcreted Protein DataBase (SEPDB) (31), which resulted in a reduction of the list to 107 genes. We further manually refined this list to include only genes with well-established roles in neural growth-related functions, resulting in the definition of the “*neural growth*” gene list of 22 genes (Table S1). AUCell analysis revealed that the enrichment of the “*neural growth*” gene list was distinctively expressed in TAM clusters in all the analysed scRNA datasets (Fig. 1D).

**Legend to Figure 1:**
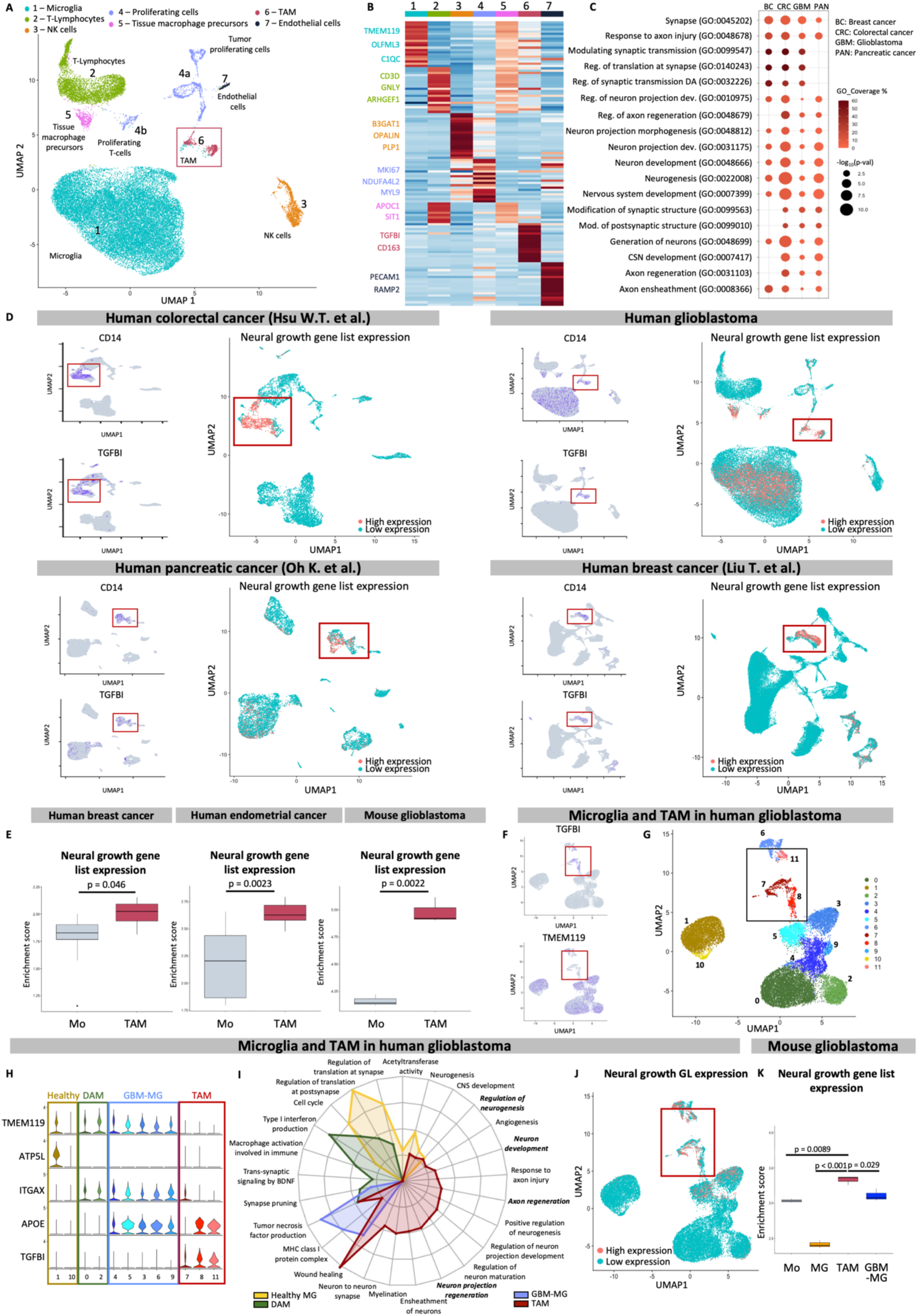
TAM show neural growth gene signature. **A)** Uniform Manifold Approximation and Projection (UMAP) plot on human GBM, astrocytoma, and healthy tissue cells. Cells were clustered in seven different groups. Red square indicates the TAM cluster. **B)** Heatmap showing the different gene expression among the clusters in Figure 1A and the gene marker used to identify the clusters. **C)** Bubble plot showing the expression of gene ontologies representing TAM neural growth signature in our GBM and public available breast, colorectal and pancreatic human scRNA datasets. The bubble color represents the coverage of the gene ontology, and the size represents the selected gene ontology’s p-value. **D)** (Left) UMAP plots showing CD14 and TGFBI expression (blue cells present the expression of the relative marker) in all the human scRNA datasets analyzed; (right) UMAP plot showing the AUCell analysis showing the increased expression of the “neural growth” gene list in TAM clusters (pink cells present higher expression of the genes included in the neural growth list). Red square indicates the TAM clusters. **E)** Single sample gene set enrichment analyses of the 22 genes included in the “neural growth” gene list of monocytes and TAM from human breast (monocytes: n=22; TAM: n=4) and endometrial cancer (monocytes: n=10; TAM: n=9), and of monocytes and TAM from mouse GBM (monocytes: n=3; TAM: n=3) showing enriched expression in TAM groups. **F, G)** UMAP plots of TAM and microglia cells included in the human GBM showing that clusters 7, 10, 11 are TAM cells (red square in G) identified as TGFBI^+^/ TMEM119^-^ (black square in F) and clusters 0-6 9,10 are microglia cells identified as TGFBI-/ TMEM119^+^ (blue cells in F represent the expression of the relative marker). **H)** Violin plot of marker genes describing healthy microglia (yellow cluster 1,10), disease-associated microglia (green clusters 0,2), glioblastoma-associated microglia (blue clusters 4,5,3,6,9), and TAM (red clusters 7, 10, 11) found in the subclustering of the GBM represented in G. **I)** Rounded gene ontologies plot showing that TAM (red) express GO related to neural growth which are not expressed by healthy microglia (yellow), disease-associated microglia (green), glioblastoma-associated microglia (blue). GO expression is reported as - log10(p-value). **J)** AUCell analysis of the “neural growth” gene list (pink cells present higher expression of the genes included in the nerve growth list) of TAM and microglia cells from the human scRNA of GBM, highlighting increased expression of the neural growth-related genes in TAM clusters. Red square indicates the TAM clusters. **K)** Single sample gene set enrichment analyses of “neural growth” gene list (22 genes) in monocytes, microglia, TAM, and disease-associated microglia (DAM) from mouse GBM (monocytes: n=4; microglia, TAM and DAM: n=3/condition), showing enriched expression in TAM groups. Statistical analyses were performed via Student t-test.

We further validated whether this “*neural growth*” gene signature was specifically expressed in TAM and not present in circulating monocytes (Mo). We, therefore, tested available bulk RNAseq datasets containing Mo and TAM isolated from human breast and endometrial cancer patients (GSE117970; (32)), and from mouse GBM models (PRJNA349180, (33)) (Fig. 1E). In line with results obtained from our scRNAseq analysis (Fig. 1D), TAM significantly enriched the “*neural growth*” list in human breast cancer (p=0.046; Fig. 1E), endometrial cancer (p=0.0023; Fig. 1E), and mouse GBM (p=0.0022, Fig. 1E) compared to Mo. Collectively, these findings confirmed that the *“neural growth”* gene signature is a consistent feature of TAM across various human and murine tumors (Fig. 1D, E).

TAM and microglia are dominant populations in the GBM with high heterogeneity and functional plasticity (34). Microglia cells (35) and microglia-like nerve-associated macrophages (17–20) have been described to display neurogenic properties. To further study whether the identified TAM “*neural growth”* signature was also expressed by microglia cells, we conducted a subset analysis on TAM and microglia clusters within our GBM scRNA sequencing dataset. This analysis resulted in the identification of TGFBI^+^/TMEM119^-^ TAM clusters (sub-clusters 7, 8, and 11) and remaining TGFBI^-^/TMEM119^+^ microglia clusters (sub-clusters 0-6, 9, 10; Fig. 1F, G). Clusters 1 and 10 were entirely composed of cells derived from healthy brain tissue (Fig. S1E) and expressed homeostatic microglia gene signature (Fig. 1H, I). The other microglia clusters were mostly derived from GBM samples and were divided into two distinct subclusters according to APOE expression (36): the Disease-Associated-Microglia (DAM, clusters 0 and 2) and the Glioblastoma-Associated-Microglia (GBM (34)-MG, clusters 3, 4, 5, 6, and 9) (Fig. 1H). GO analysis, visualized in the rounded GO plot (Fig. 1I), revealed that TAM cluster highly expressed, compared to all microglia clusters (e.g., homeostatic, DAM, and GBM-associated microglia), not only expected categories such as *response to hypoxia* and *angiogenesis* (5, 6), but also GOs related to *neural/axon development*, *myelination*, *tissue remodeling*, *injury response*, and *neuron/axon regeneration* (Fig. 1I). In line with this result, the expression of the “*neural growth”* gene list was highly upregulated in TAM compared to microglia clusters (Fig. 1J). We further verified the expression of “*neural growth*” signature in TAM in a publicly available bulk RNAseq database of mouse GBM comprising Mo, healthy microglia, GBM-associated microglia, and TAM populations (GSE86573 (33)). Consistently with our findings, in this dataset TAM upregulated the “*neural growth*” gene list compared to Mo (p=0.0089, Fig. 1K), healthy microglia (p<0.001, Fig. 1K), and GBM-associated microglia (p=0.029; Fig. 1K). These results highlighted a TAM-specific overexpression of the neurogenic signature compared to the healthy and disease-associated microglia.

Overall, we found that the “*neural growth*” signature is consistently retained by TAM extracted from eight different tumor datasets, both from human and mouse samples.

### TAM enhance tumor innervation through Secreted Phosphoprotein 1 (Spp1)

To functionally assess TAM “*neural growth*” potential, we studied the effect of of TAM *in vivo* on tumor innervation. To this aim, we utilized a sarcoma tumor model where TAM constitute the predominant immune cell population and have been demonstrated to play a pivotal role in tumor progression (37–39). Notably, scRNA analysis of mouse sarcoma dataset ((39); GSE201615, GSE201618, Fig. 2A) confirmed the presence of a TAM cluster, which was characterized by the upregulation of the “*neural growth”* gene signature (Fig. 2A). Adoptive transfer of macrophage in sarcoma model has been shown to promote tumor progression (39). To allow TAM detection, we *in vitro-*generated TAM-like macrophages from bone marrow hematopoietic progenitors constitutively expressing tdTomato (TAM*^tdTomato^*) from the B6.Cg-*Gt(ROSA)26Sor^tm9(CAG-tdTomato)Hze^*/J mouse line. To obtained TAM phenotype by exposing macrophages to a sarcoma microenvironment (see Material and Methods). In this condition, macrophages acquire pro-tumorigenic properties (39). We extensively characterized and verified the molecular and functional TAM properties of *in vitro* generated mouse TAM (see Extended results 1 and Fig. S2). We transplanted (day 0) MN/MCA1 sarcoma cells within skeletal muscles of a recipient mouse (40) coinjected or not with 5 x 10^5^ TAM*^tdTomato^*, and at day 14 TAM-treated mice received a second injection of 5 x 10^5^ TAM*^tdTomato^* (Fig. 2B). TAM*^tdTomato^* transplanted into the sarcoma are continuously exposed *in vivo* to tumor-derived stimuli, enabling their integration into the tumor microenvironment and sustaining their pro-tumoral functionality. Three weeks after the tumor graft (day 21), we analyzed the tumor area and the presence of nerves by immunofluorescence detection of the axon neurofilament-200 marker (NF200)(41). In line with the common pattern shared by nerve and blood vessel wiring (42), we observed NF200^+^ axons lining CD31^+^ vessels (Fig. S3A). As expected for tumor with so high proliferative rate (40), we did not observe any difference in the tumor area (Fig. S3B). TAM*^tdTomato^* transplantation determined a 2-fold increase of nerve number (p=0.046; Fig. 2C-E) and NF200^+^ axons (p=0.024; Fig. 2F, G), suggesting a role of TAM in promoting tumor innervation.

**Legend to Figure 2:**
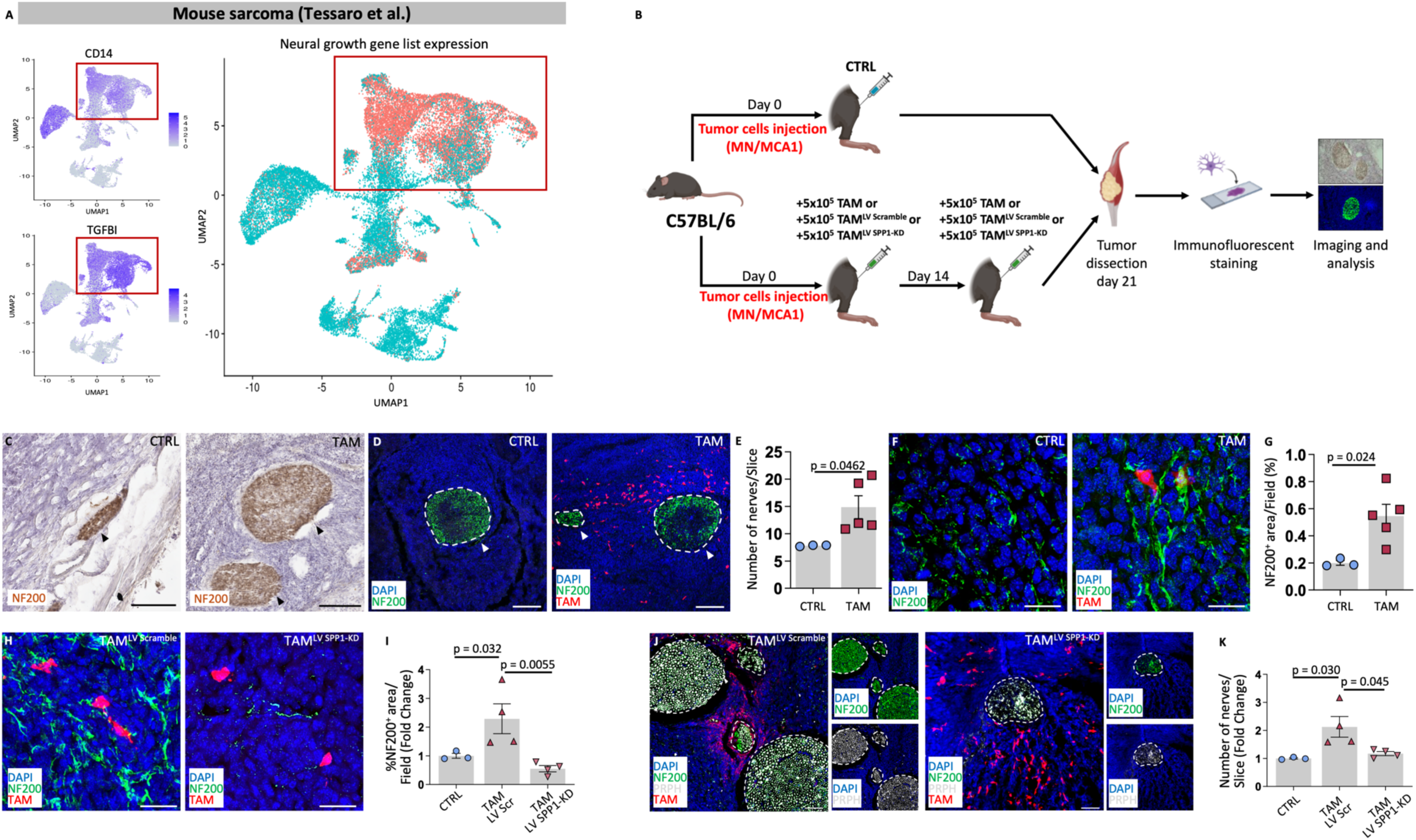
TAM promote tumor innervation in mouse sarcoma through spp1. **A)** (Left) UMAP plot showing CD14 and TGFBI expression (red square) in the mouse sarcoma scRNA dataset (blue cells present the expression of the relative marker); (right) AUCell analysis of the “neural growth” gene list (pink cells present higher expression of the genes included in the list) in the mouse sarcoma scRNA dataset showing its expression is increased in the CD14^+^/TGFB^+-^ TAM cluster (red square). **B)** Schematic representation of the experimental workflow. (day 0) C57Bl6 mice received MN/MCA1 cells alone or dispersed with TAM, TAM^LV^ ^Scramble^, or TAM^LV^ ^Spp1-KD^. (day 14) Treated animals received a second injection of respectively TAM, TAM^LV^ ^Scramble^ and TAM^LV^ ^Spp1-KD^. (day 21) Experimental animals were sacrificed and tumors dissected for further analysis. **C)** Immunohistochemical transversal sections of sarcoma tumor from control (CTRL) and TAM-treated mice stained with NF200 (brown) and hematoxylin (nuclei, purple). Black arrows indicate nerves within the tumor. Scale bar=150μm. **D)** Confocal immunofluorescence maximum intensity projections of z-stack transversal sections of sarcoma tumor from CTRL and TAM-treated mice stained with NF200 (green) and DAPI (blue). TAM are shown in red. White arrows indicate nerves within the tumor. Scale bar=200μm. **E)** Mean number of nerves per transversal section of sarcoma tumor in CTRL and TAM-treated mice, showing increased number of nerves in TAM-treated mice (CTRL: n=3; TAM: n=5). **F)** Confocal immunofluorescence maximum intensity projections of z-stack transversal sections of sarcoma tumor from CTRL and TAM-treated mice at 21 days stained with NF200 (green) and DAPI (blue). TAMs are shown in red. Scale bar=20µm. **G)** Percentage of the NF200^+^ area normalized for the total area of the field considered in transversal sections of sarcoma tumor from CTRL and TAM-treated mice, showing increased NF200 expression in TAM-treated mice (CTRL: n=3; TAM: n=5). **H)** Confocal immunofluorescence maximum intensity projections of z-stack transversal sections of sarcoma tumor from TAM^LV^ ^Scramble^- and TAM^LV^ ^Spp1-KD^ -treated mice stained with NF200 (green) and DAPI (blue). TAMs are shown in red. Scale bar=20µm. **I)** Percentage of the NF200^+^ area normalized for the total area of the field considered in transversal sections of sarcoma tumor from TAM^LV^ ^Scramble^- and TAM^LV^ ^Spp1-KD^ -treated mice, showing increased NF200 expression in TAM-treated mice calculated as fold change respect to CTRL, showing increased NF200 expression in TAM^LV^ ^Scramble^-treated mice (CTRL: n=3; +TAM^LV^ ^Scramble^ and +TAM^LV^ ^Spp1-KD^: n=4/condition). **J)** Confocal immunofluorescence maximum intensity projections of z-stack transversal sections of sarcoma tumor from CTRL, TAM^LV^ ^Scramble^- and TAM^LV^ ^Spp1-KD^ -treated mice stained with NF200 (green), peripherin (PRPH, white), and DAPI (blue). TAM are shown in red. Scale bar=200μm. **K)** Mean number of nerves per transversal section of sarcoma tumor in CTRL, TAM^LV^ ^Scramble^- and TAM^LV^ ^Spp1-KD^ -treated mice, calculated as fold change respect to CTRL, showing increased number of nerves in TAM^LV^ ^Scramble^ -treated mice (CTRL: n=3; TAM^LV^ ^Scramble^ and TAM^LV^ ^Spp1-KD^: n=4/condition). Data are expressed as mean±SEM. Statistical differences were evaluated by unpaired Student t-test or 1-way ANOVA.

To understand the molecular mediators of TAM neurogenic effect, we first performed a bioinformatic analysis and identified in both the *in vivo* humans and mouse datasets, the genes with large fold changes that were also strongly statistically significant; among these, we focused on the genes included in the “*neural growth*” gene list and selected secreted phosphoprotein 1 (SPP1) as the consistently highly upregulated gene (Fig. S3C, D). SPP1 was already described to play neurotrophic functions (29, 35, 43–47). We confirmed the expression and the secretion of SPP1 by *in vitro*-generated TAM (Fig. S3E). To validate the role of SPP1 as a mediator of TAM neurogenic effect, we first tested the effect of SPP1 inhibition in TAM*^tdTomato^ in vivo* on tumor innervation. We obtained SPP1 knockdown TAM (TAM^LV SPP1-KD^) by transducing TAM with a lentiviral vector expressing short hairpin against SPP1 (LVshSPP1:GFP), which strongly inhibited SPP1 expression (p=0.0006; Fig. S3F). Then, we tested TAM^LV SPP1-KD^ effect on tumor innervation in the sarcoma mouse model. We administered 5 x 10^5^ TAM^LV SPP1-KD^ or scramble transduced TAM (TAM^LV Scramble^) at day 0 and day 14 after tumor cell transplantation (Fig. 2B).

While we confirmed the TAM effect on tumor nerve growth assessed as expression of NF200^+^ axons with the administration of TAM^LV Scramble^ (CTRL vs TAM^LV Scramble^: p=0.032), TAM^LV SPP1-KD^ were not able to increase NF200^+^ axons (CTRL vs TAM^LV SPP1-KD^: p=0.54; TAM^LV Scramble^ vs TAM^LV SPP1-KD^: p=0.0055; Fig 2H, I) and the number of NF200^+^ and Peripherin^+^ nerve fibers compared to control (CTRL vs TAM^LV SPP1-KD^: p=0.77; CTRL vs TAM^LV Scramble^: p=0.030; TAM^LV Scramble^ vs TAM^LV SPP1-KD^: p=0.045; Fig. 2J, K, Fig. S3G) suggesting the role of SPP1 in TAM tumor innervation effect.

Collectively, these data showed that TAM enhances tumor innervation and neural growth and identified SPP1 as one relevant TAM-neurogenic effector.

### TAM play a direct role in neurite outgrowth and axonal regrowth

We showed that TAM own a “*neural growth*” signature and enhanced neural growth in tumor. To assess whether TAM have a direct role in neuronal outgrowth, we initially confirmed the upregulation of the genes included in the “*neural growth*” gene list in our bulk RNA sequencing of mouse *in vitro*-generated TAM compared to *in vitro*-generated M2 macrophages (Fig. 3A). We then tested *in vitro* the TAM*^tdTomato^* and M2*^tdTomato^* capability to promote neural sprouting in both human motor neurons (iPSC-MNs) derived from human induced Pluripotent Stem Cells (iPSCs) (48), and mouse neurons derived either from mouse neural stem cells (mNSCs) and from dorsal root ganglia (mDRGs) (49, 50). In line with the data observed *in vivo* in the tumor model, after co-culturing with TAM*^tdTomato^*, iPSC-MNs approximately doubled their neurite outgrowth assessed by the expression of the neurite-specific marker β3-Tubulin (β3Tub) (CTRL vs TAM: p=0.0041, Fig. 3B, C). This effect was not observed when iPSC-MNs where co-cultured with M2*^tdTomato^* (CTRL vs M2*^tdTomato^*: p=0.9993, M2*^tdTomato^* vs TAM p=0.0039; Fig. 3B, C). Similarly, TAM*^tdTomato^* co-cultured with mNSC increased the expression of the neuronal marker β3Tub 2.4-fold compared to control and to M2*^tdTomato^* (CTRL vs TAM: p<0.0001; CTRL vs M2: p<0.0001; TAM vs M2 p<0.0001; Fig. 3D, E). TAM*^tdTomato^* promoted neuronal outgrowth also in the peripheric nervous system model of mDRG neurons (CTRL vs M2: p=0.8837 CTRL vs TAM: p=0.0107; TAM vs M2 p=0.0053 Fig. 3F, G). These data showed that TAM, but not M2, exhibit direct neuronal outgrowth properties.

**Legend to Figure 3:**
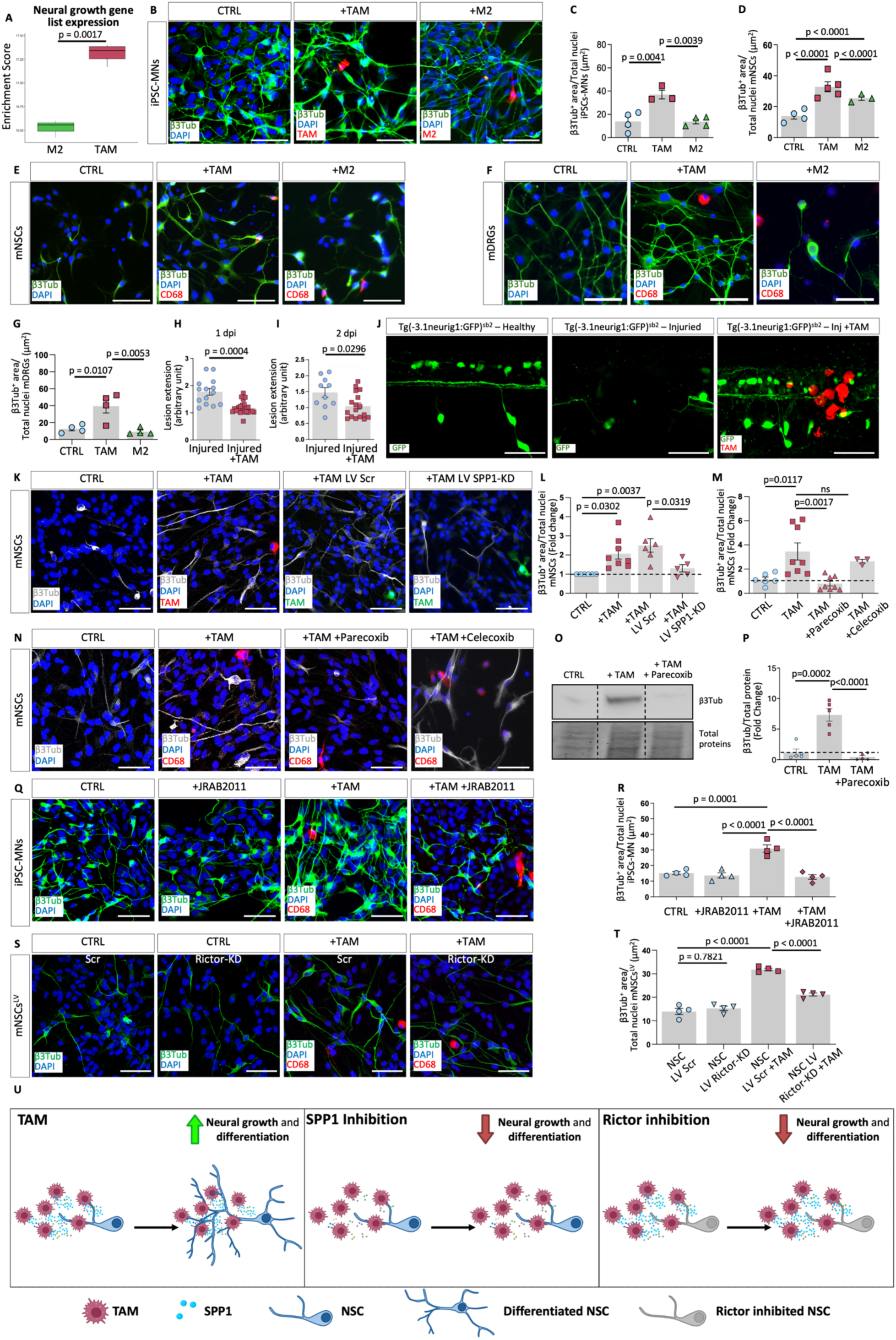
TAM enhance neuronal growth and axonal regeneration via SPP1. **A)** Gene set enrichment analyses of “neural growth” gene list in mouse M2 and TAM *in vitro*-generated (n=3/condition), showing enriched expression in TAM groups. **B)** Immunofluorescence images of human iPSC-derived MNs (CTRL) alone or cocultured with TAM*^tdTomato^* or ^M2^*^tdTomato^*(red), stained with the neurite marker β3Tub (green) and DAPI for nuclei (blue). Scale bar=50µm. **C)** Barplot showing increased β3Tub expression in iPSC-derived MNs treated with TAM (CTRL and M2: n=4/condition; TAM: n=3). **D)** Barplot showing increased β3Tub expression in mNSCs treated with TAM compared to all other conditions (CTRL: n=4; TAM: n=5; M2: n=3). **E, F)** Immunofluorescence images of (E) mNSCs (CTRL) or (F) DRG, cocultured with either TAM, or M2 co-cultures, stained with β3Tub (green) and DAPI (blue). CD68^+^ TAM and M2 macrophages are shown in red. Scale bar=50µm. **G)** Barplots showing increased β3Tub expression DRGs treated with TAM (CTRL: n=4; TAM: n=3). **H-J)** Barplots showing the quantification of the lesion extension in 2 days post fertilization (dpf) injured Tg(-3.1*neurog1*:GFP)*^sb2^* and injured TAM-treated Tg(-3.1*neurog1*:GFP)*^sb2^* zebrafish embryos at 1 day post injury (dpi) (H), and 2dpi (I), highlighting a smaller lesion in TAM-treated embryo (Injured: n=14; Injured +TAM: n=23 at 1dpi; Injured: n=10; Injured: +TAM: n=17 at 2 dpi). Representative immunofluorescence images of Tg(-3.1*neurog1*:GFP)*^sb2^* zebrafish embryos in healthy, injured, and injured TAM-treated condition in J. GFP-neurogenin is shown in green, and TAM are shown in red. Scale bar=100µm. **K)** Confocal immunofluorescence maximum intensity projections of z-stack images of mNSCs (CTRL), cocultured with either TAM, or with TAM^LV^ ^Scramble^, and TAM^LV^ ^Spp1-KD^ stained with β3Tub (white), and DAPI (blue). Transduced TAM are shown in green. Scale bar=50µm. **L)** Barplot showing the increased β3Tub expression (fold change respect to CTRL) in mNSCs treated with TAM^LV^ ^Scramble^, which was inhibited when TAM^LV^ ^Spp1-KD^ were used (CTRL and +TAM^LV^ ^Scramble^: n=6/condition; +TAM: n=8; +TAM^LV^ ^Spp1-KD^: n=5). **M)** Barplot showing the increased β3Tub expression (fold change respect to CTRL) in mNSCs treated with TAM, which was inhibited with Parecoxib, but not Celecoxib, administration (CTRL: n=6; +TAM and +TAM+Parecoxib: n=8/condition; +TAM+Celecoxib: n=3). **N)** Confocal immunofluorescence maximum intensity projections of z-stack images of mNSCs (CTRL), cocultured with TAM with or without either Parecoxib or Celecoxib, stained with β3Tub (white) and DAPI (blue). Scale bar=50μm. **O)** Representative western blot and quantification **(P)** showing the increase β3Tub expression in mNSCs (CTRL) when cocultured with TAM, and the inhibition of this effect following Parecoxib administration. Proteins are indicated by their specific molecular weights (β3Tub, 55kDa) (n=5/condition). **Q)** Confocal immunofluorescence maximum intensity projections of z-stack images of iPSC-derived MNs (CTRL), iPSC-derived MNs +JRAB2011, iPSC-derived MNs +TAM, and iPSC-derived MNs +TAM+JRAB2011 co-cultures, stained with β3Tub (green), CD68 (red), and DAPI (blue). Scale bar=50µm. **R)** Barplot showing the increase of β3Tub expression in iPSC-derived MNs cocultured with TAM, which was inhibited following JRAB2011 administration (n=4/condition). **S)** ^Scramble^ and mNSC^LV Rictor-KD^ co-cultured w/o TAM stained with β3Tub (green), CD68 (red), and DAPI (blue). Scale bar=50µm. **T)** Barplot showing the increase of β3Tub expression in mNSC^LV^ ^Scramble^ cocultured with TAM, which was inhibited with mNSC^LV^ ^Rictor-KD^ (n=4/condition). **U)** Simplified schematic representation of the experimental design of the *in vitro* experiments in Fig. 3. TAM induced neural growth and differentiation in NSC. The pharmacological (Parecoxib)/lentiviral inhibition of SPP1 in TAM (TAM^LV^ ^SPP1-KD^) inhibit TAM-induced neural growth and differentiation. The pharmacological (JRAB2011)/lentiviral inhibition of Rictor in NSC (NSC^LV^ ^Rictor-KD^) inhibit TAM-induced neural growth effetc. Data are expressed as mean±SEM. Statistical differences were evaluated by unpaired Student t-test or 1-way ANOVA.

To test whether TAM may promote axonal regrowth, we transplanted TAM*^tdTomato^* in a zebrafish model of axonal regeneration. We injected TAM*^tdTomato^* in Tg(-3.1*neurog1*:GFP)*^sb2^* zebrafish (51) embryos expressing the reporter protein GFP in neuronal cells, allowing spinal axonal fiber expression at the lesion site. At both 1 and 2 dpi, TAM*^tdTomato^* transplantation significantly reduced the extension of the neural damaged area compared to vehicle- (dPBS) treated zebrafish, suggesting a role of TAM in the promotion of the axonal regeneration (p=0.0004 and p=0.0296 respectively; Fig. 3H-J).

These data indicated that TAM have a direct role in neuronal outgrowth and axon regeneration both *in vitro* and *in vivo*.

We then further assessed if SPP1 was a relevant mediator in this TAM direct effect on neural outgrowth on mNSCs *in vitro*. We found that TAM^LV SPP1-KD^ were not able to increase β3Tub expression as well as TAM^LV Scramble^ (CTRL vs TAM: p=0.0302; CTRL vs TAM^LV Scramble^: p=0.0037; TAM^LV Scramble^ vs TAM^LV SPP1-KD^: p=0.0319; Fig 3K, L). We further confirmed the SPP1 role by using Parecoxib, a known pharmacological antagonist of the N4R4A transcription factor required for SPP1 expression (52). Parecoxib significantly reduced TAM SPP1 expression (p=0.0073; Fig. S3H). Spp1 pharmacological downregulation inhibited TAM-induced neurite outgrowth (CTRL vs TAM: p=0.0117; TAM vs TAM+Parecoxib: p=0.0017 Fig. 3M, N). Western blot analysis confirmed the effect of TAM on neuronal β3Tub upregulation, which was inhibited by Parecoxib (CTRL vs TAM: p=0.0002; TAM vs TAM+Parecoxib: p<0.0001; Fig. 3O, P). We further verified that the increased effect on neurite outgrowth was not observed when using Celecoxib, a COX2 inhibitor not affecting Spp1 expression (Fig 3M, N). Collectively, these data confirmed the role of Spp1 as required mediator of TAM induced neural growth.

In neurons, Spp1 signalling has been shown to promote neural maturation by activating Rictor signalling pathway (53). Therefore, we investigated whether the neurogenic response induced by TAM was regulated by Rictor signaling activation in neurons. To verify this hypothesis, we tested the effect of Rictor pharmacological inhibition by JRAB2011(54) on the TAM-induced neuronal differentiation of human iPSC-MNs. JRAB2011 treatment reduced the TAM-induced increase of β3Tub expression, indicating a role of Rictor in TAM effect on neural differentiation (TAM vs TAM^JRAB2011^: p<0.0001; Fig. 3Q, R). To exclude any potential direct effect of JRAB2011 administration on the cell culture, we assessed the β3Tub expression in human iPSC-MNs and found no difference between cells cultured with or without the inhibitor (Fig. 3Q, R). To further confirm the role of Rictor signaling in mediating TAM neurogenic effect, we downregulated Rictor expression in neurons derived from mNSC (NSC^LV Rictor-KD^) by lentiviral transduction, and we evaluated the β3Tub content following co-culture with/without the presence of TAM. Following 2 days of co-culture, we observed a reduced TAM-effect on β3Tub expression in NSC^LV Rictor-KD^ compared to NSC^LV Scramble^ (p<0.0001, Fig. 3S, T). No difference in β3Tub expression between NSC^LV Rictor-KD^ and NSC^LV Scramble^ was found, excluding any phenotypical variation in mNSC (p=0.782; Fig. 3S, T). Our results suggested that Rictor expression in neurons is required for TAM neurogenic effect.

Altogether these data indicated a direct role of TAM in promoting neural growth, confirmed Spp1 as relevant mediator of TAM, and Rictor as required signalling pathway for TAM effect in neurons (Fig. 3U).

### Adoptive transfer of TAM promotes neural regeneration in a model of severe CNS lesion

Adult mammalian CNS is characterized by a maladaptive wound repair (21). We exploited this newly identified TAM neurogenic property for adult CNS regeneration. To highlight the regenerative potential of TAM, we selected a severe complete-compressive-contusive spinal cord injury (scSCI) mouse model (41), which develops a permanent complete loss of motor functions and is considered the most predictive of the human condition (55, 56). Unlike the commonly used rodent models, which recover spontaneously and quickly after SCI (57), the scSCI model led to complete paralysis of the posterior legs without spontaneous functional recovery and severe impairment of the bladder functionality, which requires intense animal care (Fig. 4A-E; SMov. 1, 2) (41). In addition, the scSCI mouse model used in this study exhibited neurobiological pathology of the spinal cord, effectively mirroring the injury response observed in humans with complete SCI, a condition that remains incurable and profoundly affects the quality of life for patients (58). Following the primary insult, which vastly damages the neural tissue, secondary reactions develop subacute phase, cellular loss, inflammatory cell infiltration, tissue ischemia (59), and the damaged extracellular architecture of the tissue, promote the formation of large cysts, which isolate the injured parenchyma from the intact neural tissue by a fibro-glial scar which is not conducive to regeneration and repair (60, 61).

**Legend to Figure 4:**
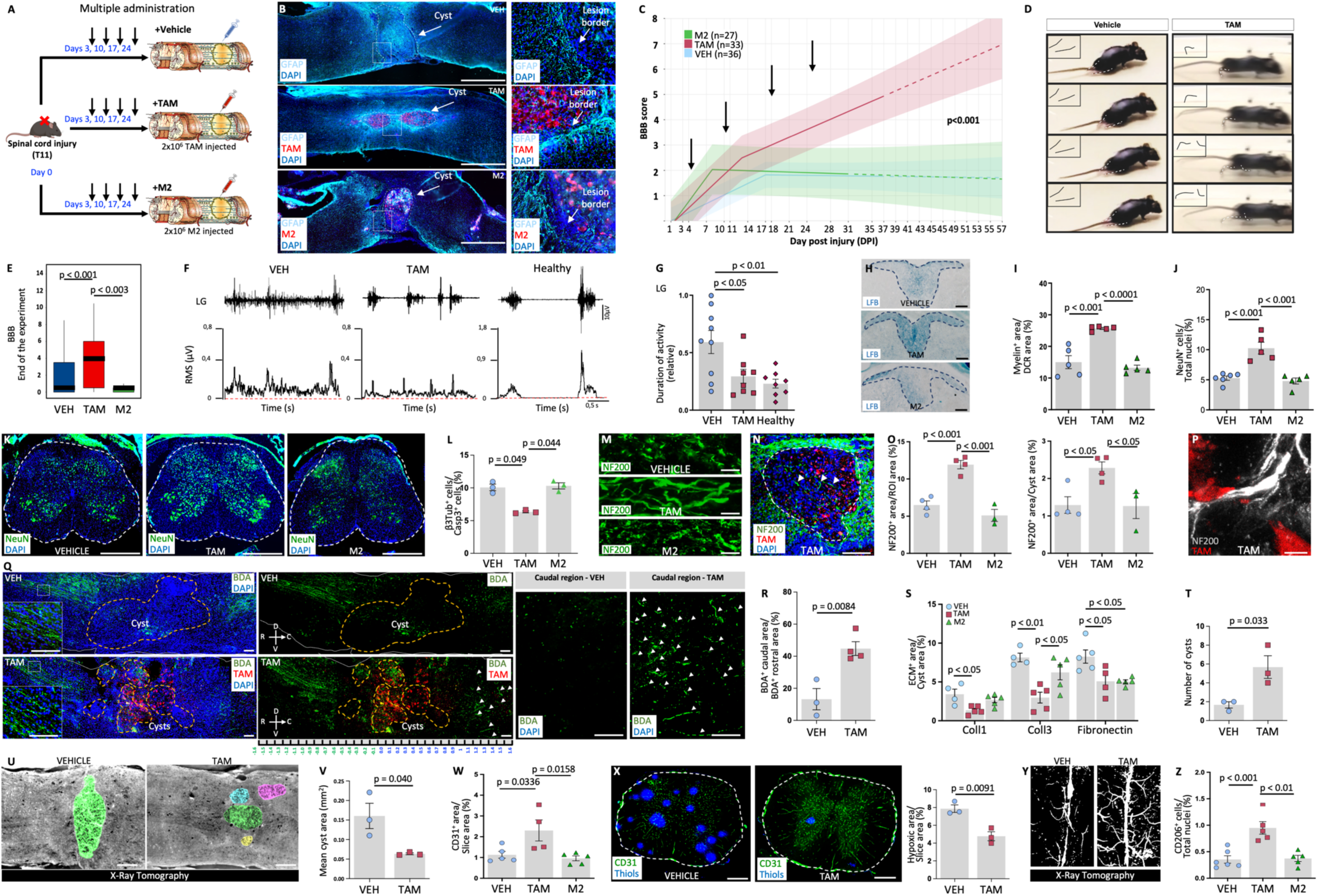
TAM transplantation promotes partial motor functional recovery and neural regeneration in severe spinal cord injury. **A)** Schematic representation of the workflow for the multiple administrations in severe contusive-compressive spinal cord injury (scSCI). **B)** Confocal immunofluorescence of longitudinal spinal cord sections of VEH, TAM-, and M2-treated scSCI stained with GFAP (cyan) and DAPI). Dashed white lines indicate the cyst. TAM and M2 are shown in red. **C)** BBB score analysis of mice motor function in VEH (light blue), TAM- (red), and M2-treated (green) scSCI mice estimated using a mixed-effect regression model. The three lines showed significantly different slopes indicating TAM effect in locomotor recovery of scSCI mice (VEH: n=36; TAM: n=33; M2: n=27; see supplementary information for additional statistical information). Black arrows indicate the days of injection/transplantation. Dashed lines represent the predicted response of the mice after the end of the experiment. **D)** Representative photograms of VEH and TAM-treated scSCI mice steps. **E)** Mean BBB score at the end of the experiment 31-37 dpi of VEH, TAM-, and M2-treated scSCI mice. TAM-treated scSCI mice showed higher BBB score compared to all other groups. **F)** Examples of *Lateral Gastrocnemius* (LG) electromyography (EMG) recordings expressed in µV (upper traces) and the related instantaneous (time constant: 0.04s) root mean square (RMS; time interval: 0.04s, time constant 0.01s) values (bottom graph) of VEH and TAM-treated scSCI mice and of a healthy mouse. scSCI animals present a distinctive EMG pattern characterized by lower RMS amplitudes and longer periods of activity compared to healthy mice. Dotted red lines indicate the selected threshold to distinguish LG-muscle activity. **G)** *Lateral Gastrocnemius* (LG) duration of activity relative to the total duration of recording in VEH and TAM-treated scSCI mice and in healthy mice. A value of 1.0 corresponds to a muscle constantly active throughout the recording session (VEH: n=9; TAM and healthy: n=8/condition). Adoptive transfer of TAM partially restored the duration of activity to a normal condition (Healthy). **H)** Magnifications of the dorsal column region (DCR) from brightfields image of transversal spinal cord sections of VEH, TAM-, and M2-treated scSCI mice, stained with myelin with the Luxol Fast Blue (LFB) protocol. The dotted blue line delineates the DCR. Scale bar=100µm. **I)** Percentage of the Myelin^+^ area normalized for the total DCR area. TAM-treated scSCI mice showing an increased myelin content compared to all the other groups (n=5/condition). **J)** Percentage of NeuN^+^ cells normalized for the total nuclei, TAM-treated SCI mice showed an increased NeuN^+^ cell content compared to all the other groups (VEH: n=6; TAM and M2: n=5). **K)** Immunofluorescence images of transversal spinal cord sections of VEH, TAM-, and M2-treated scSCI mice stained with NeuN (green), and DAPI (blue). The dotted white line delineates the section area. Scale bar=500µm. **L)** Graph showing that TAM treatment reduces the % of β3Tub^+^ cell death in iPSC-derived MNs compared to CTRL and M2. **M)** Magnification of longitudinal spinal cord sections of VEH, TAM-, and M2-treated scSCI mouse stained with NF200 (green) and DAPI (blue). Scale bar=50µm. **N)** Magnification from an immunofluorescent image of a longitudinal spinal cord section of a TAM-treated scSCI mouse stained with NF200 (green) and DAPI (blue) showing the lesioned area. The dotted white line delineates the cyst area. TAM-treated mice showed NF200 expression inside the cyst area (white arrows). Scale bar=100µm. **O)** (Left), Percentage of the NF200^+^ area normalized for the total area in the region of interest (ROI) in VEH, TAM-, and M2-treated scSCI mice. (Right) Percentage of the NF200^+^ area normalized for the cyst area in VEH, TAM-, and M2-treated scSCI mice. TAM-treated scSCI mice showed increased NF200 expression in both the lesioned parenchyma and in the cyst compared to VEH and M2-treated scSCI mice (VEH and TAM n=4; M2 n=3). **P)** Magnification of a longitudinal spinal cord section of a TAM-treated scSCI mouse stained with NF200 (white). TAM are in close contact with NF200^+^ axon fibers. Scale bar=5µm. **Q)** (Left) Immunofluorescent images of longitudinal spinal cord sections of VEH- and TAM-treated scSCI mice stained with Biotinylated Dextran Amine (BDA, green) and DAPI. Yellow dotted lines indicate the cyst border. White lines delineate the spinal cord parenchyma. TAM are shown in red. Scale bar=100µm. The insets represent magnifications of the rostral regions. Scale bar=100μm. (Middle) Immunofluorescent images of longitudinal spinal cord sections of VEH- and TAM-treated scSCI mice stained with BDA (green). Yellow dotted lines indicate the cyst border. White lines delineate the spinal cord parenchyma. TAM are shown in red. Scale bar=100µm. (Right) Magnification of the caudal region. White arrows indicate the BDA^+^ fibers. Scale bar=100µm. **R)** Percentage of BDA^+^ caudal area normalized for the BDA^+^ rostral area in VEH- and TAM-treated scSCI mice, showing higher BDA^+^ caudal area normalized for the BDA^+^ rostral area in TAM-treated compared to VEH-treated scSCI mice (n>3/condition). **S)** Percentage of relevant extracellular matrix (ECM) components (Coll1, Coll3, and Fibronectin) normalized for the total cyst area in VEH, TAM-, and M2-treated scSCI mice. In the cyst area, TAM-treated scSCI mice showed decreased Coll1 and Coll3 content compared to VEH and M2-treated mice, and reduced fibronectin expression compared to VEH (n>3/condition). **T)** TAM-treated scSCI mice showed an increased number of cysts compared to VEH-treated mice TAM-treated scSCI mice (n=3/condition). **U)** Representative images of longitudinal sections of VEH and TAM-treated scSCI mice obtained by X-ray tomography, VEH mice showed a cyst as a single structure, while TAM-treated mice showed cysts formed by smaller multilobes. Scale bar=200µm. **V)** TAM-treated scSCI mice show reduced mean cyst area compared to VEH-treated scSCI mice (n=3/condition). **W)** (Left) Percentage of CD31^+^ area normalized for the total section area in VEH, TAM- and M2-treated scSCI mice. TAM-treated scSCI mice showed higher CD31 content compared to both VEH- and M2-treated scSCI mice (VEH and M2: n=5; TAM: n=4). **X)** (Left) Immunofluorescence images of transversal spinal cord sections of VEH- and TAM-treated scSCI mice stained with reduced thiols marker (blue) and CD31 (green). The dotted white line delineates the section area. Scale bar=500µm. (Right) Percentage of hypoxic area normalized for the total section area in VEH- and TAM-treated scSCI mice, showing a reduced hypoxic area in TAM-treated scSCI mice compared to VEH-treated scSCI mice (n=3/condition). **Y)** Representative 3D rendering image of VEH- and TAM-treated spinal cord vessels obtained by X-ray computed tomography, showing increased TAM-treated mice vascular network compared to CTRL. **Z)** Percentage of CD206^+^ cells normalized for the total nuclei showing increased CD206^+^ cell content in TAM-treated scSCI mice compared to all the other groups. (VEH: n=6; TAM and M2: n=5). Data are expressed as mean±SEM. Statistical differences were evaluated by unpaired Student-t test and 1-way ANOVA.

#### TAM neurogenic property in scSCI: motor function

The comparison between TAM*^tdTomato^* and M2*^tdTomato^* transcriptomic profiles clearly highlighted that TAM enhanced the neuro-regenerative and wound healing potential compared to M2 (Fig. Fig. 3A, S2C). M2 macrophages have been reported to be effective in preclinical mild SCI models (57), though they had no beneficial effects in human clinical trials (62). M2 macrophages were never tested in scSCI animal model. We therefore tested both TAM and M2 neuro-regenerative properties in scSCI. We performed intraparenchymal transplantation of TAM*^tdTomato^*, M2*^tdTomato^* macrophages, or vehicle three days after the scSCI, corresponding to the later subacute phase (2– 4 days post-injury, dpi) (58, 61). TAM*^tdTomato^* and M2*^tdTomato^* transcriptomic analysis after 7 (t7) days from transplantation in the injured spinal cord showed that both the cell phenotypes were transient, suggesting that repeated administrations of cells could improve their regenerative efficacy (Fig. S4). Accordingly, we increase the dose from 1 (Fig. S5A-E) to 3 or 4 administrations (Fig. 4A). We performed multiple transplantations of 2×10^6^ TAM*^tdTomato^*, 2×10^6^ M2*^tdTomato^* or vehicle at 3, 10, 17 and 24 dpi (Fig. 4A). Considering animal wellbeing, the potential sufferance and damage associated to intra-parenchymal injections, we limited the dose escalation to a maximum of four cell transplants administered once a week and the end of the experiment to 37 days post injury (Fig. 4A). To assess the functional relevance of the treatments, we performed a double-blind evaluation of the animal motor function scored as BMS (see Supplementary Table S2a, b; Fig. S5F, G) (63) and as BBB adapted for mouse (64, 65), which resulted more sensitive to describe the recovery of stepping and coordination (see Material and Methods, Fig. 4C, E, Fig. S5E) (64, 65). Of note, following the commonly used SCI contusive mouse model (70 kdyn) reported in the literature, mice exhibited a mild impairment of the hindlimb paws corresponding to an extensive movement of the ankle joint (BBB score = 7, adapted for mouse), and showed spontaneous motor recovery (63–65). In this experiment, following scSCI (41), mice resulted in severe disability as shown by a BBB ≤ 0.5 at 1DPI, which remained stably ≤ 1.8 in all the mice for the entire experiment (VEH mean BBB at 37 DPI 1.8 ± 0.636; Fig. 4C-E and Fig. S5E), resulting in complete paralysis of the hindlimb paws (Fig. 4C-E; SMov. 1, 2). We evaluated the motor recovery of each animal at least every 3 days during the entire duration of the experiment by using the BBB score (65) (n=97 total animals tested). After scSCI, TAM-treated mice exhibited a progressive improvement of the hindlimb functionality compared to vehicle, reaching a significant partial motor recovery from day 21, with a mean BBB score at the end of the study (37 DPI) of 5.6 ± 1.09 (VEH vs TAM: p<0.001; Fig. 4E) corresponded to an extensive movement of at least one ankle (SMov. 1, 2). With this scSCI preclinical model, adoptive transfer of M2 was not effective and resulted in no improvement of motor disability. M2-treated mice maintained, as the vehicle-treated mice, a mean BBB score ≤ 2 over the entire experiment, never reaching the ability of a slight movement of the ankle (p=0.9 compared to vehicle; Fig. 4C-E, Fig. S5E). Accordingly, TAM-treated mice improved motor recovery also compared to M2-treated mice (p=0.003, Fig. 4E). The use of the linear mixed-effect regression model (66) with a single change-point on the whole longitudinal analysis spanning from 1 DPI to 37 DPI further highlighted the statistically significant differences between the slopes of VEH- and M2-treated groups in comparison to that of TAM-treated group (p<0.001) (Fig. 4C). Interestingly, while the average BBB score grew linearly in all groups before a change-point, after this change-point (13 dpi for TAM; Table S2a), the average BBB score continued to grow linearly only in the TAM group, with a slope of 0.102 (p < 0.001). At the end of the study TAM treatment resulted in an effect size (ratio of means) of 2.7 (95%CI 0.49-4.9, p=0.017) and 11.64 (95%CI=2.5-20.77, p=0.013) compared to vehicle- and M2-treated groups respectively (Fig. 4C; Table S2a). Additionally, the mixed-effects regression model predicts that the TAM treatment will increase the BBB score from 5.6 at 37 dpi to 6.98 at 57 dpi, 10 days after the experiment ends (dashed lines in Fig. 4C; BMS-based results that replicate the BBB-based ones are presented in Fig. S5F, G, and Table S2b). Spasticity, a symptom of the upper motor neuron impairment, is a common hallmark resulting from an injury, and it is a major cause of disability in individuals affected by a variety of CNS diseases and trauma (67). In our model, the severe injury of the spinal cord was associated with the onset of intrinsic tonic spasticity (41), evaluated through the analysis of the assisted movement of the ankle joint from 14 DPI onwards. In line with improved motor function, we observed a statistic reduction of the ankle joint spasticity following TAM treatment (Fig. S5H, I). The use of a mixed-effect linear regression model confirm that the three slopes were statistically different (Fig. S5I; Table S2c). To confirm the long-lasting effect of TAM*^tdTomato^* administration on motor neuron function, we performed *in vivo* electromyography (EMG) analysis of the foot extensor and flexor muscles during voluntary contractions in awake healthy mice, and in TAM- and vehicle-treated mice after 35 days from the injury. These muscles are antagonist to each other and are innervated by motor neurons whose cell body is caudal to the level of scSCI. As expected (68), severely injured animals displayed a distinctive EMG pattern, characterized by a lower amplitude and longer periods of activity compared to healthy mice (Fig. 4F, G and Fig. S5J, K). This EMG pattern was consistent with muscle paralysis and motor neuron hyper excitability. As a measure of motor neuron excitability, we quantified the amount of muscle activation relative to the total duration of recording, a ratio that is larger with hyper-excitable motor neurons (67). Adoptive transfer of TAM*^tdTomato^* restored the duration of activity of foot extensor muscles to normal in the entire population of recorded muscles, as revealed by the cumulative probability analysis (p<0.05; Fig. 4F, G and Fig. S5J). No differences were observed for foot flexor muscles (Fig. S5K). The recorded EMG activities were dependent on the voluntary activation of motor neurons, as no signal was detected following total transection of the spinal cord at thoracic 10 vertebra level (i.e., cranially to the scSCI level).

The overall assessments of the motor function of TAM-treated mice showed a statistically significant partial recovery compared to vehicle- and M2-treated mice.

#### TAM neurogenic property in scSCI: neural parenchyma

To investigate if the different therapeutic effects observed between TAM*^tdTomato^* and M2*^tdTomato^* could be attributed to any difference in their biodistribution, we quantified the distribution of transplanted cells by bioluminescence imaging of the emitted fluorescence exhibited by spinal cord and visceral organs of injured mice (Fig. S6). Immunohistochemistry confirmed a similar accumulation of transplanted cells in the lesioned and peri-lesioned spinal cord parenchyma between TAM and M2 (Fig. S6A-C). No tdTomato cells were found in all other visceral organs considered (heart, lung, pancreas, spleen, stomach, kidney, tests, bladder, skin lymph nodes, brain, liver and gut) indicating that they remain both localized at the injury site (Fig. S6D, E) and do not disseminate throughout the body.

To assess the impact of adoptive transfer of TAM*^tdTomato^* on spinal cord neural parenchyma repair, we quantified the abundance of myelin and spinal cord neurons at the site of injury. We used GFAP staining to delimitate the fibrotic-glial interface (41). For the spinal cord tissue analysis, we considered a perilesional region approximatively from +0.35 cm rostral to -0.35 cm caudal to the cyst (Fig. S5A). After scSCI, the lesion site developed a cyst surrounded by glial/fibrotic scar as shown by the GFAP staining (Fig. 4B). Compared to vehicle- and M2-treated animals, TAM-treated mice promoted myelination (1.7 fold: VEH vs TAM: p<0.001; TAM vs M2: p<0.0001; Fig. 4H, I) and doubled the number of all spinal neurons (VEH vs TAM: p<0.001; TAM vs M2: p<0.001; Fig. 4J, K) and specifically the ChAT-expressing spinal motor neurons (VEH vs TAM: p<0.001; TAM vs M2: p<0.0001; Fig. S7A, B). We validated the direct trophic and pro-survival effects of TAM by *in vitro* co-culturing TAM with iPSC-MNs in conditions of oxygen and glucose deprivation (OGD), which promote neuronal death. We found that TAM were able to decrease neuronal death following OGD up to 50% compared to control and M2 (VEH vs TAM: p=0.049; TAM vs M2: p=0.044 respectively; Fig. 4L; Fig. S7C). These results indicated that TAM promoted a direct effect on neuronal survival. Adoptive transfer of TAM*^tdTomato^* in scSCI also increased by 2-fold the expression of the NF200^+^ axons in the injured spinal cord parenchyma and those passing through the cyst compared to vehicle- and M2-treated mice (both p<0.05 and p<0.05 respectively; Fig. 4M-O), suggesting that TAM treatment improved axonal regrowth. Notably, we observed that some transplanted TAM*^tdTomato^* were in close contact to axonal fibers (Fig. 4P).

We then confirmed the axonal regrowth property of TAM, by labelling the descending corticospinal tract fibers with the anterograde axonal tracer BDA in the scSCI animal model (69). We quantified the presence of axons crossing the lesioned area (cyst). Notably, our scSCI model resulted in the formation of a large cyst which completely disrupts axonal connections and neural tissue. scSCI distinctly differs from spinal cord hemisection, the commonly used model to test anterograde axonal tracer BDA where half spinal cord is sharply cut. TAM treatment showed a 3.4-fold increase in BDA positive fibers in the spinal cord region caudal to the lesion compared to vehicle-treated mice (p=0.0084; Fig. 4Q, R), indicating a higher number of axonal fibers crossing and extending through the cyst.

Altogether these data showed the TAM improve motor function in scSCI, which is supported by a direct effect on neuronal cell survival and axonal regrowth.

### Adoptive transfer of TAM rewires the injured neural parenchyma microenvironment following scSCI

Efficient CNS repair requires not only neural survival and re-growth but also comprehensive restoration of the spinal cord parenchyma. This includes re-establishing vascular perfusion, resolving extracellular matrix (ECM) deposition, and mitigating chronic inflammation. Beyond their newly identified neurogenic activity, TAM exhibit wound-healing functions such as ECM remodeling, angiogenesis, hypoxia response, immune modulation, detoxification, stress response, and glucose metabolism. These capabilities likely support tissue regeneration and contribute to TAM therapeutic efficacy in spinal cord injury (scSCI). Comparative transcriptomic analysis with M2 macrophages, coupled with functional validation, confirmed that *in vitro*-generated TAM expressed the characteristics of *bona fide* TAM (Fig. S2). Consequently, we evaluated the TAM pro-regenerative effects on ECM remodeling, angiogenesis, and immune modulation within the injured spinal cord parenchyma.

#### Pro-regenerative effects of TAM: ECM remodeling

In injured spinal cord parenchyma, cyst size and composition are main aspects inhibiting spinal cord regeneration (58). Similarly, we found that relevant ECM components, such as collagen type-1 (Coll1), collagen type-3 (Coll3), and fibronectin (58–61), were decreased in the cyst of TAM-treated mice (p<0.05, p<0.01 and p<0.05 respectively; Fig. 4S; Fig. S7D). Compared to the cysts of M2-, the cysts of TAM-treated mice decreased the expression of Coll3 (p<0.05; Fig. 4S; Fig. S7D’’) and showed similar Coll1 and fibronectin content (Fig. 4S; Fig. S7D’, D’’’). We further analysed the volume and morphology of the cyst of vehicle- and TAM-treated mice using, for the first time in SCI, the *X-ray phase-contrast micro computed tomography*, which allows tissue analysis at micrometer resolution (70–73). To perform this analysis, we identified the cysts by a lighter edge which isolate the lesion scar from the outer parenchyma (Fig. 4T, U; Fig. S7E). The tomography considered for each spinal cord 735 slices covering a segment of 1 mm cantered to the lesion. Analysis of the cyst volume by quantifying GFAP stained tissues (Fig. S7F) shows no differences in the volumes between the different experimental groups. As expected for the scSCI model used, the cysts in vehicle-treated samples appeared as single compact structure, which completely separate the rostral and caudal (respect to lesion) spinal cord parenchyma in two discontinued portions (Fig. 4U, Fig. S7G, H, SMov. 3). Of note, whole tissue reconstruction obtained by X-ray phase-contrast micro computed tomography allowed us to detect that the cyst in TAM-treated spinal cords appeared fragmented in small and multiple lobes (p=0.033; Fig. 4T, U, Fig. S7G, H, SMov. 4) and did not separate the tracts of the spinal cord (Fig 4U). The mean area of the cysts, objectively measured by X-ray phase-contrast micro computed tomography, resulted 2.5-fold reduced in TAM-compared to vehicle-treated mice (p=0.040; Fig. 4U, V). These data suggested that TAM induce cyst remodeling possibly by using their enzymatic activity on the cyst ECM.

#### Pro-regenerative effects of TAM: angiogenesis

TAM are described to support angiogenesis trough VEGF in tumors (*42*). As shown in Fig. S2M-O we confirmed that adoptive transfer of TAM was able to promote angiogenesis in atransplanted in scSCI, by analyzing the vessel number and length and by measuring the extent of hypoxic regions at the spinal cord lesion site (Fig. 4W, X; Fig S7I-L). TAM-treated mice increased the expression of CD31^+^ vessel marker compared to vehicle- and M2-treated mice (p=0.0336 and p=0.0158 respectively; Fig. 4W, S7I). To understand whether the increase in vessels in TAM-treated mice correlates to a better tissue oxygenation in the injured spinal cord, we used an established protocol based on staining of oxidized thiols and quantified the hypoxic area (74). We observed that TAM reduced 1.7-fold the hypoxic area in scSCI (p=0.0091; Fig. 4X), indicating the increase of functional vasculature in the injured spinal cord. In line with the increased tissue oxygenation in TAM-treated mice, the mean maximum vessel length and mean branch number were higher in TAM-compared to vehicle-treated mice (p=0.0091 and p=0.013 respectively; Fig. S7J-L). Furthermore, Z-maximum projection of the entire spinal cord X-ray phase-contrast micro computed tomography showed an increased blood vasculature in TAM-treated spinal cords (Fig. 4Y, SMov. 5, 6). These data showed that TAM administration promote angiogenesis and spinal cord tissue oxygenation in scSCI.

#### Pro-regenerative effects of TAM: immune modulation

SCI microenvironment favours a persistent inflammatory setting associated with prolonged M1 polarization of infiltrating macrophages, which may contribute to spinal cord failure to regenerate and repair (62, 75). To assess the ability of TAM to modulate scSCI chronic inflammation, we quantified cells expressing the macrophages/microglia marker Iba1, and the M2 marker CD206 (75) in lesioned spinal cords (Fig. 4Y and Fig. S7M-O). Compared to vehicle- and M2-treated mice, TAM treatment exhibited an increased number of Iba1^+^ (p<0.01 and p<0.05 respectively; Fig S7M, N) and of CD206^+^ microglia/macrophages (p<0.001 and p<0.01 respectively; Fig. 4Z and Fig. S7O) in the spinal cord, suggesting a TAM immune-modulatory properties promoting M2 polarization, which is typically associated to wound healing process (76).

Collectively, these results indicated that TAM have a unique neuro-regenerative property and can modulate the scSCI-induced microenvironment by increasing ECM-remodeling, angiogenesis, and modulating chronic inflammation.

### Preclinical long-term (one year) assessment for tumorigenicity and toxicity confirms the safety of adoptive transfer of TAM in SCI

Given the multiple and robust mechanisms of action of TAM on the spinal cord, to highlight the translational potential of TAM for CNS regeneration, we evaluated the long-term toxicity and tumorigenic potential following single- and repeated-intraparenchymal administration of TAM in the spinal cord of mice that underwent laminectomy. One year after the transplantation, we performed a necroscopy, which includes comprehensive anatomical and histopathological investigations after single or repeated administration of vehicle or TAM. Furthermore, in all studied animals, we monitored weekly body weight and behavioral/clinical signs, and we did not observe weight or behavioral alterations, including locomotor functionality. Long term pre-clinical anatomical and histopathological investigations of adoptive transfer of TAM revealed no major histopathological findings in the visceral organs examined and, importantly none tumoral lesion of myeloid/macrophagic origin were observed (see Extended results 2 and Fig. S8A-C).

### TAM neural regenerative activity in CNS is mediated by the Rictor pathway

To better assess the molecular modification of the scSCI parenchyma following TAM administration, we analyzed the whole proteomic profile of treated and not treated lesioned spinal cord tissue. PCA showed a clear separation between vehicle- and TAM-treated proteome, indicating TAM-induced effects on the spinal cord injured parenchyma (Fig. 5A). This was also confirmed by the number of significant changes, with 227 proteins upregulated and 233 downregulated when TAM was compared to vehicle treatment (Fig. S9A) and supported by clustering of the log_2_ (Abundances) of differentially abundant proteins (DAPs) and enrichment in Gene Ontology terms, which revealed clear differences between TAM- and vehicle-treated groups (Fig. S9B, C). As M2 treatment had no therapeutic effect on motor recovery, we also compared the upregulated protein expression of TAM- and M2-treated spinal cords to vehicle (Fig. S9D). TAM-treated spinal cord tissue showed more upregulated proteins related to neurons, synapses, and myelin than M2-treated spinal cord tissue (p=0.038, p=0.0015 and p=0.0038, respectively; and 95 downregulated proteins (Fig. 5C), with clear differences in the log_2_(Abundances) (Fig. S9E). The enrichment analysis on identified DAPs and the enriched Pathway Analysis (IPA) terms obtained from the spinal cord proteomic analysis revealed clear differences between TAM- and M2-treated groups (Fig. 5D, E). We found a significant enrichment of 29 core pathways and 9 upstream regulators (p≤0.05). Among the significantly enriched IPA terms, we found *EIF2 signaling* and *Integrin signaling* to be decreased in TAM-compared to M2-treated mice. Furthermore, *MAPK signaling* was shown to be down-regulated in this comparison, while *PTEN signaling* and *Acute phase response signaling* were up-regulated. Analysis of upstream regulators associated with the DAPs revealed that, compared to both M2- and vehicle-treated spinal cords, TAM-treated spinal cords showed an up-regulation of Rictor (Fig. 5D, E), which is part of the mTORC2 complex (77). Rictor overexpression in SCI has been shown to promote SCI recovery (78). The proteomic results identifying Rictor as neural tissue effector of TAM action, were in line with the data obtained with *in vitro* neuronal cells, where Rictor inhibition damper TAM neurogenic effect (Fig. 3Q-T). We further investigated whether inhibition of the mTORC2 pathway *in vivo* could reduce the therapeutic effect of TAM on scSCI motor recovery. To induce the knockdown of Rictor, we injected a recombinant AAV9 viral vector encoding GFP and a short hairpin against the mouse Rictor sequence in the lesioned spinal cords at 3, 10, and 17 dpi. The GFP immunostaining confirmed that AAV9 was efficiently distributed in the injured spinal cord parenchyma at 24 dpi (Fig. 5F). AAV9 transduction significantly reduced the amount of Rictor (p=0.012; Fig. 5G, H) and decreased its downstream Ras homologous (Rho) protein family mediators (77) in the tissue homogenates of TAM-treated spinal cords (RhoA/B/C: p=0.0006; RhoQ: p=0.045; Fig. 5I-K). Of note, in TAM-treated Rictor^KD^ mice the locomotor recovery was significantly reduced (at 24 dpi: p=0.022; Fig. 5L) further supporting the role of Rictor in TAM-mediated effects. Collectively, these data were consistent with the role of Rictor and the mTORC2 pathway in TAM-induced therapeutic effect for SCI.

**Legend to Figure 5:**
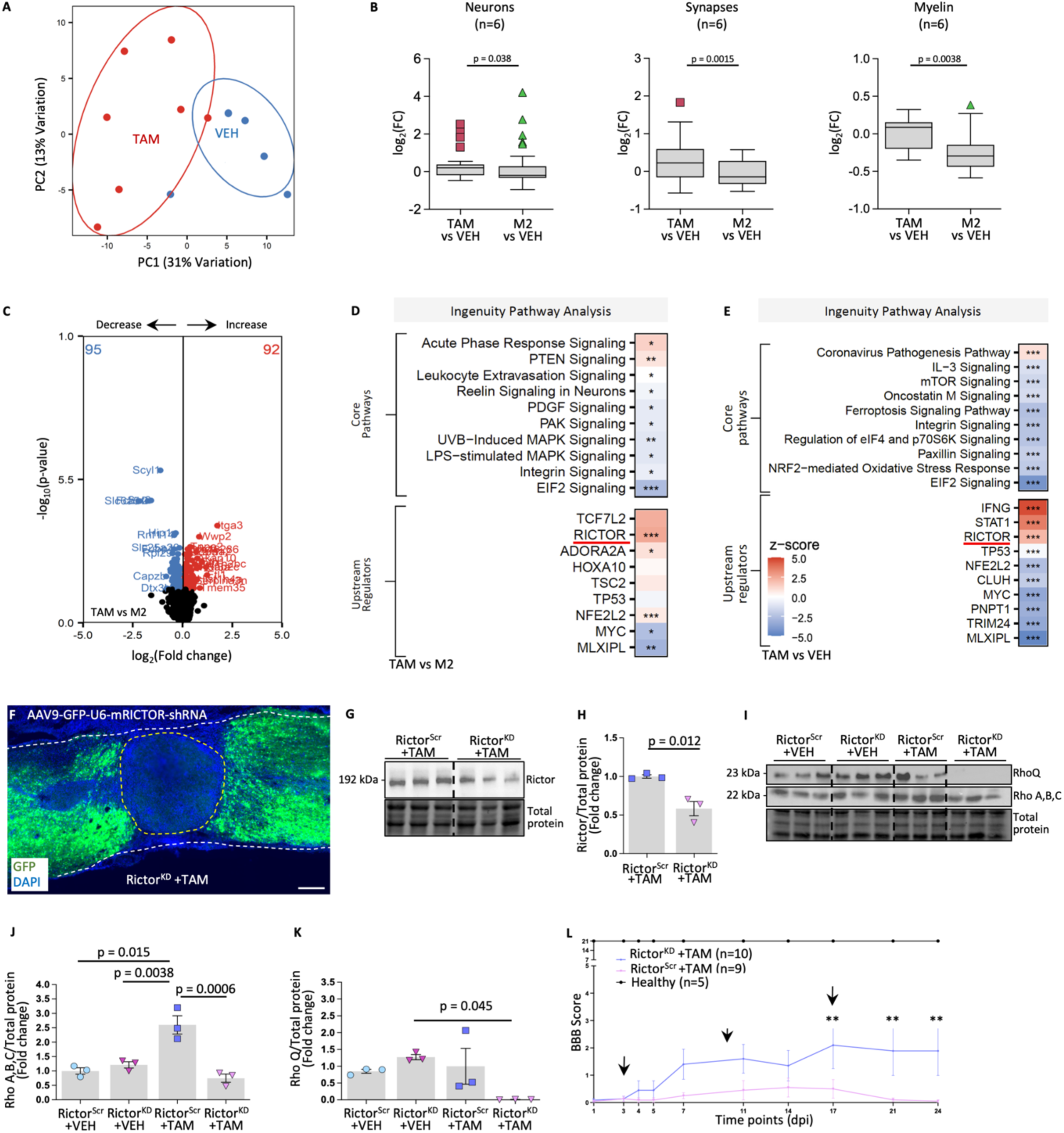
Protemic analysis and functional validation of Rictor mediated TAM therapeutic effect in scSCI. **A)** PCA of protein samples from VEH- and TAM-treated scSCI spinal cords (VEH: n=5; TAM: n=7). Each dot represents a sample. TAM and VEH groups are well separated, indicating a distinct proteomic profile. **B)** Boxplot showing that TAM-treated scSCI mice have a higher content of proteins related to neurons, synapses, and myelin expressed then M2. Protein content is expressed as log2(FC) of TAM/VEH and M2/VEH. **C)** Volcano plot showing differential protein abundance in the protein extracts from TAM- and M2-treated scSCI spinal cords. Proteins are separated according to their log2-transformed intensity fold-change (TAM/VEH; x-axis) and their -log10-transformed p-value (y-axis), (TAM: n=7; M2: n=7). Protein extracts from TAM-treated scSCI spinal cords showed 92 up-regulated proteins and 95 down-regulated proteins compared to protein extracts from M2-treated scSCI spinal cords. **D)** Ingenuity pathway analysis of more significant core pathways and upstream regulators in the TAM/M2 comparison. The red line shows RICTOR as one of the upstream regulators for the group differences. **E)** Ingenuity pathway analysis of core pathways and upstream regulators in the TAM/VEH comparison. The red line shows RICTOR as one of the upstream regulators for the group differences. **F)** Immunofluorescence image of a longitudinal spinal cord section from a Rictor^KD^ scSCI mouse stained with GFP (green) and DAPI (blue). AAV9 viral vector expressing GFP is efficiently distributed in the spinal cord parenchyma close to the lesion site. Dotted white lines delineate the spinal cord, the yellow one delineates the cyst area. Scale bar=200µm. **G)** Representative Rictor immunoblots of protein extracts from Rictor^Scr^ and Rictor^KD^ scSCI spinal cords treated with TAM. **H)** Rictor protein expression was decreased in TAM-treated Rictor^KD^ scSCI mice compared to TAM-treated Rictor^Scr^ scSCI mice (n=3/condition). **I)** Representative Rho A, B, C, and Q immunoblots of protein extracts from VEH-treated Rictor^Scr^ and Rictor^KD^ scSCI mice, and TAM-treated Rictor^Scr^ and Rictor^KD^ scSCI mice. **J)** Rho A, B, and C protein expression is increased only in TAM-treated Rictor^Scr^ scSCI mice (n=3/condition). **K)** Rho Q protein expression is increased only in TAM-treated Rictor^Scr^ scSCI mice (n=3/condition). **L)** Line plot of the mean BBB score of TAM-treated Rictor^Scr^ and Rictor^KD^ scSCI mice showing increased locomotor recovery only in TAM-treated Rictor^Scr^ scSCI mice compared to Rictor^KD^ scSCI mice (Rictor^Scr^+TAM: n=9, Rictor^KD^+TAM: n=10). Data are expressed as mean±SEM. Statistical differences were evaluated by unpaired Student t-test, 1-way ANOVA, and 2-way ANOVA with repeated measurements.

### Human TAM show neural growth functional phenotype

Our single cell transcriptomic data on multiple tumor samples showed that human and mouse TAM display a unique “*neural growth*” signature (Fig. 1D) and that mouse TAM have functional neurogenic properties, enhance tumour innervation and exhibit therapeutic effect in CNS lesions. Therefore, we assessed the expression of the TAM “*neural growth*” signature and the functional neurogenic properties also on *in vitro*-generated TAM obtained from human blood monocytes (see Materials and Methods). We confirmed that *in vitro*-generated human TAM (hTAM) expressed TAM molecular profile and function (see Extended results 3 and Fig. 6A, B, and Fig. S10). In line with these results, hTAM significantly upregulated also the newly identified “*neural growth*” signature compared to *in vitro*-generated human M2 macrophages (hTAM vs M2: p=0.0015; Fig. 6C). We further tested hTAM potential to enhance neural growth using human neuronal cells. We co-cultured hTAM with iPSC-MNs or SH-SY5Y-differentiated neuronal cells. We found hTAM located in close contact with neurons (Fig. 6D). Consistent with the data obtained with murine TAM, the addition of hTAM increased β3Tub expression in iPSC-MNs (p=0.0017; Fig. 6E, F). The increase in neurite outgrowth was not present when iPSC-MNs were co-culture with hM2 macrophages (CTRL vs hM2: p=0.8642; hTAM vs hM2: p=0.0119; Fig. 6E, F). We further assessed if the TAM effector Spp1 had a direct role on neural outgrowth also in hTAM. We first assessed its expression by immunofluorescence (Fig. 6G), FACS (Fig.6H, I), and RT-PCR analysis (Fig. 6J). Pharmacological Spp1 inhibition with Parecoxib (Fig. 6K) significantly reduced hTAM-induced neurite outgrowth in both iPSC-MNs (p=0.0088; Fig. 6L, M) and in SH-SY5Y-differentiated neuronal cells (p= 0.0009; Fig. 6N, R). This effect was not observed when using Celecoxib, a COX2 inhibitor not affecting Spp1 expression (Fig. 6L-N, R). We confirmed the role of SPP1 for the hTAM neural growth effect by down-regulating SPP1 expression using specific siRNA (Fig. 6K). While hTAM^siRNAScramble^ induce β3Tub expression of 2.1-fold (p=0.0006 compared to control, Fig. 6O, S), hTAM^siRNASpp1-KD^ were not able to increase β3Tub expression (p=0.557, hTAM^siRNASpp1-KD^ vs control; hTAM^siRNAScramble^vs hTAM^siRNASpp1-KD^; p=0.0233; Fig. 6O, S). Rictor/mTORC2 pathway was shown to be relevant mediator in neuronal cells and spinal cord tissue of the TAM neurogenic effect (Fig. 3Q-U, 5C-L). We further validated the involvement of Rictor/mTORC2 in hTAM neural growth effect (Fig. 6P, Q), and found that pharmacological inhibition of Rictor activity (JRAB2011), inhibited hTAM effect on neurite outgrowth in both SH-SY5Y-differentiated neuronal cells (p=0.0025; Fig. 6P, T), and in iPSC-MNs (p=0.0018; Fig. 6Q, U).

**Legend to Figure 6:**
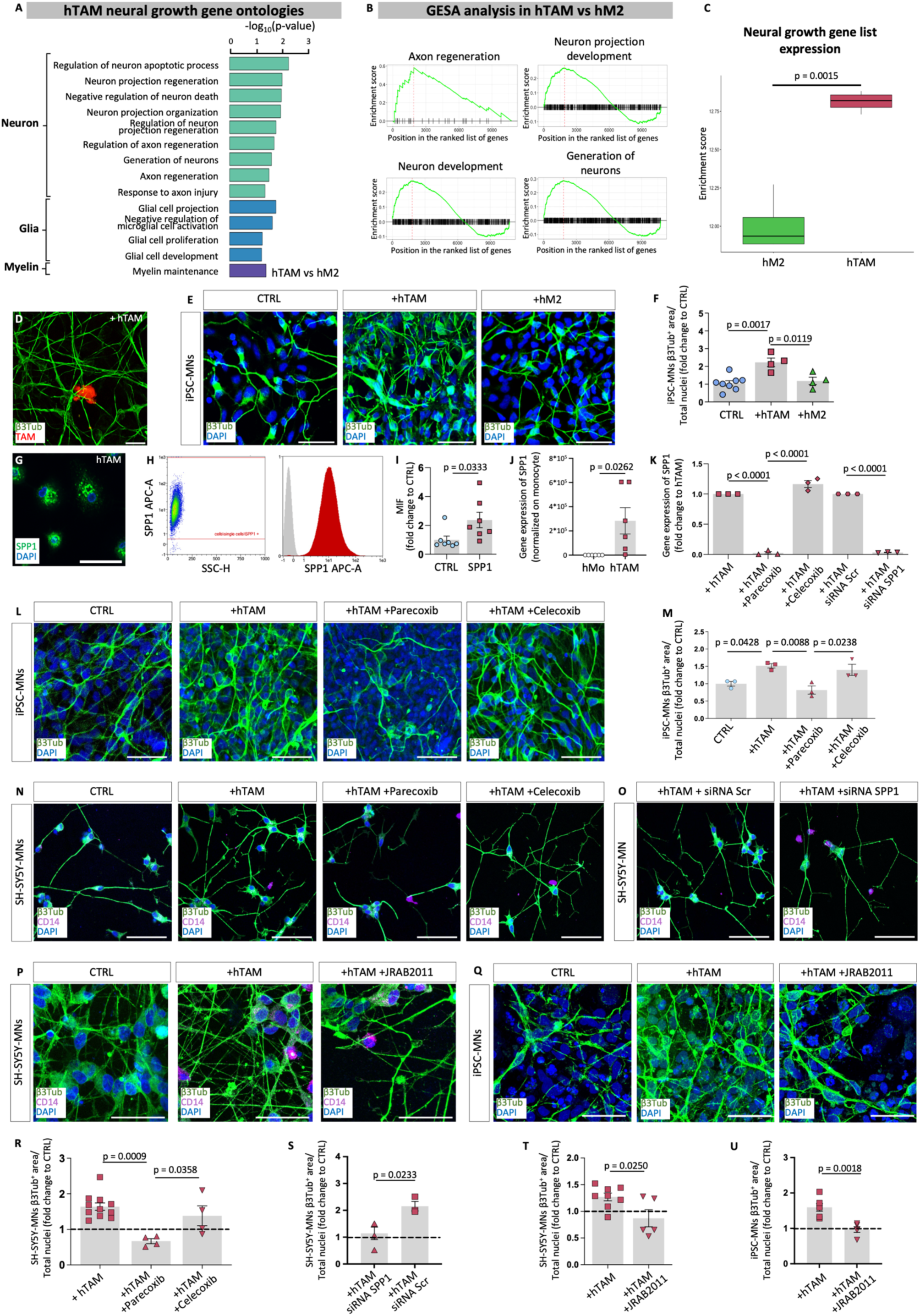
Human TAM display neural regenerative properties. **A)** Barplots showing enriched upregulated gene ontologies neural growth-related in hTAM compared to hM2. Barplot length is the -log10(p-value) returned from the enrichment analysis. **B)** Gene Enrichment Set Analysis (GESA) on neuro-related gene ontologies of *in vitro*-generated human TAM (hTAM) and M2 (hM2), showing enrichment of the genes in TAM compared to M2. **C)** Single sample gene set enrichment analysis of the “neural growth” gene list in hM2 and hTAM, showing enriched expression in hTAM group (n=4/condition). **D)** Magnification of human iPSC-derived MNs co-cultured with human TAM stained with β3Tub (green) and CD14^+^ (red). Scale bar=10µm. **E)** Immunofluorescence images of human iPSC-derived MNs (CTRL), human iPSC-derived MNs +hTAM, human iPSC-derived MNs +hM2 co-cultures, stained with β3Tub (green) and DAPI (blue). Scale bar=50µm. **F)** Barplot showing increse β3Tub^+^ area (%) following hTAM co-culture (CTRL: n=8; hTAM: n=4; hM2: n=4). **G)** Immunofluorescence images of hTAM stained with SPP1 (green) and DAPI (blue). Scale bar=10µm. **H)** Representative hTAM flow cytometry analysis showing the expression of SPP1 (red), unstained negative control (gray). **I)** Barplot showing the fold-change of the mean fluorescence intensity (MIF) of SPP1 compared to an unstained control sample, indicating TAM-specific expression of SPP1 (n=7/condition). **J)** Barplots showing the fold-change of the relative gene expression of SPP1of hTAMs compared to monocytes (n=6/condition). **K)** Gene expression of SPP1 in hTAM, hTAM+Parecoxib, hTAM+Celecoxib, hTAM siRNA^Scramble^ and hTAM siRNA^SPP1^ expressed as fold-change respect to hTAM, showing the efficacy of the SPP1 inhibition both via the pharmacological and the lentiviral route (n=3/condition). **L)** Immunofluorescence images of β3Tub neurites (green) of iPSC-derived MNs (CTRL), cocultured with hTAM alone or with parecoxib or celecoxib. Scale bar = 10µm. **M)** Barplot showing inhibition of hTAM-induced increase β3Tub expression of iPSC-derived MNs following Parecoxib. Celecoxib was not impairing hTAM effect (n=3/condition). **N-P)** Series of immunofluorescence images of SH-SY5Y-MN neurites (β3Tub, green), cocultured with hTAM (CD14, magenta) alone or in different condition: **N)** with Parecoxib or Celecoxib Scale bar = 10µm; **O)** with hTAM siRNA^Scramble^ or hTAM siRNA^SPP1^, Scale bar = 10µm; **P)** with JRAB2011. Scale bar = 50µm; **Q)** Immunofluorescence images of β3Tub neurites (green) of iPSC-derived MNs (CTRL), co-cultured with hTAM alone or with JRAB2011. Scale bar=50µm. **R)** Barplot showing inhibition of hTAM-induced increase β3Tub expression of SH-SY5Y-MN following Parecoxib. Celecoxib was not impairing hTAM effect. (hTAM: n=11; hTAM +Parecoxib: n=4; hTAM +Celecoxib: n=4) **S)** Barplot showing reduced SH-SY5Y-MN β3Tub expression following hTAM siRNA^SPP1^ co-culture compared to hTAM siRNA^scramble^. (hTAM siRNA^SPP1^ n=4; hTAM siRNA^scramble^ n=3) **T-U)** Barplots showing inhibition of hTAM-induced increase β3Tub expression (fold change) of SH-SY5Y-MNs (hTAM n=8; hTAM +JRAB2011 n=5) **(T)** or iPSC-derived MN (hTAM n=6; hTAM +JRAB2011 n=5) **(U)** following JRAB2011 administration. Data are expressed as mean ± SEM. Statistical differences were evaluated using the Student t-test and 1-way ANOVA.

Altogether these findings were in line with the finding obtained with murine TAM, indicating that hTAM are endowed with a neuro-regenerative phenotype. In addition to features related to wound healing, angiogenesis, ECM remodeling, and immune modulation (see Extended results 3 and Fig. S10), hTAM enhanced neural growth. Consistently with results on murine TAM, we confirmed Spp1 as a relevant neural growth effector also for hTAM, and Rictor as mediator of hTAM effect on neurons.

## Discussion

Beyond their primary role in neurotransmission, nerves are integral to various physiological processes, including proper organogenesis (2, 3), responses to external *stimuli*, and to tissue repair and regeneration in the adulthood (79). They also have a role in supporting tumor growth and metastasis (80) contributing, with nerve-secreted factors, to make tumors more aggressive (81). Consistent with this, several tumors have been shown to promote neural growth and infiltration generating a self-sustaining cycle supporting each other broadening. Tumor progression is also fostered by a dynamic relationship between stromal and cancer cells. In this context, a major contribution is provided by TAM (5, 6). TAM significantly contribute to the tumor microenvironment. Neurotransmitters have been shown to support the pro-tumoral functions of TAM (1, 82–84) however, TAM role in tumor innervation development remains unclear. In this work we report, for the first time, that TAM exhibit distinctive neurogenic properties and contribute to foster tumor innervation. Although we did not investigate if the increased nerve infiltration affected tumor aggressiveness, our *in vitro* and *in vivo* data now indicate a direct role of TAM in neurite outgrowth and axonal regeneration, suggesting the establishment of a vicious circle between TAM and neuronal growth during tumor progression. Notably, in this study we demonstrate that TAM “*neural growth*” signature not only is detected among different types of human cancers (e.g. GBM, breast, colon, pancreas, endometrial tumors), but is also preserved across different species (human and mouse) and detectable in *in vitro* settings.

Bioinformatic and functional validation experiments of this study characterized the mechanism behind the TAM interactions that produce these neurogenic effects. We identify SPP1 as a relevant factor produced by TAM and required to enhance nerve growth in tumor. SPP1 is known as major non-collagenous bone matrix protein involved in bone development and homeostasis (85), and in trophic/wound healing functions (86). SPP1 contributes to tumorigenesis and metastasis in different cancer types, including brain, breast, gastric tumors (87), mainly supporting tumor cell proliferation, adhesion and migration, immune response, and angiogenesis (87). SPP1 is included in the marker gene set identifying a specific TAM population linked to poor prognosis in different tumor types (88). By scRNA-seq data analysis, this population has been linked to metastasis, angiogenesis, and cancer cell stem cell activation (89, 90), but the underlying molecular mediators are undefined. This study now reveals a direct role of SPP1 as TAM-mediated neurogenic factor involved in cancer-dependent neural growth. We believe this is the first evidence indicating that SPP1 is not simply a marker of TAM subsets associated with negative prognosis (88), but rather a mediator directly supporting their negative prognostic value. Spp1 plays neurotrophic functions in developing (35) and adult brain (29), peripheral nervous system (44) and neurons (46, 53). Our *in vitro* results on different neuronal cultures, show Spp1-mediated direct role of both murine and human TAM on neural growth. Spp1 signaling has been shown to promote neural maturation by activating Rictor signaling pathway in neurons (46, 53). Our data suggested that Spp1, produced by TAM, mediates Rictor activation in neuronal cultures. Activation of the mTORC2 complex, induced by Rictor (77), improved the intrinsic axonal growth capacity in adult sensory neurons after injury (91). The proteomic data identified Rictor induction as a major TAM-induced regenerative effect in spinal cord parenchyma. Rictor knockdown reduces the locomotor recovery of TAM-transplanted mice, indicating that it is required for TAM effects on *in vivo* spinal cord repair. This is in line with previous evidence indicating that Rictor activation improves motor recovery in a rodent model of mild SCI (78).

The newly identified neurogenic property of TAM drives interest toward their potential significance as therapeutic tools in neurological disorders associated to CNS tissue loss. To this aim, we used a preclinical model of severe complete-compressive-contusive SCI lacking any spontaneous clinical recovery, characterized by severe neural tissue damage with secondary ischemia and deposition of ECM molecules (41). In this challenging experimental setting, a very poor tissue environment develops in the trauma area which is not conducive to regeneration and repair (58–61, 92, 93). Of note, this model is also associated with accumulation of chronically inflamed M1 macrophages, which persist throughout the late stages (58–61, 92, 93). Administration of M2 macrophages have been reported to have some therapeutic potential in a mild preclinical SCI model (26, 94, 95). However, this favourable macrophage phenotype was shown to be inadequate when tested in the human SCI setting, which is usually characterized by severe CNS injury (10, 62). Likewise, in our experimental setting of severe complete-compressive-contusive SCI, M2 macrophages-based therapy have no effect on tissue regeneration and motor recovery. On the contrary, the adoptive transfer of TAM during the SCI subacute phase supports a multiple molecular, cellular, and structural modifications which together contribute to promote neural tissue regeneration and a functional motor recovery. Results in TAM-treated animals confirm the remarkable TAM neuroregenerative potential as indicated by histological results, highlighting the significant increase in motor neurons and myelin content, and by the associated improvement of the electromyographic and clinical functionalities. Our data indicate that TAM administration has a positive action on most of the major pathological secondary events responsible for the instauration of the SCI hostile microenvironment, including promotion of ECM remodeling, reduction of tissue ischemia, and modulation of the inflammation (13, 96). In different tumors, TAM have been described to remodel the ECM by secreting several molecules, including the metalloprotease (MMP) 2 and 9 (13). In line with this evidence, our *in vitro* functional data and *ex vivo* transcriptomic and histological analysis, indicates that TAM, administered following CNS lesion, mediate the remodeling of the ECM composition reducing the expression in the cysts of key ECM components (Coll1, Coll3, fibronectin). This TAM-mediated ECM remodeling had a significant functional impact on scSCI neural tissue. Our X-ray tomography objectively demonstrated the transformation of large cysts, which initially separated the rostral and caudal portions of the spine, into smaller, multilobed structures (Fig. 4T, SMov. 3, 4) that no longer block the spinal cord tracts. In several tumors, including glioma or fibrosarcoma, tumor progression is also critically dependent on the formation of an immunosuppressive environment, and TAM have shown strong immunomodulatory properties (6). Accordingly, in our experimental setting TAM show *in vitro* immunomodulatory properties and increase the number of CD206^+^ macrophage/microglia cells, suggesting they may favour M1-to-M2 switch on endogenous macrophages. Our results confirm the capacity of TAM to produce several angiogenic molecules that promote tumor angiogenesis (97), both at the molecular and functional (zebrafish) level. After TAM adoptive transfer in severe SCI animals, we observe an increase of the spinal cord vascularization with beneficial effect on the neuronal tissue oxygenation, a critical aspect for neuronal survival (98, 99). We conclude that the therapeutic effect of TAM adoptive transfer following severe scSCI on healing the neural tissue involves direct effects on neural cells as well as profound changes in the SCI hostile microenvironment mediated by effects on stromal and immune cells. In addition, the safety assessment following single dose and repeated dose of TAM revealed no macroscopic or histological alterations associated with TAM administration demonstrating the safety of its adoptive transfer *in vivo*. TAM neurogenic property is present also in the human context. TAM “*neural growth*” molecular signature and neural growth properties are exhibited also by hTAM, which are characterized by up-regulation of pathways associated with neuro-regenerative effects, including ECM remodeling, angiogenesis, axogenesis, and immune-modulation. Moreover, as found in mouse, hTAM demonstrated to enhance neural growth by Spp1 effector, which mediate TAM neurogenic effect by inducing Rictor in neurons.

The additional neurogenic role of TAM in the tumor progression may be considered as novel therapeutic target to reduce neural addiction of cancer and might represent a novel strategy for the development of effective next-generation cancer treatments. Moreover, TAM may represent a new class of disease-modifying therapy for CNS lesions.

## Materials and Methods

### Bioinformatic analyses

#### Single-cell RNA bioinformatic analysis

On the 32,882 single-cell transcriptomes obtained from human glioblastoma (GBM), astrocytoma IDH+/+ or mutated IDH-/-tumor regions and the adjacent healthy area, the Seurat 5.0.1 scRNA-seq standard pipeline (100) was used to produce uniform approximation and projection (UMAP) plots. Raw data from human pancreatic, breast, and colorectal cancers, and data from mouse sarcoma were analyzed using the standard pipeline from Seurat 5.0.1. Gene marker expression on the different scRNA datasets was performed via the FeaturePlot function. Differential gene expression analysis for the extraction of the TAM’s DEGs was performed via the FindMarkers function. For the analysis of the enriched genes in the neural growth list, the AUCell package (101) and pipeline were used.

#### Neural growth gene list definition

Differentially expressed genes (DEGs) of TAM were identified using Seurat’s FindMarkers function (100) from the comparison between the TAM cluster (TGFBI^+^) and the CD14^-^ /TMEM119^-^ clusters included in the GBM, astrocytoma, and adjacent healthy tissue dataset. Enrichment analysis was then performed on these genes via the go_enrich function of the GOfuncR package (102). To describe this new finding, in accordance with previous studies (25, 28–30), we first include all the DEGs of the TAM cluster in the GBM datasets composing these GOs (102)To describe this new finding, in accordance with previous studies (25, 28–30), we first include all the DEGs of the TAM cluster in the GBM datasets composing these GOs (102). From the selected GO and the DEGs we obtain a list of 152 genes. We then refined this list by including only genes related to the cell secretome and itemised in the SEcreted Protein DataBase (SEPDB) (31), which resulted in a reduction of the list to 107 genes. We further manually refined this list to include only genes with well-established roles in neural growth-related functions, resulting in the definition of the “*neural growth*” gene list of 22 genes.

#### Cell sorting from mouse spinal cord tissue

Briefly, 0.5 cm scSCI mouse spinal cords centered to the lesion site were dissected and collected in tubes. The dissected spinal cord tissues were washed in ice-cold PBS 1X, resuspended in PBS 1X containing enzyme A, enzyme R and enzyme D (Multi Tissue Dissociation Kits, Miltenyi Biotec) and dissociated using the gentleMACS^TM^ Dissociator (Miltenyi Biotec) for 45 min (37C_Multi_F program). After dissociation, samples were centrifugated and TAM were enriched using the Percoll-density gradient method. Then, TAM were isolated using FACS cell sorter (BD Fusion, Becton Dickinson) using the fluorescent dye Zombie Yellow (BioLegend) as viability marker. In all the phases of the protocol the samples were kept in ice.

#### Mouse macrophage RNA sequencing and analysis

M2 and TAM mouse samples total RNA was isolated using DirectZOL RNA Miniprep kit (ZymoResearch) according to the manufacturer’s instruction. Quantification and quality check (RNA integrity number RIN>7) were assessed by using Qubit4 (Invitrogen) instrument. Libraries preparation and processing were performed with the kit SMARTer total stranded RNA seq. All samples were sequenced paired end with the Illumina NextSeq550 (sequencing kit: NextSeq 500/550 High Output v2.5 Kit; 150 cycles). The bcl2fastq v2.20.0.422 tool was applied to demultiplex samples and generate FASTQ files starting from base call (BCL) files. The quality of the fastq files was checked with fastQC version 0.11.8. Mapping and sorting were done with the STAR v.2.7.3a tool. The reference genome used was GRCm38 v.M23. BAM files were checked with the RSeQC v.3.0.1 quality control package. The count matrix was generated with feature Counts (103). Other RNA-Seq data were collected from the SRA database described in Chen et al. (104), and Bowman et al. (33) that were selected since they consist of monocytes and TAM samples free from any therapy or drug, induced gene expression or gene knockdown. The data from Chen et al. (104)and Bowman et al. (33) were then aligned to the same reference mouse genome. The count data were then subjected to DEGs analysis using DESeq2 package (105). Reads with low expression values were discarded. The considered DEGs are the ones with a significant p-value (<0.05). The gene set variation analysis was performed with the GSVA package (106), and the boxplots were computed using the package ggpubr; the statistical analysis between the different samples was computed by performing a Student’s t-test comparison. Gene ontology analysis was performed with the go_enrich analysis of the GOfuncR package (102).

#### Bioinformatic analysis on proteomic mouse gene datasets

The Principal Component Analysis (PCA) was performed with the DESeq2 R library, with function. Finally, the counts matrix was normalized generating cpm (counts per million) matrix. Gene ontology analyses was performed via the go_enrich function of GOfuncR package. The glmQLFit model was adopted and p-values were adjusted with the Benjamini Hochberg correction. Differentially expressed genes (FDR ≤ 0.05, |log2FC|≥ 1) were subsequently filtered considering both gene ontology and IPA annotation (Qiagen Ingenuity Pathway Analysis, v01-13) in order to define receptome and secretome genes lists. In particular, genes were selected if at least they were classified in one of the following categories: GO:0005576: extracellular region; GO:0005615: extracellular space; GO:0009986: cell surface; GO:0031012: extracellular matrix; GO:0005886: plasma membrane; GO:0016020: membrane; GO:0005125: cytokine activity; GO:0008083: growth factor activity: IPA: extracellular space; IPA: plasma membrane. Gene Ontology annotations were performed in R with AnnotationDbi v.1.44.0 and org.Mm.eg.dbv.3.7.0 packages. Up (and down) “exclusive” genes were listed considering the intersections of only up (or down) regulated genes in the TAM pre vs M2 pre and TAM pre vs TAM post differential analyses (genes). Furthermore, “exclusive” up (and down) regulated genes lists in TAM samples were functionally analyzed (gProfiler2 v.0.1.9 and msigdbr v.7.1.1 packages) and results were shown in bar plots.

#### Human macrophage RNA sequencing and analysis

For the *in vitro* M2, and TAM human samples total RNA was isolated using DirectZOL RNA Miniprep kit (ZymoResearch) according to the manufacturer’s instruction. Quantification and quality check (RNA integrity number RIN>7) were assessed by using Qubit4 (Invitrogen) instrument. Libraries preparation and processing were performed with the kit SMARTer total stranded RNA seq. All samples were sequenced paired-end with the Illumina NextSeq550 (sequencing kit: NextSeq 500/550 High Output v2.5 Kit; 150 cycles). The FASTQ files were then aligned with the reference genome GRCh38.p13. The count matrix was generated with feature Counts (103). Data of monocytes and TAM from breast and endometrial cancer described in Cassetta et al. (32) were already available as gene counts. The Principal Component Analysis (PCA) was performed with the DESeq2 R library (105), with Variance Stabilizing Transformation data. Genes were filtered by expression with the filterByExpr function. The count data were then subjected to DEG analysis using the DESeq2 package. Reads with low expression values were discarded. The considered DEGs are the ones with a significant p-value (<0.05). The gene set using the package ggpubr; the statistical analysis between the different samples was computed by performing a Student’s t-test comparison. Gene ontology analysis was performed with the go_enrich analysis of the GOfuncR package (102).

### In vitro studies

#### Murine bone marrow-derived macrophage (BMDM) generation and polarization

BMDMs were generated from male tdTomato mice (B6.Cg-*Gt(ROSA)26Sor^tm9(CAG-tdTomato)Hze^*/J or C57BL/6 mice, 8-12 weeks old). Briefly, after flushing femurs, cells were differentiated in Iscove’s Modified Dulbecco’s Medium (IMDM Euroclone, #ECB2072L) supplemented with heat-inactivated FBS (10%, Euroclone, #ECS0186L), penicillin/streptomycin (100 U/mL, Gibco, #15140-122), L-glutamine (2 mM, Gibco, #25030081) and M-CSF (100 ng/ml, Miltenyi Biotec, #130101704) for 7 days. To differentiate them in TAM, BMDM were cultivated in tumor conditioned medium (TCM), (107) and IMDM (1:1) and placed in hypoxia (1% O_2_) for 16-18 h at 37°C, 5%CO_2_. M1 macrophages were generated by conditioning BMDM with 100 ng/mL of LPS + 20 ng/mL of IFN-γ for 16-18 hours at 37°C, 5%CO_2_. M2 macrophages were generated by conditioning BMDM with IL-4 (20 ng/ml; Miltenyi Biotec) for 16-18 hours at 37°C, 5%CO_2_

#### Tumor conditioned medium (TCM) for TAM differentiation

10-week-old C57Bl/6 mice were injected intramuscularly in the left limb with 1 x 10^5^ MN-MCA1 cells. At day 21, animals were sacrificed, tumors were harvested and digested in 0.125% endotoxin free trypsin at 37°C for 40 min. Tumor-derived cells were filtered and plated in RPMI 1640 (Lonza, Basel, Switzerland) supplemented with 10% heat-inactivated FBS, 100 U/ml penicillin-streptomycin, and 2 mM L-glutamine. The cell-free supernatant was collected after 16 hours.

For human TAM polarization, TCM was obtained from the supernatant of GBM cell line U87 (ATCC), cultured with DMEM GlutaMAX (Gibco), supplemented with 10% heat-inactivated FBS (10%), 100 U/ml penicillin-streptomycin collected after 18h of cultured at 37°C, 5%CO_2_, and 80% cell confluence.

#### Co-culture of mouse neural stem cells (mNSCs) and mouse TAM

E14.5 mouse neural stem cells (mNSCs) were extracted from brain embryos and maintained in culture as previously described (49). Briefly, neurospheres were dissociated with StemPro™ Aldrich, #P6407)-coated coverslips on 24-well plates at a concentration of 4 x 10^4^ cells/well and cultured at 37°C, 5%CO_2_, with DMEM/F-12 GlutaMAX (Gibco, #31331-028) supplemented with B27 (2%; Gibco, #17504-044), N2 (1%; Gibco, #17502-048), penicillin/streptomycin (1%; Gibco) enriched with 20 ng/ml EGF and 20 ng/ml bFGF. At day 2, 15 x 10^3^ mouse TAM (+TAM) were added in each well of culture for 24h. Cells were then fixed in 4% PFA (Mondial), 4% sucrose and left in PBS 1X at 4°C for subsequent immunostaining analysis.

#### Isolation of single cell neurons from mouse dorsal root ganglia (DRG)

DRG were dissected from C57Bl/6 mice and placed on a Petri dish containing fresh HBSS, centrifuged at 1300 rcf and cultured at 37°C, 5% CO2, for 5 days into a 12-well plate in DRG medium composed by Neurobasal Medium-L-Glutamine (Gibco), supplemented with heat-inactivated FBS (10%, Lonza), B27 (2%, Gibco), penicillin-streptomycin-fungizone (1%; Lonza) and Glutamine 100x (0.5 mM; Gibco). DRG were then mechanically dissociated with collagenase (1%; Roche; #10103578001) and trypsin (0.25%; Sigma-Aldrich; T4665), filtered and cultured into a poly-lysine precoated 12-well plate in DRG Medium supplemented with NGF (50 μg/ml, DBA). Single cell neurons derived from DRG were seeded at a concentration of 6.5 x 10^4^ cell/well of a 12-well plate for 72 hours and they were allowed to differentiate for 5 days.

#### Co-culture of mouse single cell neurons isolated form murine DRG and mouse M2 and TAM

Mouse single cell neurons isolated from DRG were co-cultured adding 15 x 10^3^ mouse M2 (+M2 or TAM (+TAM) in each well. After 5 days of co-culture (37°C, 5% CO_2_), cells were fixed in 4% PFA (Mondial), 4% sucrose and left in PBS 1X at 4°C for subsequent immunostaining analysis.

#### TAM^LV Scramble^ and TAM^LV SPP1-KD^ generation and co-cultures with mNSCs

BMDM were differentiated in Iscove’s Modified Dulbecco’s Medium (IMDM) supplemented with heat-inactivated FBS (10%, Lonza), penicillin/streptomycin (100 U/mL; Lonza), L-glutamine (2 mM; Lonza) and M-CSF (100 ng/ml; Miltenyi Biotec) for 7 days. Cells were then transduced with a multiplicity of infection (MOI) of 10 with pLV[shRNA]-EGFP:T2A:Puro-U6>mSPP1-shRNA (VectorBuilder) or pLV[shRNA]-EGFP:T2A:Puro-U6>scramble-shRNA (VectorBuilder). The virus-containing medium from infected cells was removed 12 hours post-infection, TCM was added, and cultures were placed in hypoxia (1% O_2_) for 16-18 h at 37°C, 5%CO_2_ in order to obtain TAM^LV SPP1-KD^ or TAM^LV Scramble^ to 4 x 10^4^ mNSCs on poly-lysine (Sigma-Aldrich)-coated coverslips on 24-well plates. After 2 days, cells were fixed in 4% PFA (Mondial) 4% sucrose and left in PBS 1X at 4°C for subsequent immunostaining analysis.

#### Spp1 inhibition in mNSCs and mouse TAM co-culture (Parecoxib)

Spp1 inhibitor Parecoxib (163 μM; TargetMol) (108) was added to 4 x 10^4^ mNSC co-cultured with 15 x 10^3^ mouse TAM (+TAM+Parecoxib). The solvent of Parecoxib, DMSO, was added to 4 x 10^4^ control mNSCs (CTRL) and mNSCs co-cultured with 15 x 10^3^ mouse TAM (+TAM). After 2 days, cells were fixed in 4% PFA (Mondial) 4% sucrose and left in PBS 1X at 4°C for subsequent immunostaining analysis.

#### COX2 inhibition in mNSC and mouse TAM co-culture (Celecoxib)

COX2 inhibitor Celecoxib (25 μl/ml, LKT Laboratories) was added to 4 x 10^4^ control mNSCs (CTRL) and to mNSC co-cultured with 15 x 10^3^ mouse TAM (+TAM). After 2 days, cells were fixed in 4% PFA (Mondial) 4% sucrose and left in PBS 1X at 4°C for subsequent immunostaining analysis.

#### NSC^LV Scramble^ and NSC^LVRictor-KD^ generation and TAM co-culture

mNSCs were transduced with a (MOI) of 2.5 with pLV[shRNA]-EGFP:T2A:Puro-U6>mRictor (VectorBuilder) or pLV[shRNA]-EGFP:T2A:Puro-U6>mScramble (VectorBuilder). The virus-containing medium from infected cells was removed 12 hours post-infection. Co-culture was performed by adding to 4 x 10^4^ NSC^LV Scramble^ or NSC^LV Rictor-KD^, 15 x 10^3^ of mouse TAM on poly-lysine (Sigma-Aldrich)-coated coverslips on 24-well plates. After 2 days, cells were fixed in 4% PFA (Mondial) 4% sucrose and left in PBS 1X at 4°C for subsequent immunostaining analysis.

#### ELLA Mouse Immunoassay

TAM were generated and polarized *in vitro* for 7 days as previously described. Culturing supernatant was replaced with test media 24 hours prior to the experiment to avoid any confundant from exogenous proteins. The supernatant was then collected and processed for ELLA microfluidic immunoassay. 50 µl of supernatant was diluted 1:1 with sample diluent; the solution obtained was then added to the sample inlet of the ELLA cartridge, in accordance with the cartridge. Sample results were reported using Simple Plex Runner v.3.7.2.0 (ProteinSimple) and were available approximately 90 minutes after initiation of the run start. The experiment was conducted on 3 different supernatant collections.

#### Human monocyte-derived macrophage generation and polarization

Mononuclear cells were freshly isolated from leukocyte concentrate (buffy coat) collected from healthy donor volunteers enrolled at the Hematology Unit of the Humanitas Research Hospital, Rozzano, Milan, Italy. The biological material was anonymized. The monocyte population was enriched by positive selection of CD14-labeled target cells using a human magnetic antibody cell sorting (MACS) system (Miltenyi Biotec, Bergisch-Gladbach, Germany), according to the manufactureŕs procedure. The monocyte population was enriched using a human magnetic antibody cell sorting (MACS) system (Miltenyi Biotec), according to the manufactureŕs procedure. Collected monocytes were centrifuge at 1250 RPM for 5 minutes, supernatant discard, and cells plated in a Petri dish in different conditions according to the polarization. M0 were incubated at 37°C, 5% CO_2_ for 1 week in RPMI + FBS (10%) + M-CSF (100 ng/ml). M2 were incubated at 37°C, 5% CO_2_ for 1 week in RPMI + FBS (10%) + M-CSF (100 ng/ml) and IL-4 (20 ng/ml, added at day 6). TAM were incubated at 37°C, 5% CO_2_ for 6 days in a medium composed by TCM (30%) and RPMI (70 %) + FBS (10%) + M-CSF (100 ng/ml). Subsequently, TAM were incubated in hypoxia condition (1% O_2_) for 16-18 hours. All the polarized macrophages were detached using StemProAcutase 1X (Gibco) for 10 minutes at 37°C.

#### Generation of human induced pluripotent stem cells (iPSCs)

iPSCs were generated as previously described (48), reprogramming skin biopsy-derived fibroblasts or peripheral blood mononuclear cells (PBMCs) of three healthy donors. Briefly, fibroblasts were seeded in a well of a 6-well plate in specific fibroblast medium and PBMC isolated from peripheral blood samples, were seeded on 24-well plate in StemPro-34 medium (Thermo Fisher Scientific) supplemented with SCF (100 ng/ml), FLT-3 (100 ng/ml), IL-3 (20 ng/ml), and IL-6 (20 ng/ml) cytokines (all from Peprotech) with daily medium changes and fresh cytokines addition. After 4 days, transduction was performed using CytoTune-iPS 2.0 Sendai Reprogramming Kit (Thermo Fisher Scientific). Seven/three days later, reprogrammed fibroblasts and PBMCs were plated onto Matrigel coated dishes and thereafter monitored for the emergence of iPSC colonies. Colonies, passed using an EDTA solution (0.5 μM), were expanded in Essential 8 medium for at least six passages before being characterized and differentiated.

#### Motor neuron (MN) differentiation from iPSCs

iPSCs were differentiated to MNs as previously described (48). Briefly, iPSCs were allowed to grow in suspension in HuES medium supplemented with basic FGF (20 ng/ml; Peprotech) and Rho-associated kinase (ROCK) inhibitor Y27632 (20 μM; Selleckchem) to obtain embryoid bodies (EBs). The third day, SB431542 (10 μM, Tocris) and LDN193189 (0.2 μM, Sigma-Aldrich) were added to the cultures to induce neuralization. The fourth day, neural induction of EBs was performed using DMEM/F12 supplemented with L-glutamine (2 mM), penicillin (10 U/ml), streptomycin (10 μg/ml), MEM NEAA (0.1 mM), heparin (2 μg/ml; Sigma-Aldrich), N2 supplement (1%, Thermo Fisher Scientific), ROCK inhibitor (20 μM), ascorbic acid (AA, 0.4 μg/ml, Sigma-Aldrich), retinoic acid (RA, 1 μM; Sigma-Aldrich), brain-derived neurotrophic factor (BDNF, 10 ng/ml; Peprotech). SB431542 and LDN193189 were added until day 7 when cultures were supplemented with 1 μM smoothened agonist (SAG; Merck) and 0.5 μM purmorphamine (Pur; Sigma-Aldrich). For ten days medium was changed every alternate day. At day 17, EBs were dissociated with 0.05% trypsin (Sigma-Aldrich) and plated on poly-lysine (Sigma-Aldrich)/laminin (Thermo Fisher Scientific)-coated coverslips on 24-well plates at a concentration of 15 x 10^3^ cells/well. Cells were cultured in neural differentiation medium (Neurobasal, Thermo Fisher Scientific), L-glutamine (2 mM), penicillin (10 U/ml), streptomycin (10 μg/ml), MEM NEAA (0.1 mM), N2 supplement (1%), B27 (2%; Thermo Fisher Scientific), laminin (Lam; 1 μg/ml), glutamate (Glu; 25 μM; Sigma-Aldrich), ascorbic acid (0.4 μg/ml), glial-derived neurotrophic factor (GDNF, 10 ng/ml) and ciliary neurotrophic factor (CNTF, 10 ng/ml) (both from Peprotech). MNs were allowed to differentiate for 10 days and finally co-cultured with macrophages.

#### Differentiation of the SH-SY5Y Human Neuroblastoma cell line

SH-SY5Y cells (ATCC, CRL-2266) were differentiated and cultured using a combination of previously reported protocols (109, 110). Briefly, undifferentiated cells were maintained in basic growth medium, comprised of Dulbecco’s Modified Eagle’s Medium (DMEM) (Gibco Life Technologies, #11965-092), supplemented with 10% heat-inactivated fetal bovine serum (hiFBS) 1X penicillin/streptomycin (Gibco Life Technologies). To induce differentiation, cells were growth in T25 flasks at the density of 500.000/flask with Neurobasal-A media (Gibco Life Technologies) supplemented 1X B27 (Gibco Life Technologies), GlutaMAX Supplement (Gibco Life Technologies; #35050061) and 1X penicillin/streptomycin and Retinoic Acid all trans (RA) (10 µM; Med Chem) for an additional 20 days. On day 21 (DIV 21) induced cells were enzymatically detached and seeded onto poly-D lysin (0.5 mg/ml; Merck) and Laminin-coated (1:200 dilution; Merck) glass coverslips (12 mm^2^) at density 50.000/coverslip. The following day, the medium was changed to Neurobasal-A media (Gibco Life Technologies) supplemented with BDNF (50 ng/mL; Novus Bio/R&D Systems), potassium chloride (20 mM; Sigma), 1X B27 (Gibco Life Technologies), GlutaMAX Supplement (Gibco Life Technologies; #35050061) and 1X penicillin/streptomycin and RA for an additional 4 days. Cultivated cells were maintained below passage 12 at 37°C/5% CO_2_.

#### Co-cultures of iPSC-derived MNs and mouse M2 and TAM

iPSC-derived MNs were co-cultured adding 15 x 10^3^ mouse M2 (+M2), or mouse TAM (+TAM) in each well. After 48 hours of co-culture (at 37°C, 5% CO_2_), cells were fixed in 4% PFA (Mondial) 4% sucrose and left in PBS 1X at 4°C for subsequent immunostaining analysis.

#### Co-cultures of iPSC-derived MNs and human M2 and TAM

iPSC-derived MNs were co-cultured adding 15 x 10^3^ human M2 (+hM2), or human TAM (+hTAM) in each well. After 48 hours (37°C, 5% CO_2_), cells were fixed in 4% PFA (Mondial) 4% sucrose and left in PBS 1X at 4°C for subsequent immunostaining analysis.

#### Co-cultures of SH-SY5Y and human TAM

SH-SY5Y neurons were co-cultured 1:1 adding 5 x 10^4^ TAM (+TAM) in each well. After 48 hours of culture (37°C, 5% CO_2_), cells were fixed in 4% PFA (Mondial) 4% sucrose and left in PBS 1X at 4°C for subsequent immunostaining analysis.

#### Spp1 inhibition (Parecoxib) in SH-SY5Y and TAM co-culture

Spp1 inhibitor Parecoxib (163 μM; TargetMol) (108) was added to SH-SY5Y co-cultured with 5 x 10^4^ TAM at the beginning of the co-culture (+TAM+Parecoxib). The solvent of Parecoxib, (+TAM). After 2 days, cells were fixed in 4% PFA (Mondial) 4% sucrose and left in PBS 1X at 4°C for subsequent immunostaining analysis.

#### COX2 inhibition (Celecoxib) in SH-SY5Y and TAM co-culture

COX2 inhibitor Celecoxib (50 μM, LKT Laboratories) was added to control SH-SY5Y (CTRL) and SH-SY5Y co-cultured with 1:1 with 5 x 10^4^ TAM at the beginning of the co-culture. After 2 days, cells were fixed in 4% PFA (Mondial) 4% sucrose and left in PBS 1X at 4°C for subsequent immunostaining analysis.

#### Spp1 inhibition (siRNA) in human TAM and co-culture with SH-SY5Y or iPSC-derived MNs

On polarization’s day 5, *in vitr*o hTAM were incubated for 6 hours at 37°C with predesigned siRNA against human SPP1 (Qiagen, cat.#1027416) or Scramble (Qiagen, cat.# 1027417) and HiPerFect Transfection Reagent. Transfection reagents were washed away with culturing media and cultures let to recover for 24 hours. On polarization’s day 7, 5 x 10^4^ or 15 x 10^3^ transduced hTAM were added to SH-SY5Y or iPSC-derived MNs, respectively, prepared as described above. After 2 days, cells were fixed in 4% PFA (Mondial, cat. #FM0622) and left in PBS 1X at 4°C for subsequent immunostaining analysis.

#### mTORC2 selective inhibition in iPSC-derived MNs or SH-SY5Y co-cultures

To analyze the involvement of mTORC2 pathway, mTORC2 selective inhibitor JRAB2011 (10 μM; MedCheExpress) (111) was added to the culture medium of iPSC-derived MNs with DMSO (CTRL) colture, of SH-SY5Y with DMSO (CTRL) colture, of iPSC-derived MNs with DMSO co-cultured with 15 x 10^3^ mouse TAM (+TAM), and of SH-SY5Y with DMSO co-cultured with 5 x 10^4^ human TAM (+TAM) respectively at the beginning of the co-culture. After 2 days, cells were fixed in 4% PFA (Mondial) 4% sucrose and left in PBS 1X at 4°C for subsequent immunostaining analysis.

#### Oxygen deprivation Caspase 3 activation by immunofluorescence

SH-SY5Y differentiated neurons culturing medium were replaced with DMEM Low glucose medium supplemented with penicillin/streptomycin (1%, Gibco), GlutaMAX Supplement (Gibco Life Technologies; #35050061) and 1% BSA. 5×10^4^ human TAM were seeded and both transferred to hypoxic condition. After two hours, cultures were quickly fixed in 4% PFA and immunocytochemistry anti human β3-Tubulin and Caspase 3 was run, DAPI staining was performed. At least 10 of 20x zoom 1 fluorescence images were taken for condition and the number of β3-Tubulin positive cells and Caspase-3 and β3-Tubulin double positive cells was calculated.

#### Immunophenotyping

hTAM (2 x 10^5^ cells) were stained with the following antibodies: Anti-human CD14, VioBlue REAfinity (Miltenyi Biotec); Anti-human CD68, APC REAfinity (Miltenyi Biotec); Anti-human CD163, APC-Vio770 REAfinity (Miltenyi Biotec); Anti-human CXCR4, VioBright FITC REAfinity (Miltenyi Biotec); Anti-human HIF-1A, Alexa Fluor 647 (BD Biosciences); Anti-human VEGFA, PE (Abcam); Anti-hOsteopontin, APC (R&D). Mouse TAM (2×10^5^ cells) were stained with the following antibodies: Anti-mouse CD14, APC (eBioscience); Anti-mouse CD206, Alexa Fluor 647 (BD Biosciences); Anti HLA-DR, APC-Cy7 (BD Biosciences); Anti-mouse iNOS, FITC (Miltenyi Biotec) according to manufacture instructions. Data were acquired by MACSQuant10 (Miltenyi Biotec) and analyzed by FlowJo10 software. Mean Fluorescence Intensity (MFI) fold changes to negative controls were calculated for each marker.

#### Transwell migration assay

Transwell migration assays were performed in 24-well plates using transwell inserts accordingly to (112). Briefly, M2 and hTAM cells were seeded in RPMI-1640 medium with 25 ng/mL or 100 ng/mL CXCL12 and incubated at 37°C in 5% CO_2_ for 16 hours. Cells were fixed with 2% PFA and stained with DAPI (Thermo Fisher Scientific). Migrated cells were viewed with a fluorescence microscope (Nikon Eclipse Ni), imaged with a Nikon DS-2MBWc camera, and the number of DAPI nuclei counted using ImageJ software.

#### qPCR on human TAM and monocytes

The probe signal was normalized to an internal reference, and a cycle threshold (Ct) was taken significantly above the background fluorescence. The Ct value used for subsequent calculation was the average of two replicates. The relative expression level was calculated using transcript level of *Ptprc* as endogenous reference. Data analysis was done according to the comparative method followed by the User Bulletin No. 2 (Applied Biosystems).

#### In vitro immunofluorescence staining and quantification analysis

Immunofluorescent stainings on mNSCs, iPSC-derived MNs and single neurons obtained from DRGs were performed as previously described (49, 98). For the immunostaining of hTAM, 100,000 human monocytes were seeded on glass coverslips and polarized to obtain hTAM. On day 7 after hypoxia, hTAM were washed twice in cold PBS and fixed in PFA 4% at room temperature for 10 minutes. Briefly, slides were first incubated in blocking solution and then with primary antibodies. After they were incubated with the appropriate secondary antibodies, washed with PBS 1X and incubated with the nuclear dye. Slides were then mounted using DABCO. The quantitative analysis was done with the ImageJ software (National Institutes of Health, USA). The following primary antibodies were used for immunofluorescence analysis: anti-β3Tubulin (mouse, 1:400; Promega, cat. #G7121), anti-Caspase3 (rabbit, Cell Signaling, #9661), anti-CD14 (rabbit, Epigentek, #A72619), anti-CD68 (rat, 1:200; Invitrogen, cat. #14-0681-82), SPP1/OPN APC-conjugated (mouse, R&D; #IC14331A). Appropriate secondary antibodies were used: goat anti-mouse AlexaFluor 488 (1:500; ThermoFisher, cat. #A-11029, West Grove, USA), donkey anti-rabbit AlexaFluor 568 (1:500; ThermoFisher, cat. #A-21245, West Grove, USA), donkey anti-rat Alexa-Cy3 (1:1000; Jackson, cat. #712-166-153). Immunofluorescent images were acquired with a fluorescence microscope (Nikon Eclipse Ni or Nikon Eclipse Ti), a FluoView FV4000 confocal microscope or a LEICA SP confocal microscope equipped with 20x and 40x objectives. For each sample were analysed at least 3 fields for each glass coverslip in at least 3 different glass coverslips. In each field, a threshold was selected, and the results were normalized to the number of nuclei in the field considered.

#### Zymography analysis

Briefly, before and after macrophages conditioning, supernatants were collected, centrifuged at 300g, ultracentrifuged at 13000g and stored at -20°C until analysis. The protein content was quantified with Pierce BCA Protein Assay kit (Thermo Scientific) and diluted with 5X non-reducing sample buffer to a quantity of 40 µg of proteins/sample. Gels were then prepared as follows: 7.5% acrylamide separating gel containing gelatin (4 mg/mL) and stacking gel. After electrophoresis, gels were stained with Coomassie Brilliant Blue R250 (Sigma Aldrich) and acquired with Chemidoc MP Imaging System (Bio-Rad) and quantified with the Image Lab software (version 6.0.1; Bio-Rad).

#### Extracellular flux analysis

Oxygen Consumption Rate (OCR) and Extracellular Acidification Rate (ECAR) were measured in M2 and TAM using a Seahorse XF324 Extracellular Flux Analyzer (Agilent Technologies, Santa Clara, USA). On the day of the assay cells were seeded at the density of 7 x 10^4^ cells/well in a V7 XFe24-well cell culture microplate coated with poly-D-lysine and centrifuged at 400g for 2 minutes to attach them to the bottom of the plate. To measure mitochondrial OXPHOS, cells were incubated in Seahorse XF DMEM medium (Seahorse Bioscience, cat. #103575-100, North Billerica, USA) supplemented with 10 mM glucose, 1 mM sodium pyruvate, and 4 mM L-glutamine, pH 7.4, at 37°C in a non-CO_2_ incubator for 1 hour. OCR was measured at the baseline and after sequentially adding 1 µM oligomycin A, 1 µM carbonyl cyanide 4-(trifluoromethoxy) phenylhydrazone (FCCP) and 0.5 μM each of rotenone and antimycin A. To measure glycolysis, cells were incubated in Seahorse XF DMEM medium (Seahorse Bioscience, cat. #103575-100, North Billerica, USA) supplemented with 4 mM L-glutamine, pH 7.4, at 37°C in a non-CO_2_ incubator for 1 hour. ECAR was measured at the baseline and after sequentially adding 10 mM glucose, 1 μM oligomycin A, and 50 mM 2-deoxy-D-glucose. Data were normalized to the DNA content *per* well that was quantified with the CyQUANT cell proliferation assay kit (Thermo Fisher Scientific, cat. #C35007, Waltham, USA), accordingly to the manufacturer’s instructions, and analyzed as previously described (98).

#### Immune modulatory effect of TAM on M1 polarized macrophage, and FACS analysis

To investigate the immune modulatory effect of TAM on M1 polarized macrophage, primary murine macrophages (M1) from WT animals and tdTAM from B6.Cg-Gt(ROSA)26Sortm14(CAG-tdTomato)Hze/J mice were co-cultured. M1 macrophage polarization were obtained by adding to a differentiated M0 macrophage culture 100ng/ml of LPS (Sigma Aldrich) for 16 hours. M1 macrophages were plated alone or with co-culture with TAM (1 :10 TAM/M1 ratio) in RPMI, 10% FBS, 200 mM L-Glutammine and P/S for 24 hours. Thereafter, macrophages were detached with Accutase for 15 minutes at 37°C and analyzed by FACS analysis. Briefly, 2 x 10^5^ cells were permeabilized for intracellular cell staining according to Trascription Factor Staining kit (Milteniy Biotec). Permeabilized cells were stained with 1:100 anti-mouse iNOS FITC (Milteniy Biotec) for 1 hour at RT. tdTomato negative M1 macrophages were gated and FITC Median Fluorescence Intensity (MFI) were quantified by MACSQuant10 (Miltenyi Biotec). Data from n=7 independent experiments were collected and MFI of M1 with or without TAM co-culture were compared.

### In vivo studies

#### Tumor innervation in the murine tumor model

All experimental procedures were approved by the National Institute of Health (protocol N.949/2018-PR). In order to induce tumor formation, we used 3-MCA-derived sarcoma cell line MN/MCA1 (37), kindly provided by Francesca Consonni (40). MN/MCA1 were plated in a T75 flask in fresh RPMI supplemented with heat-inactivated FBS (10%), penicillin-streptomycin (1%), L-Glutamine (1%) for 24 hours. At the specific experimental time-point (day 0 and day 14), MN/MCA1 cells were washed with PBS, detached with trypsin, resuspended in fresh sterile PBS and injected intramuscularly in the left hindlimb of C57BL/6 mice. At day 0, control mice (CTRL) received 1 injection (100 µl/injection) of cell suspension (MN/MCA1, 10^5^ cells/mouse), TAM-treated mice received a combination of MN/MCA1 (10^5^ cells/mouse) and TAM (5 x 10^5^ cells/mouse, TAM-treated mice), or a combination of MN/MCA1 (10^5^ cells/mouse) and TAM^LV Scramble^ (5 x 10^5^ cells/mouse, TAM^LV Scramble^-treated mice), or a combination of MN/MCA1 (10^5^ cells/mouse) and TAM^LV SPP1-KD^ (5 x 10^5^ cells/mouse, TAM^LV SPP1-KD^-treated mice). At day 14, TAM-treated mice (+TAM) received a second injection (100 µl/injection) of TAM suspension (5 x 10^5^ cells/mouse), TAM^LV Scramble^-treated mice (+TAM^LV Scramble^) received a second injection (100 µl/injection) of TAM^LV Scramble^ suspension (5 x 10^5^ cells/mouse), TAM^LV SPP1-KD^-treated mice (+TAM^LV SPP1-KD^) received a second injection (100 µl/injection) of TAM^LV SPP1-KD^ suspension (5 x 10^5^ cells/mouse). At day 21 mice were sacrificed, and the tumor mass were fresh dissected and post fixed overnight in 4% PFA, 4% sucrose, and stored in 30% sucrose at 4°C for further analysis.

#### Zebrafish transection spinal cord injury model

Two days post fertilization (2 dpf) zebrafish embryos from the Tg(-3.1*neurog1*:GFP)*^sb2^* (113) were collected, staged according to the reference guidelines(114) and raised at 28°C in E3 medium fish water. Before manipulations, embryos were dechorionated, and anesthetized. Two days post fertilization (2 dpf) zebrafish embryos belonging from the Tg(-3.1*neurog1*:GFP)*^sb2^* (113) line, were anesthetized and spinal cord were transected at 10th spinal segment level, as previously reported (115). 25 x 10^4 tdTomato^TAM (injured+TAM) or dPBS (injured) were transplanted directly embryo, the extension of the lesion at 1 and 2 dpi was measured using the Image-J software (USA National Institutes of Health, USA).

#### TAM angiogenetic potency analysis in the zebrafish model

Zebrafish embryos at 2 dpf of the Tg(*fli1a*:GFP) *^y1^* line (116) were transplanted in the perivitelline region with 25 x 10^4^ TAM or with dPBS as a control. Analyses were done counting the number and the length of spikes in the sub-intestinal vessel using the Image-J software (USA National Institutes of Health, USA), as previously described (107).

#### Severe complete contusive compressive SCI (scSCI) mouse model

To maintain the colony of tdTomato mice following the 3R principle, we preferred to isolate tdTomato monocytes from male mice and therefore we chose to injury, and thus to transplant, male mice. Severe scSCI was performed as described previously (41). Briefly, 7-week-old C57BL/6 male mice were anesthetized and underwent a laminectomy at thoracic 11 (T11) vertebra. Then, by using a MASCIS Impactor a 5 g rod was dropped from 6.25 mm height and left in compression for 11 seconds. Subcutaneous injections of Baytril (10 mg/Kg) and Altadol (2 mg/Kg) were provided. Animal care was performed daily till the end of the experiment.

#### Cell transplantation

TAM- and M2-treated mice were anesthetized, and the injured spinal cord parenchyma was exposed. Each animal received 4 injections (1 μl/injection) of cell suspension (TAM-/M2-treated mice, 0.5 x 10^6^ cells/μl) or saline solution (vehicle-treated mice) in 4 points separated each other by 2 mm (0.5 mm depth) close to the lesion hematoma. All the animals received single (day 4 post injury) or multiple (3, 10 and 17 dpi or 3, 10, 17 and 24 dpi) transplantations of TAM/M2 or vehicle. Subcutaneous injections of Baytril (10 mg/Kg) and Altadol (2 mg/Kg) were provided the day of the surgery and for the first 2 days post transplantation (dpt).

#### Locomotor evaluation and ankle joint flexibility analysis

The locomotor function of vehicle-, TAM-, and M2-treated mice was evaluated by using the Basso Bettie and Bresnahan (BBB) score adapted to mice BMS (65) using a modified open-field BBB scale (0–21). All assessments were performed in blind condition by 2 observers. The locomotor 23, 24, 25, 28, 31, 32, 35, 36 and 37 dpi. For statistical analysis of the BBB score, the mean of the left and right hindlimb scores was taken to yield a single BBB score per mouse.

To evaluate the spasticity of the muscles involved in the ankle dorsiflexors and plantar flexors, the ankle joint flexibility analysis was performed from 14 dpi as previously described (41). The following scores were assigned: “0” to the lack of passive flexure (spastic condition) corresponding to an angle of 180° between the tibia and the foot, “0.25” to an angle of 135°, “0.5” to an angle of 90°, “0.75” to an angle of 45°, and “1” (complete passive flexure) corresponding to an angle of 0°. For statistical analysis of the BBB and the AJF score, the mean of the left and right hindlimb scores was taken to yield a single BBB/AJF score per mouse.

#### In vivo electromyographic recording

*In vivo* electromyographic (EMG) recording of spontaneous muscle activity was performed in *Tibialis Anterior* (TA) and *Lateral Gastrocnemius* (LG) muscles. In each muscle of anesthetized animals, we inserted thin stainless-steel wires with bare tips and connected them to the amplifier (CyberAmp 320, Axon Instruments). As reference electrode, a cut-short 20G stainless steel needle was inserted under the skin in the back of the animals and connected it to ground. The recording began 10 minutes after full recovery from anesthesia, and lasted 3 minutes for each muscle, during which time the animal was gently stimulated to move every 10-15 seconds. We estimated the amount of muscle paralysis by averaging the root mean square (RMS) values of the 10 largest EMG signals. Spasticity was estimated by calculating the duration of muscle activity relative to the total duration of recording. In total we recorded 31 TA muscles from 17 mice, and 29 LG muscles from 16 mice.

#### Corticospinal Tract Tracing in severe contusive SCI mouse model

3 days before the scSCI surgery, vehicle- and TAM-treated mice received tracer injections into the motor cortex areas. Mice were anesthetized and positioned in a digital stereotaxic apparatus (RWD Life Science) with the head held in place by ear bars. The scalp was retracted, and a hole was drilled above the motor cortex. After the brain parenchyma exposure, using the digital stereotactic apparatus 1 μl of 2 mg/ml biotinylated dextran amine (BDA; MW 10,000; Molecular Probes) in PBS was injected at 6 specific coordinates (3 sites/side) using a 10 µl Hamilton microsyringe. Mediolateral (ML) coordinates: ±1 mm lateral to Bregma; anteroposterior (AP) coordinates from microsyringe was left in place for 1 minute after the injection to allow tracer diffusion. The scalp was then sutured, and mice were allowed to recover. Subcutaneous injections of Baytril (10 mg/Kg) and Altadol (2 mg/Kg) were provided the day of the surgery and for the first 3 days post injection.

#### mTORC2 selective inhibition in severe scSCI mouse model

The day of TAM transplantation (3, 10 and 17 dpi), with a 10 μl Hamilton microsyringe connected to a digital stereotaxic apparatus (RWB Life Science), 4.8 x 10^10^ genome copies of custom-made recombinant AAV9-GFP-U6-mRICTOR-shRNA (Rictor^KD^ mice) (Biolabs, Malvern) or AAV9-GFP-U6-scramble-shRNA (Rictor^Scr^ mice) (Biolabs, Malvern) were injected (0.5 μl/injection) in 6 points of the exposed spinal cord parenchyma of injured mice. Injections were separated each other by 0.5 mm (0.5 mm depth) interspersed with the points of TAM transplantation.

### *Ex vivo* analyses

#### Tissue fixation and processing

Animals were intracardially perfused with 4% paraformaldehyde (PFA) and 4% sucrose. Spinal cords were extracted, incubated overnight in 4% PFA, 4% sucrose, and stored in 30% sucrose at 4°C. For histochemical analysis, 1.5 cm of dissected spinal cords (0.75 cm rostral and 0.75 cm caudal from the site of the lesion) were cryosectioned (25 μm-thick sections, for BDA imaging 80 μm-thick longitudinal sections) and stored at −20°C before analysis.

#### Ex vivo immunofluorescence, histochemical staining and quantification analysis

The staining for BDA was performed using the Alexa Fluor^TM^ 488 Tyramide SuperBoost^TM^ Kit (Invitrogen). Immunofluorescent and histochemical staining of TAM-, M2- and vehicle-spinal cord transversal and longitudinal sections were performed as previously described (41). The same protocol was used to perform immunofluorescent stainings on MN/MCA1, MN/MCA1+TAM, MN/MCA1+TAM^LV Scramble^ and MN/MCA1+TAM^LV SPP1-KD^ tumor samples. Briefly, for immunofluorescent stainings, slides were first incubated in blocking solution and then with primary antibodies. After they were incubated with the appropriate secondary antibodies, washed with PBS 1X and incubated with the nuclear dye. Slides were then mounted using DABCO for fluorescent/confocal microscopy. The following primary antibodies were used for 1, rat, 1:200; BD Biosciences, cat. #557355, Franklin Lakes, USA), anti-CD68 (rat, 1:200; Thermo Fischer Scientific, cat. #14-0681-82, Waltham, USA), anti-CD206 (goat, 1:400; R&D Systems, cat. #AF2535, Minneapolis, USA), anti-choline acetyltransferase (ChAT, goat, 1:200; Millipore, cat. #AB144P, Burlington, USA), collagen-I (Coll1, rabbit, 1.200; Abcam, cat. #AB34710, Cambridge, UK), collagen-III (Coll3, mouse, 1.200; GeneTex, cat. #GTX26310, Irvine, USA), anti-fibronectin (rabbit, 1:200; Dako, cat. #A0245, Nowy Sącz, Poland), anti-GFP (rabbit, 1:400; Thermo Fisher Scientific, cat. #A11122, Waltham, USA), anti-glial fibrillary acidic protein (GFAP, goat, 1:200; Abcam, cat. #AB53554, Cambridge, UK), anti-ionized-calcium binding adapter molecule-1 (Iba1, rabbit, 1:200; WAKO, cat. #019-19741, Osaka, Japan), anti-NeuN (mouse, 1:200; Millipore, cat. #MAB377, Burlington, USA), anti-neurofilament-200 (rabbit, 1:200; Sigma, cat. #N4142, St. Louis, USA), anti-peripherin (rabbit, 1:200; Millipore, cat. #AB1530), anti-RFP biotinylated antibody (rabbit, Rockland, cat. #600-401-379, Philadelphia, USA.). Appropriate secondary antibodies were used: donkey anti-goat AlexaFluor 488 (1:500; Jackson, cat. #AB-2340428, West Grove, USA), donkey anti-goat AlexaFluor 647 (1:500; Thermo Fischer Scientific, cat. #A21447, Waltham, USA), donkey anti-mouse AlexaFluor 488 (1:500; Thermo Fisher Scientific, cat. #A2120, Waltham, USA), donkey anti-mouse AlexaFluor 647 (1:500; Thermo Fischer Scientific, cat. #A32787, Waltham, USA), donkey anti-rabbit AlexaFluor 488 (1:500; Thermo Fisher Scientific, cat. #A21206, Waltham, USA), donkey anti-rabbit AlexaFluor 647 (1:500; Thermo Fischer Scientific, cat. #A32795, Waltham, USA), donkey anti-rat AlexaFluor 488 (1:500; Thermo Fischer Scientific, cat. #A11006, Waltham, USA), TO-PRO-3 (1:3000; Thermo Fisher Scientific, cat. #T3605, Waltham, USA) or DAPI (1:2000; Thermo Fisher Scientific, cat. #D-1306, Waltham, USA). For histochemical RFP and NF200 staining, briefly after endogenous peroxidase neutralization sections were incubated in blocking solution, and then with the anti-RFP or anti-NF200 antibody. After washing steps, slices were incubated with the appropriate secondary biotinylated antibody. Tissue sections were visualized using VectaStain ABC kit (Vector Laboratories, Rome, Italy) and developed in DAB (3,3-diaminobenzidine) peroxidase substrate (Sigma-Aldrich, Milan, Italy). Sections were then washed with H_2_O and stained with hematoxilin, dehydrated and mounted with Entellan (Merck Millipore) for light microscopy. For LFB staining, briefly sections were hydrated in EtOH solutions, incubated with 0.1% LFB solution (Sigma Aldrich) differentiated in Li_2_CO_3_ solution (Sigma Aldrich) and then dehydrated in EtOH solutions, cleared in xylene (Carlo Erba) and mounted with

#### Image analysis and quantification

The quantitative analysis was done with the ImageJ software (National Institutes of Health, USA). Immunofluorescent images were acquired with a fluorescence microscope (Nikon Eclipse Ni, Nikon Eclipse Ti), a FluoView FV4000 confocal microscope or a LEICA SP confocal microscope equipped with 20x and 40x objectives. Image quantification: n is defined as biological independent sample and indicated in the figure legend for each experiment; for each biological replicate we analysed the following number of technical replicates: *Tumor samples: i)* the absolute number of nerves was counted manually in the whole tumor slices in at least 8 section for each sample, *ii)* the NF200 content was evaluated in at least 5 fields (2658.10 µm x 2292,43 µm) in at least 8 sections (large image) for each tumor sample. In each field, a threshold was selected, and the results were normalized to the area of the field considered. *Spinal cords: i)* the Neun, ChAT, NF200, CD31, Iba1, CD68 and CD206 content was evaluated in at least 8 entire section of spinal cord for each sample; *ii)* the Coll 1, Coll3 and fibronectin content was evaluated in at least 3 entire section of spinal cord for each animal; *iii)* the Luxol content was evaluated in at least 14 section of spinal cord for each animal, *iv)* the BDA analysis was performed considering at least 6 spinal cord sections for each animal. The total number of positive cells was *in silico* (Neun, Iba1, CD206) or manually (ChAT) counted or a threshold was selected (Luxol, NF200, Coll 1, Coll3, fibronectin and BDA) and the results were normalized to the total number of nuclei in the slice (approximately 5414 ± 44 nuclei/slice; Neun, ChAT, Iba1, CD206), or the total area of the spinal cord slice (Coll 1, Coll3 and fibronectin), or a region of interest (ROI: 4429.92 µm x 1839.75 µm) centered at the cyst (NF200), or the dorsal columns region area of the spinal cord slice (Luxol) or the rostral/caudal regions (ROI:12246.63 µm x 2968.02 µm) of the spinal cord slice (BDA). Morphometric blood vessels analysis was performed using NIH ImageJ software on vehicle- and TAM-treated spinal cord fields (4 fields/animals) from transversal sections stained with the CD31 marker. Three to six mice were considered for the analysis; the exact number of mice is reported in each graph.

#### Analysis of tissue oxygenation

Tissue oxygenation analysis was performed evaluating the oxidizes thiols accordingly to (74). Briefly, mice were transcardially perfused with 1.85% Iodoacetamide (IAA) and 1.25% N-ethyl IAA, and 1.25% NEM. Spinal cords were dissected and after transversal cryosectioning, disulfides were reduced with 4mM Tris (2-carboxyethyl) phosphine hydrochloride (TCEP; Sigma Aldrich) in PBS for 1 hour. Those reduced thiols were then labelled with 7-diethylamino-3-(4’-maleimidylphenyl)-4-methylcoumarin (CPM; Sigma Aldrich) for 30 min. Samples were then washed and mounted with DABCO. The slice area was delimited with a region of interest (ROI) and the hypoxic areas were quantified with ImageJ software (National Institutes of Health, USA) as percentage of positive pixels in the ROI normalized to the total pixels of the ROI. For each independent biological sample at least 11 entire spinal cord section were analysed.

#### tdTomato M2/TAM distribution analysis

tdTomato M2/TAM distribution analysis was performed on M2/TAM-treated mice which received multiple transplantations (3, 10 and 17 dpi). Slices were stained with the nuclear marker DAPI (1:2000; Thermo Fisher Scientific, Waltham, USA) and the number of tdTomato M2^+^/DAPI^+^ cells or TAM^+^/DAPI^+^ cells in the lesion section was manually counted with the CellCounter plugin in NIH ImageJ software (National Institutes of Health, USA). At least 5 transversal injured spinal cord sections of M2- or TAM-treated mice were analyzed in technical replicates (glass slides) on at least 4 animals.

#### Ex vivo optical imaging

Fluorescent images were acquired by IVIS Spectrum optical imager (Perkin Elmer, Waltham, USA), equipped with a cooled (-90 °C) back-thinned and a back-illuminated CCD camera. Images calibrated in radiance units (p/s/cm2/sr) were corrected for dark measurements and cosmic rays. The total flux (p/s) was measured by drawing a region of interests (ROIs) over the images. Image acquisition, processing and analysis were performed with Living Image 4.5 (Perkin Elmer, Waltham, USA). Extracted organs (heart, lungs, pancreas, spleen, stomach, kidney, testes, bladder, skin, lymph nodes, brain, liver, gut, spinal cord) were imaged with the following image parameters: excitation filter = 535 nm, emission filter = 680 nm, automatic exposure time, aperture stop f = 2, binning B = 8 with a Field of View (FoV)=6.6 cm-13.0 cm. The line intensity profile (3-pixel width) of the fluorescent emission along the spinal cord were measured on the fluorescent images.

#### X-ray phase-contrast micro computed tomography

X-ray phase-contrast imaging was performed at the I13-2 beamline of the Diamond Light Source synchrotron radiation facility. Three-dimensional representations of unstained spinal cord samples were obtained through in-line phase-contrast projection imaging (117) and by using a single defocused-image phase retrieval algorithm (117). The volumetric reconstructions were performed using the dedicated micro computed tomography processing pipeline available at the beamline (70–73). X-ray phase-contrast imaging was performed at the I13-2 beamline of the Diamond Light Source synchrotron radiation facility. Three-dimensional representations of unstained spinal cord samples were obtained through in-line phase-contrast projection imaging (117) and by using a single defocused-image phase retrieval algorithm (117). The volumetric reconstructions were performed using the dedicated micro computed tomography processing pipeline available at the beamline (70–73). The cyst areas were manually segmented from the main volumes for subsequent analysis using a custom-designed plugin of ImageJ software (National Institutes of Health, USA).

#### Immunoblot analysis

Immunoblot analysis was performed as previously described (41). Briefly, spinal cord samples were homogenized in radioimmunoprecipitation assay (RIPA) buffer in the presence of protease and phosphatase inhibitors. After blocking, membranes were probed with the proper primary antibodies overnight and subsequently incubated with the appropriate horseradish peroxidase-conjugated secondary antibodies. ECL-based signals (Clarity Western ECL Substrate; Bio-Rad Laboratories) were acquired with Chemidoc MP imaging system (Bio-Rad Laboratories), normalized on total protein staining and analyzed with the Image Lab Software (version 6.0.1; Bio-Rad).

#### Quantitative Real Time PCR

Total RNA was extracted from fresh cell pellet using Genejet RNA purification kit (Thermofisher Scientific). cDNA was synthetized from 1 µg of total RNA using SuperScript reverse trascriptase (Thermofisher Scientific). Quantitative real-time PCR (RT-PCR) was carried out using TaqMan Universal PCR Master Mix on QuantStudio 5 RealTime instrument, and the relative amount of mRNA for PTPRC, CD14, CD206, CXCR4, GLUT1, MT2A, MT1X, ANGPTL4 and SPP1 genes (Hs04189704_m1, Hs02621496_s1, Hs07288635_g1, Hs00607978_s1, Hs00892681_m1, Hs02379661_g1, HS00745167_s1, HS01101123_g1; Hs04189704_m1, Hs00959010_m1, Applied Biosystem) was calculated as fold increase on PTPRC expression (internal control). Data were obtained using ΔΔ^−Ct^ method.

#### Proteomic analysis

Protein from spinal cord samples of 6 vehicle-, 7 TAM-, and 7 M2-treated mice were extracted in lysis buffer (150 mM NaCl, 1% Triton X-100, 50 mM Tris HCl pH 7.4, 0.5% sodium deoxycholate, and 0.1% sodium dodecyl sulfate in ddH2O supplemented with complete Roche protease inhibitor) (118) and their concentrations in the obtained supernatants, were determined using the DC Protein Assay (Bio-Rad Laboratories) according to the manufacturer’s instructions. Briefly, a paramagnetic bead approach was applied (119, 120), during which proteins were reduced with TCEP and alkylated with IAA, followed by a protein clean-up on. Afterwards, proteins were enzymatically cleaved using trypsin (1:50), and a peptide clean-up was performed using ACN. Finally, the peptides were eluted in two steps, first with 87 % ACN in 10 mM ammonium formate (pH 10) and then with 2 % DMSO (dimethyl sulfoxide) in water. Thus, two fractions of peptides were obtained, which were subsequently analyzed using a Q Exactive HF mass spectrometer (MS; Thermo Fisher Scientific), which was coupled to a liquid chromatography (LC, UltiMate 3000 RSLC nano system; Dionex) via a chip-based electrospray ionization source (TriVersa NanoMate; Advion) as described before (121).

#### Preclinical safety assessment (histopathology)

22 mice, 12 VEH and 10 TAM-treated subjected only to laminectomy, were examined after 1 year from the treatment. A complete necropsy was performed, and the following organs were collected for histopathology: spinal cord, brain, heart, lungs, liver with gallbladder, kidneys, spleen, pancreas, intestine, and bone marrow (vertebral bodies). Organs/tissues with grossly detectable lesions were also sampled. Samples were fixed in 10% neutral buffered formalin and routinely processed for paraffin embedding. 4μm thick sections were stained with hematoxylin-eosin and evaluated under a light microscope. A pathologist blinded to treatment groups performed the assessments.

### Statistical analysis

To calculate the number of animals needed for each experiment, we based our approach on the F-and one within-subject factor (time, with 17 measurements). The primary objective of this experiment is to study the interaction between these two factors. The effect size was set at f = 0.25, the type I error rate at 5%, the power at 80%, and the correlation between measurements at 0.5. Under these conditions, the minimum required sample size is 5 animals per group.

The effect of TAM multiple transplantation on locomotor functionality (BBB, BMS; Fig. 4) was estimated using a mixed-effects regression model. Fixed effects included time (dpi) and treatment, while mouse ID was treated as a random effect. The nonlinear relationship between BBB/BMS and time was modelled using a linear spline with one knot within each treatment group. A time x treatment interaction term was also included in the model. AJF (Fig. S4) was modelled by a linear mixed-effect model without splines. Fixed effects were time (dpi) and treatment; mice ID was the random effect. A time x treatment interaction term was also considered. In the two models, the significance of the interaction term was tested by comparing the models with and without the interaction term using a (one-tailed) F-test, with the F-statistic calculated based on the Kenward-Roger method. The only covariates considered in the analysis are treatment and time.

Each measurement was taken from n distinct biological samples, technical replicates for each distinct biological sample were indicated in the relative material and methods section. The number of distinct biological samples used in each experiment is indicated in the figure legends, at least three distinct biological samples per condition were used in each experiment. Statistical tests were performed using GraphPad Prism software (GraphPad). These tests included the unpaired two-sided Student’s t-test and the F-test for both one-way ANOVA and two-way ANOVA with repeated measures, followed by the (two-tailed) Tukey’s post-hoc test. We quantified the differences between the mean BBB values in the three groups by using the ratio of means as a measure of effect size (122). Data are presented as mean ± SEM. Statistical p values are indicated as exact values whenever suitable, or it is indicated by asterisks: *p < 0.05; **p < 0.01; ***p < 0.001; ****p < 0.0001.

## Study approval

All procedures on mice were approved by the National Institute of Health (protocol N.750/2019-PR) and by the Animal Ethics Committee of the University of Verona (CIRSAL, Centro Interdipartimentale di Servizio alla Ricerca Sperimentale).

Zebrafish embryos were raised and maintained under standard condition according to the national guidelines (Italian decree March 4, 2014, n. 26).

All human studies were approved by the Ethical Committee (Comitato Etico Indipendente, IRCCS Istituto Clinico Humanitas, protocol N.6b2b3.n.1ch). Ethical clearance for the use of human subjects were obtained from the designated health facility (Comitato Etico Indipendente, IRCCS Istituto Clinico Humanitas). Written informed consent was obtained from each person after receiving information about use of their tissue samples.

## Supporting information

Supplementary informations + Supp Figures and Tables

## Data availability

Data are available upon request to the corresponding author.

## Author contribution

S.D. I.D. F.B. designed the study, performed experiments, analyzed and interpreted data. L.M., A.C., E.B., participated in experimental design, performed the analysis and interpreted data. I.K., K.S. performed proteomic analysis. M.S. performed statistical analysis. F.Bo. performed bioluminescence imaging. E.S., S.G. F.C., M.D.C., N.M., S.F., P.B., N.P., G.P. assisted with cell culture, molecular analyses, animal work and histochemical analysis. M.T.S., C.C. and G.M. performed cell sorting. G.B. and L.M. performed the electrophysiology. A.P., A.Pe. performed zebrafish experiments. M.E., A.D. performed X-ray analysis. A.C., E.S., E.R. performed RNA sequencing and analysis. I.D., F.B., M.L., G.F., S.D. wrote the paper. All authors discussed results and commented on the manuscript. I.D and F.B. supervised major parts of the study; supervised the project.

## Acknowledgements

Project funded under the National Recovery and Resilience Plan (PNRR), Mission 4 Component 2 Investment 1.3 - Call for tender No. 341 of 15/03/2022 of Italian Ministry of University and Research funded by the European Union – NextGenerationEU Award Number: Project code PE0000006, Concession Decree No. 1553 of 11/10/2022 adopted by the Italian Ministry of of the nervous system in health and disease” (MNESYS). European Union project FETPROACT-2018-2020 HERMES [grant number 824164], is acknowledged for the support on research provided to I.D, G.C., G.P. and G.P., J.H. We thank the Italian patient associations GALM and La Colonna and the HEMERA s.r.l. for financial support. We thank the “Centro Piattaforme Tecnologiche-CPT”, E. Lorenzetto for the technical support, and “Centro Interdipartimentale di Servizio alla Ricerca Sperimentale-CIRSAL” of the University of Verona. K. Schubert and I. Karcossa are grateful for support by the UFZ-funded platform ProMetheus for proteome and metabolome analysis. M. Endrizzi is grateful for support by the Wellcome Trust 221367/Z/20/Z; EPSRC EP/R513143/1; National Research Facility for Lab X-ray CT (NXCT) through EPSRC grants EP/T02593X/1 and EP/V035932/1 and for the provision of beamtime MG32416 on ID13-2 at the Diamond Light Source, and the assistance of the beamline scientists there. We thank Prof. A. Mantovani for helpful suggestion on the manuscript; Prof. F. Marchesi for useful discussion; A. Mattè for helpuful suggestions on metalloproteases assays; R. Zamfir, L. Binaschi, A. Lattanzi, T. Marawanyika, C. Amoroso, G. Fierabracci, S. Zorzin, G. Pedrotti and N. Domenichini for technical assistance.

