## Supplementary informations + Supp Figures and Tables for "Tumor-associated macrophages enhance tumor innervation and spinal cord repair"

### SUPPLEMENTARY MATERIALS

#### Extended results 1

##### *In vitro*-generated tumor-associated macrophages (TAM) show *bona fide* TAM signature and functional properties

Tumor-associated macrophage (TAM) functions are supported by a specific gene expression signature that distinguishes them from other forms of macrophage activation (1). Though TAM share some similarities with alternative activated macrophages (also alluded to M2), they are endowed with unique angiogenic, ECM-remodeling, and immunomodulatory properties (2, 3). Starting from isolated mouse bone marrow monocytes, we established an *in vitro* setting mimicking the tumor microenvironment to generate macrophages whose own gene expression hallmarks and functional properties resemble *bona fide* TAM (Fig. S2). To assess TAM signature, we first performed a whole transcriptome analysis, and we found that TAM expressed *bona fide* TAM hallmarks (1) compared to the alternative activated M2 macrophages (Fig. S2A). In addition to the bulk-RNA sequencing, we characterized the different *in vitro* cultured macrophage polarizations also by FACS analysis (Fig. S2B). To assess the TAM functionality of the *in vitro*-generated cells, we first evaluated our analysis by examining their cellular metabolism, which enables TAM to adapt to nutrient-deprived and hostile microenvironments. TAM's gene ontology analysis highlighted an increased expression of genes related to glucose metabolism (Fig. S2C), including the allosteric regulator of glycolysis *Pfkfb3*, the glucose transporters *Slca2a1* and *Slc2a6*. In line with the gene expression data, metabolic evaluation of TAM revealed a significant switch to a glycolytic metabolism compared to M2 macrophages, as shown by basal glycolysis ($p=0.0002$, Fig. S2D) and glycolytic capacity ($p=0.0001$, Fig. S2E). We also observed a significant increase in the percentage of TAM glycolytic reserve ($p<0.0001$, Fig. S2F), a general decrease in mitochondrial respiration (Fig. S2G-K). However, when metabolism was interrogated in the absence of glucose, TAM showed a higher basal oxygen consumption rate than M2 (Fig. S2L). In this condition, following addition of 10mM glucose, TAM showed a strong glucose-mediated respiratory inhibition, the so-called Crabtree effect (4) ($p<0.0001$, Fig. S2L, L').

Taken together, these data indicated that TAM displayed high metabolic plasticity suggesting they may be able to survive and adapt to the injury-induced microenvironment.

We next tested other TAM functional properties, including angiogenesis, ECM remodeling, and immunomodulation (5). Angiogenic properties were investigated in a Tg(*fli1a*:GFP)$^{y1}$ zebrafish model, which allows direct visualization of vessels (6, 7). One day after TAM transplantation in perivitelline space of Tg(*fli1a*:GFP)$^{y1}$ zebrafish, a significant increase in ectopic sprouting of vessels outside the basket-like structure of the sub-intestinal vessel was observed as compared to vehicle-injected animals, thus confirming TAM angiogenic properties (p=0.0025 and p=0.024, respectively; Fig. S2M-O).

TAM have been shown to support ECM remodeling by producing several metalloproteases (MMP) (5); accordingly, gene expression analysis indicated that TAM highly expressed MMP genes and genes related to ECM remodeling (Fig. S2C). We then analyzed the activity of secreted MMPs and confirmed that, when compared to M2 macrophages, TAM significantly increased the activity of pro-MMP9 (p=0.0002; Fig. S2P, Q) and MMP9 (p=0.011; Fig. S2P, Q). No differences were found in pro-MMP2 (p=0.14; Fig. S2P, Q) and MMP2 activity (p=0.24; Fig. S2P, Q).

Finally, gene ontology analysis of TAM compared to M2 indicated up-regulation of ontologies involved in the modulation of immune responses (including *Irak2, Havcr2, Ticam2, Ripk3, Itgal, Il18rap, Nectin2, Treml2, Itgam, Tlr2, Fcgr4, Fpr2, Fcgr1, Il4ra, Trem1, Tlr5, Mif, Il15, Tnfsf13b, Il18, Il1b*), leukocyte migration (including *Il16, Mif, Cxcl16, Fn1, Mmp14, Vegfa, Il1b, Cxcl1, Cxcl2, Pf4, Cxcl3, Thbs1*) (Fig. S2C), and down-regulation of genes included in the adaptive immune response (including *Il1rl1,Il27ra, Tfrc, Fyn, Il18r1, Anxa1,Ly9, Lat2, Adgre1, Hras, Alcam*) (not shown). To assess a direct immunomodulatory effect of TAM, we measured the capability of TAM to decrease *in vitro*-generated M1 macrophage inflammatory phenotype. M1 macrophage polarization is characterized by the increase of inducible nitric oxide synthase (iNOS) (Fig S2R-T) (5). We quantified by FACS analysis, iNOS protein content in M1 exposed or not with TAM co-culture. We observed that following 24h of co-culture with TAM, the M1 iNOS content statistically decreased (p=0.024; Fig. S2T), suggesting a reduced M1 activation.

Taken together, *in vitro* results showed that TAM displayed relevant properties for neural tissue regeneration including metabolic plasticity, ECM remodeling and immunomodulatory properties.

#### Extended results 2

##### PRECLINICAL LONG-TERM (ONE YEAR) ASSESSMENT FOR TUMORIGENICITY AND TOXICITY CONFIRMED THE SAFETY OF ADOPTIVE TRANSFER OF TAM IN SCI

Given the multiple and robust mechanisms of action of TAM on the spinal cord, to highlight the translational potential of TAM for CNS regeneration, we evaluated the long-term toxicity and tumorigenic potential following single- and repeated-intraparenchymal administration of TAM in the spinal cord of mice that underwent laminectomy. One year after the transplantation, we performed a necroscopy, which includes comprehensive anatomical and histopathological investigations after single or repeated administration of vehicle or TAM. Furthermore, in all studied animals, we monitored weekly body weight and behavioral/clinical signs, and we did not observe weight or behavioral alterations, including locomotor functionality (BBB = 18 in all animals observed, data not shown). We did not observe macroscopic and histological alteration associated with the treatment of TAM in the spinal cord (multiple sections) and brain. There is evidence related to the mechanical injury following the surgical procedure of laminectomy, the presence of focal, minimal mineralization in the spinal cord of a control animal (Fig. S8A), and epidermal cysts observed in the spinal cord of three animals (two control and one TAM treated mice) (Fig. S8A). Considering non-tumoral lesions (Fig. S8B), no major histopathological findings were detected in any of the other organs examined. Inflammatory cell infiltrates were commonly detected in the lungs (Fig. S8A), liver, and kidneys, with no differences in their frequency or severity among groups. Other lesions were occasionally detected in a single or a couple of animals and were considered incidental findings. Tumoral lesions were detected in 3/10 TAM-treated animals and in 2/12 vehicle-treated mice (Fig. S8A-C). Importantly, none of these tumors was of myeloid/macrophagic origin. Overall, these data suggested that the adoptive transfer of TAM did not show evidence of systemic toxicity and was non-tumorigenic *in vivo*.

#### Extended results 3

##### *In vitro*-generated human TAM show *bona fide* TAM signature and functional properties

We first validated gene and protein expression profiles and functional properties of human *in vitro*-generated TAM (hTAM). Bulk-RNA sequencing of *in vitro*-generated hTAM and hM2 highlights, via PCA, the different gene profiles between hTAM and hM2 (Fig S10A). As murine TAM (Fig. S2A), *in vitro*-generated hTAM, enriched TAM's hallmarks as identified by Qian et al. (1) when compared to hM2 (Fig. S10B), confirming their TAM-like signature. FACS analysis of human *in vitro*-generated TAM and M2 furthermore highlights the expression of TAM's markers (including CD14, CD163, etc., Fig. S10C) compared to M2 macrophages. We confirmed the TAM signature of hTAM by quantifying the induction of gene expression by q-PCR of TAM markers, including CD206, Glut1, HIF-1a, CXCR4, and VEGFA (Fig. S10D). To further highlight the different phenotypes of hTAM cells compared to hM2, we performed a gene ontology analysis. Compared to hM2, hTAM upregulate gene ontologies (Fig. S10E) related to wound healing (including secreted ADM, BNIP3, PDGFB, VEGFA), angiogenesis (including RORA, and secreted VEGFA, ANGPTL4, CXCL8, APLN), detoxification and regulation of defence response (including NDRG1, MT1E, MT1F, MT1G, MT1H, MT1X, MT2A, MT3, DDIT4, NUPR1), response to hypoxia (including HK2, PFKFB3, SLC2A1, SLC2A3, CXCR4, PLIN2, RORA, NDRG1, EGLN3, MT3, PLOD2, and secreted ADM, BNIP3, HILPDA, ANGPTL4), ECM-remodeling (including SULF2, EGLN3, PLOD2 and secreted MMP9, VCAN, FGF11, CXCL8, PDGFB, VEGFA, ANGPTL4), neuronal survival and myelination (including MT3, JAM2, VLDLR, NUPR1, EGLN3) (secreted molecules identified accordingly to Wang et al. (8)) (Fig. 6A, and Fig. S10E). In scSCI, mouse TAM could migrate in response to injury-induced signals. We tested whether hTAM maintained a chemotactic response to CXCL12, a relevant injury-induced signal, by using a transwell assay (S10F). Results showed that hTAM had a significantly higher chemotactic response to CXCL12 than hM2 macrophages (Fig. S10G). Our analysis indicated that *in vitro*-generated human TAM consistently retained *bona fide* TAM signatures and properties.

### SUPPLEMENTARY FIGURES AND LEGENDS TO SUPPLEMENTARY FIGURES

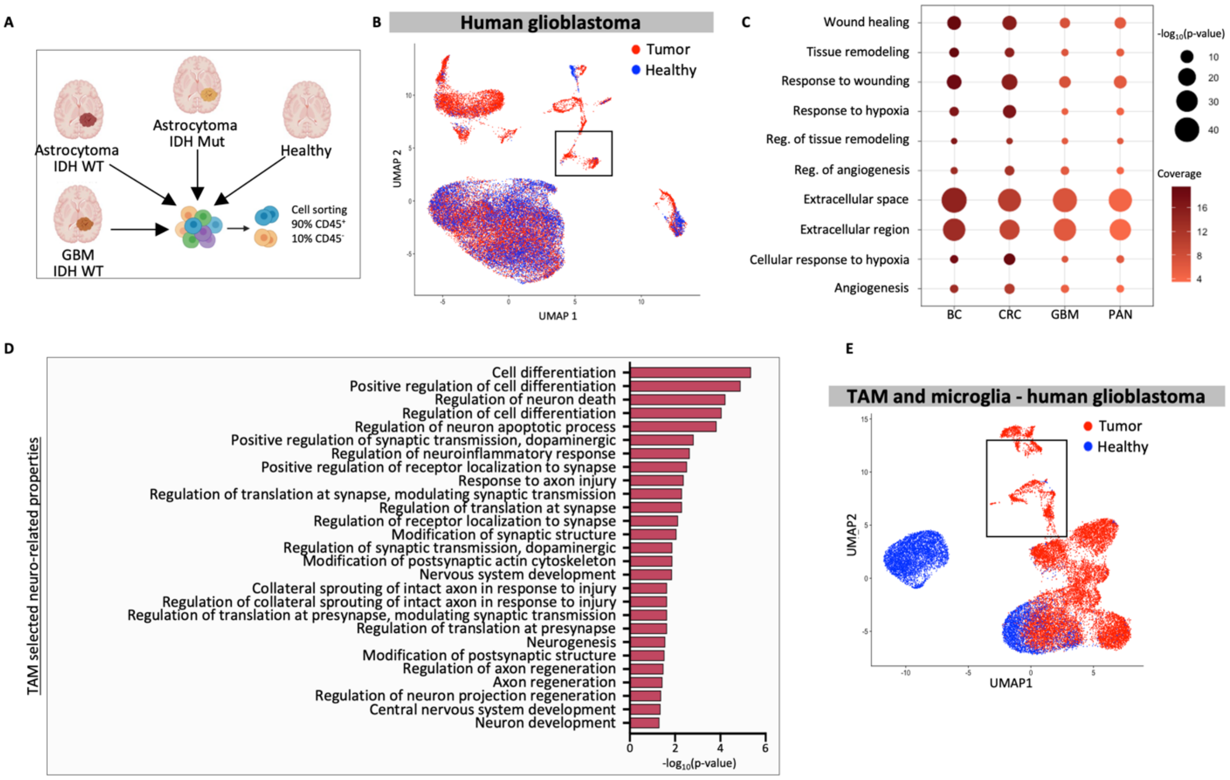


FIGURE S1

**Legend to Figure Supplementary 1: Single cell RNAseq analysis of human glioblastoma and TAM signature.**

**A)** Schematic representation of the workflow from tissue extraction to scRNA sequencing of human glioblastoma (GBM), astrocytoma, and healthy tissue samples. **B)** UMAP plot on human glioblastoma, astrocytoma, and healthy tissue cells. Red cells derive from glioblastoma or astrocytoma; blue cells derive from healthy tissue. The black square indicates the TAM cluster. **C)** Bubble plot showing the GOs expression related to TAM properties in all the human scRNA datasets considered. The bubble color represents the coverage of the GO, and the size represents the selected GO's p-value ($-\log_{10}$(p-value)). BC: Breast cancer, CRC: Colorectal cancer, GBM: glioblastoma, PAN: Pancreatic cancer. **D)** Neural growth-related gene ontologies (GOs) upregulated in TAM cluster compared to all the $CD14^-$ cells in the human glioblastoma/astrocytoma dataset. The represented GOs are the ones used for the definition of the "neural growth" gene list. **E)** UMAP plot on the $TGFBI^+$ (TAM) or $TMEM119^+$ (microglia) cells included in the human glioblastoma dataset. Red cells derive from glioblastoma or astrocytoma; blue cells derive from healthy tissue. The black square indicates the TAM cluster.

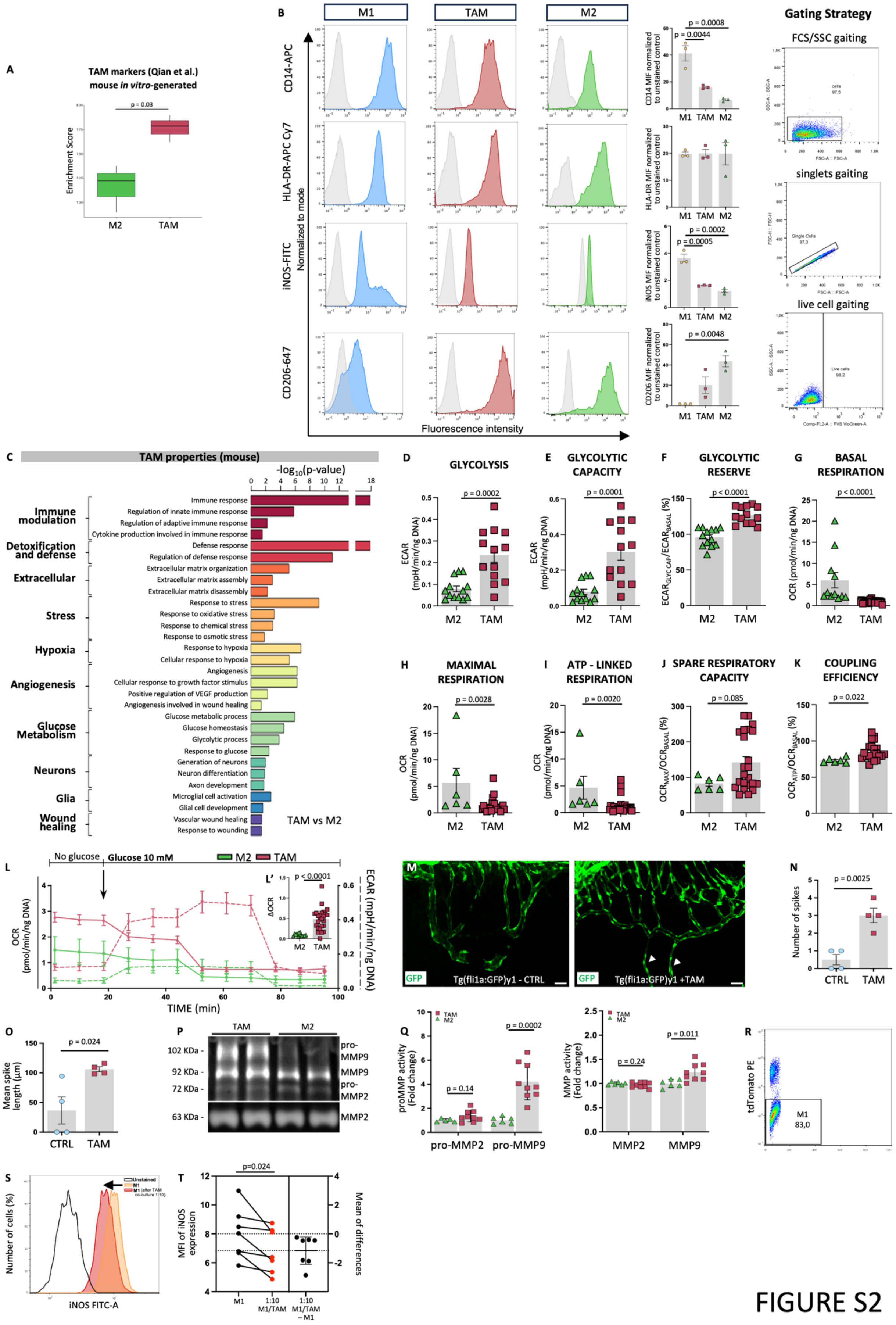
A
TAM markers (Qian et al.)
mouse *in vitro*-generated
p = 0.03
Enrichment Score
M2
TAM
B
M1
TAM
M2
CD14-APC
HLA-DR-APC Cy7
iNOS-FITC
CD206-647
Normalized to mode
Fluorescence intensity
CD14 MIF normalized to unstained control
p = 0.0008
p = 0.0044
HLA-DR MIF normalized to unstained control
iNOS MIF normalized to unstained control
p = 0.0002
p = 0.0005
CD206 MIF normalized to unstained control
p = 0.0048
Gating Strategy
FCS/SSC gaiting
singlets gaiting
live cell gaiting
C
TAM properties (mouse)
-log10(p-value)
Immune modulation
Detoxification and defense
Extracellular
Stress
Hypoxia
Angiogenesis
Glucose Metabolism
Neurons
Glia
Wound healing
Immune response
Regulation of innate immune response
Regulation of adaptive immune response
Cytokine production involved in immune response
Defense response
Regulation of defense response
Extracellular matrix organization
Extracellular matrix assembly
Extracellular matrix disassembly
Response to stress
Response to oxidative stress
Response to chemical stress
Response to osmotic stress
Response to hypoxia
Cellular response to hypoxia
Angiogenesis
Cellular response to growth factor stimulus
Positive regulation of VEGF production
Angiogenesis involved in wound healing
Glucose metabolic process
Glucose homeostasis
Glycolytic process
Response to glucose
Generation of neurons
Neuron differentiation
Axon development
Microglial cell activation
Glial cell development
Vascular wound healing
Response to wounding
TAM vs M2
D
GLYCOLYSIS
p = 0.0002
ECAR (mpH/min/ng DNA)
E
GLYCOLYTIC CAPACITY
p = 0.0001
F
GLYCOLYTIC RESERVE
p < 0.0001
G
BASAL RESPIRATION
p < 0.0001
OCR (pmol/min/ng DNA)
H
MAXIMAL RESPIRATION
p = 0.0028
I
ATP - LINKED RESPIRATION
p = 0.0020
J
SPARE RESPIRATORY CAPACITY
p = 0.085
K
COUPLING EFFICIENCY
p = 0.022
L
No glucose
Glucose 10 mM
M2
TAM
L'
p < 0.0001
ΔOCR
OCR (pmol/min/ng DNA)
ECAR (mpH/min/ng DNA)
TIME (min)
M
GFP
Tg(fli1a:GFP)y1 - CTRL
Tg(fli1a:GFP)y1 +TAM
N
p = 0.0025
Number of spikes
CTRL
TAM
O
p = 0.024
Mean spike length (µm)
P
102 KDa -
92 KDa -
72 KDa -
63 KDa -
pro-MMP9
MMP9
pro-MMP2
MMP2
Q
p = 0.14
p = 0.0002
proMMP activity (Fold change)
pro-MMP2
pro-MMP9
p = 0.24
p = 0.011
MMP activity (Fold change)
MMP2
MMP9
R
tdTomato PE
M1
83,0
S
Unstained
M1
M1 (after TAM co-culture 1:10)
Number of cells (%)
iNOS FITC-A
T
p=0.024
MFI of iNOS expression
Mean of differences
M1
1:10 M1/TAM
1:10 M1/TAM – M1

FIGURE S2

**Legend to Figure Supplementary 2: *In vitro*-generated mouse TAM resemble *bona-fide* TAM signature and functional properties.**

**A)** Single sample gene set enrichment analysis of TAM hallmarks defined by Qian et al. in *in vitro*-generated mouse M2 and TAM showing enriched expression in TAM group. **B)** (Left) FACS analysis of mouse *in vitro*-generated M1, TAM, and M2, showing different expressions of CD14, HLA-DR, iNOS, and CD206 markers. (Right) Panel showing the gating strategy used. **C)** Barplots showing enriched upregulated gene ontologies (GOs) in mouse *in vitro*-generated TAM compared to M2. Barplot length is the $-\log_{10}$(p-value) returned from the enrichment analysis. **D)** Basal glycolysis of mouse *in vitro*-generated M2 and TAM expressed in mpH/min/ng DNA. TAM showed a higher basal glycolysis compared to M2 (n=13/condition). **E)** Glycolytic capacity of mouse *in vitro*-generated M2 and TAM expressed in mpH/min/ng DNA. TAM showed higher glycolytic capacity compared to M2 (n=13/condition). **F)** Glycolytic reserve of mouse *in vitro*-generated M2 and TAM expressed as glycolytic capacity/basal glycolysis x 100. TAM showed higher glycolytic reserve compared to M2 (n=13/condition). **G)** Basal respiration of mouse *in vitro*-generated M2 and TAM expressed in pmol/min/ng DNA. TAM showed reduced basal respiration compared to M2 (M2: n=11; TAM: n=43). **H)** Maximal respiration of mouse *in vitro*-generated M2 and TAM expressed in pmol/min/ng DNA. TAM showed reduced maximal respiration compared to M2 (M2: n=6; TAM: n=26). **I)** ATP-linked respiration of mouse *in vitro*-generated M2 and TAM expressed in pmol/min/ng DNA. TAM showed reduced ATP-linked respiration compared to M2 (M2: n=6; TAM: n=31). **J)** Spare respiratory capacity of mouse *in vitro*-generated M2 and TAM expressed as $OCR_{MAX}/OCR_{BASAL}$ x 100. TAM showed a positive trend of increase compared to M2 (M2: n=6; TAM: n=23). **K)** Coupling efficiency of mouse *in vitro*-generated M2 and TAM expressed as $OCR_{ATP}/OCR_{BASAL}$ x 100. TAM showed increased coupling efficiency compared to M2 (M2: n=6; TAM: n=22). **L, L')** Representative OCR (expressed in pmol/min/ng DNA, continue lines) and ECAR (expressed in mpH/min/ng DNA, dotted lines) plot extrapolated from glycolysis stress test in M2 (green triangles) and TAM (red squares). TAMs showed a significant increase compared to M2 (quantification in L': M2: n=13; TAM: n=25). Data are normalized to total DNA content per well. **M)** Representative immunofluorescence images of the basket-like structure of the sub-intestinal vessel of 2 days post-fertilization in control (CTRL) and in TAM-treated in Tg(*fli1a:*GFP)$^{y1}$ zebrafish embryos. Scale bar = 50µm. **N)** Number of spikes sprouting outside the basket-like structure of the sub-intestinal vessel 2 days post-fertilization in CTRL and TAM-treated Tg(*fli1a*:GFP)$^{y1}$ zebrafish embryos. TAM-treated embryos showed higher number of spike sprouting outside the basket-like structure compared to CTRL (n=4/condition). **O)** Mean spike length of the vessels sprouting from the basket-like structure of the sub-intestinal vessel 2 days post-fertilization in CTRL and in TAM-treated Tg(*fli1a*:GFP)$^{y1}$ zebrafish embryos. TAM-treated embryos showed higher spike length compared to CTRL (n=4/condition). **P)** Representative gelatine zymogram of matrix metalloproteinases (MMPs) contained in the medium supernatants of mouse *in vitro*-

generated M2 and TAM. MMPs are indicated with their specific molecular weight (kDa). **Q)** Activity of pro-MMP2 and pro-MMP9 forms, and of MMP2 and MMP9 forms, expressed as fold-change respect to M2 medium supernatants. TAMs show consistently higher activity compared to M2 (pro-MMP2: M2: n=5; TAM: n=9; pro-MMP9: M2: n=6; TAM: n=9; MMP2: M2: n=6; TAM: n=9; MMP9: M2: n=6; TAM: n=9). Metabolic data are normalised to total DNA content per well. **R)** FACS analysis of mouse tdTomato$^{+}$ cells in M1 and tdTomato-TAM co-culture. M1 macrophages were negatively gated for tdTomato. **S)** Representative flow cytometry showing FITC-labelled iNOS expression in M1 macrophages and M1 macrophages co-cultured with TAM (10:1 ratio). **T)** Median Fluorescence Intensity (MFI) of iNOS expression in M1 macrophages and M1 macrophages co-cultured with TAM (10:1 ratio) (n=7/condition). Data are expressed as mean±SEM. Statistical analysis was performed via Student t-test.

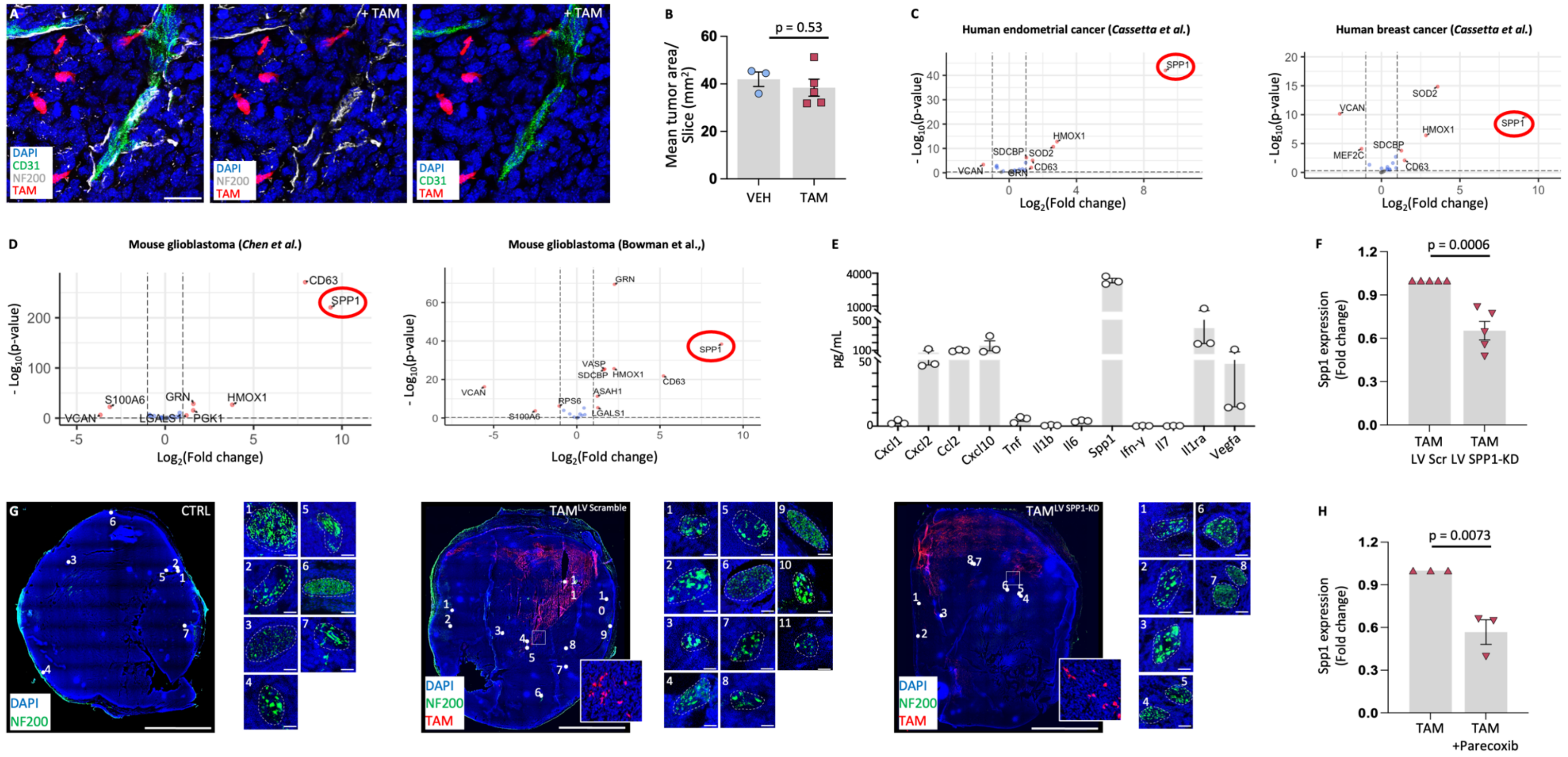
A
DAPI
CD31
NF200
TAM
+ TAM
B
p = 0.53
Mean tumor area/ Slice (mm²)
VEH
TAM
C
Human endometrial cancer (Cassetta et al.)
Human breast cancer (Cassetta et al.)
SPP1
HMOX1
SDCBP
SOD2
CD63
VCAN
GRN
MEF2C
- Log10(p-value)
Log2(Fold change)
D
Mouse glioblastoma (Chen et al.)
Mouse glioblastoma (Bowman et al.,)
CD63
S100A6
HMOX1
LGALS1
PGK1
VASP
RPS6
ASAH1
E
pg/mL
Cxcl1
Cxcl2
Ccl2
Cxcl10
Tnf
Il1b
Il6
Spp1
Ifn-γ
Il7
Il1ra
Vegfa
F
p = 0.0006
Spp1 expression (Fold change)
TAM LV Scr
TAM LV SPP1-KD
G
CTRL
TAM LV Scramble
TAM LV SPP1-KD
H
p = 0.0073
TAM +Parecoxib

FIGURE S3

**Legend to Supplementary Figure 3: TAM enhance nerve growth in mouse sarcoma via SPP1.**

**A)** Confocal immunofluorescence maximum intensity projections of z-stack transversal sections of sarcoma tumor from CTRL and TAM-treated mice stained with NF200 (white), CD31 (green) and DAPI (blue). TAMs are shown in red. Scale bar =25µm. **B)** Mean tumor area in transversal sections of sarcoma tumor from CTRL and TAM-treated mice. VEH- and TAM- treated mice showed no differences in the mean tumor area (VEH: n=3; TAM: n=5). **C)** Volcano plot showing the expression genes included in the "neural growth" gene list in TAM and monocytes isolated from human breast cancer and human endometrial cancer. Genes are separated according to their log2-transformed intensity fold-change and their -log10-transformed p-value (y-axis); the selected candidate gene (SPP1) is circled in red. **D)** Volcano plot showing the expression genes included in the "neural growth" gene list in TAM and monocytes isolated from different datasets of mouse glioblastoma. Genes are separated according to their log2-transformed intensity fold-change and their -log10- transformed p-value (y-axis); the selected candidate gene (SPP1) is circled in red. **E)** ELISA analysis showing the relative concentration of CXCL1, CXCL2, CCL2; CXCL10, TNF, IL1B, IL6, OPN/SPP1, INF-γ, IL7, IL1ra, VEGF in the supernatant of mouse *in vitro*-generated TAM (n=3/condition). *In vitro*-generated TAM showed higher concentration of SPP1 in the supernatant. **F)** Expression as fold change of the Spp1 gene in $TAM^{LV\ Spp1-KD}$ compared to $TAM^{LV\ Scramble}$. $TAM^{LV\ Spp1-KD}$ showed reduced Spp1 expression compared to $TAM^{LV\ Scramble}$ (n=5/condition). **G)** Immunofluorescence transversal sections of sarcoma tumor from control (CTRL) and $TAM^{LV\ Scramble}$, $TAM^{LV\ Spp1-KD}$-treated mice stained with NF200 (green) and DAPI (blue). TAMs are shown in red. Scale bar=5mm. Insets show the presence of transplanted $TAM^{LV\ Scramble}$, $TAM^{LV\ Spp1-KD}$ in sarcoma tumor. Labeled fibers in the sections are reported as right-side images. Scale bar=25 µm. **H)** Expression as fold change of the Spp1 gene in TAM +Parecoxib compared to TAM. TAM +Parecoxib showed reduced Spp1 expression compared to TAM (n=3/condition). Statistical analysis was performed via Student t-test.

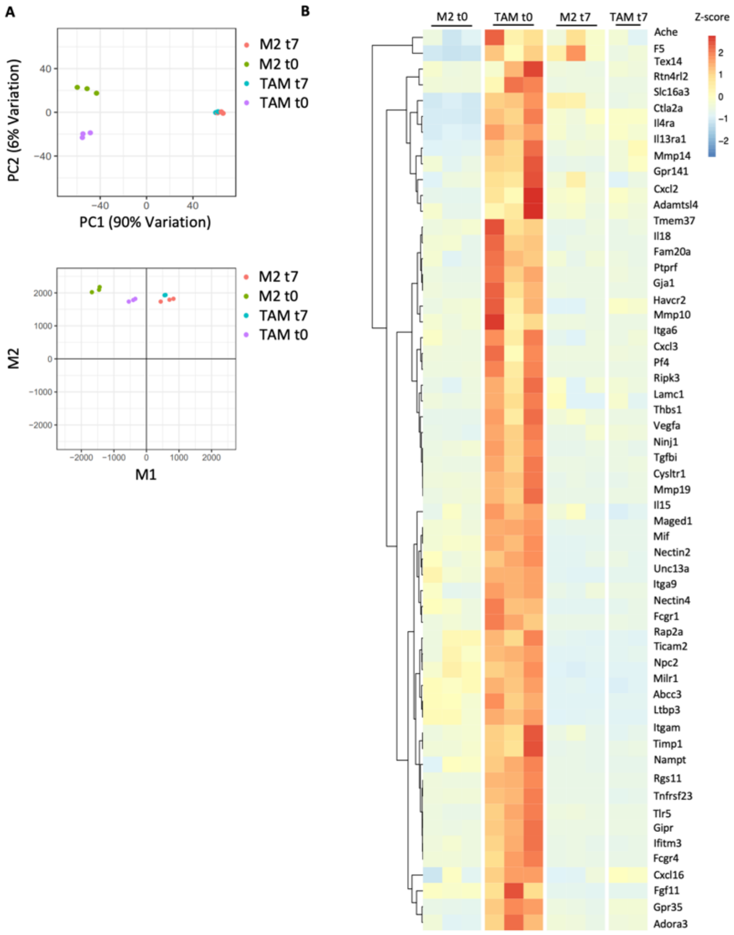

A
B
PC2 (6% Variation)
PC1 (90% Variation)
M2 t7
M2 t0
TAM t7
TAM t0
M2
M1
M2 t0
TAM t0
M2 t7
TAM t7
Z-score
Ache
F5
Tex14
Rtn4rl2
Slc16a3
Ctla2a
Il4ra
Il13ra1
Mmp14
Gpr141
Cxcl2
Adamtsl4
Tmem37
Il18
Fam20a
Ptprf
Gja1
Havcr2
Mmp10
Itga6
Cxcl3
Pf4
Ripk3
Lamc1
Thbs1
Vegfa
Ninj1
Tgfbi
Cysltr1
Mmp19
Il15
Maged1
Mif
Nectin2
Unc13a
Itga9
Nectin4
Fcgr1
Rap2a
Ticam2
Npc2
Milr1
Abcc3
Ltbp3
Itgam
Timp1
Nampt
Rgs11
Tnfrsf23
Tlr5
Gipr
Ifitm3
Fcgr4
Cxcl16
Fgf11
Gpr35
Adora3


FIGURE S4

**Legend to Supplementary Figure 4: Transcriptomic analysis of TAM and M2 macrophages before and following transplantation in scSCI.**

**A)** PCA on M2 before transplantation (t0) (green), TAM t0 (violet), M2 7 days after transplantation (t7) (orange), and TAM t7 (blue) samples. Each dot represents a sample. TAM t0 were characterized by a distinct gene expression profile compared to M2 t0. TAM t7 and M2 t7 showed comparable transcriptomic signature. Score plot on M2 t0 (green), TAM t0 (violet), M2 t7 (orange), and TAM t7 (blue) samples referred to M1. Each dot represents a sample. TAM t7 and M2 t7 showed comparable transcriptomic signatures closely related to the inflammatory M1 macrophages, which were different from their respective t0 gene profiles. M1 and M2 samples characterization was performed taking into consideration M1 and M2 signatures defined in Setten et al. Signatures were used in the single sample Gene Set Enrichment Analysis (performed with GSVA v.1.30.0) in order to evaluate gene signature scores for each sample. **B)** Z-score heatmap gene expression related to TAM t0 in comparison with M2 t0, M2 t7 and TAM t7 samples. T0 TAM receptome/secretome were characterized by the up-regulation of 58 genes.

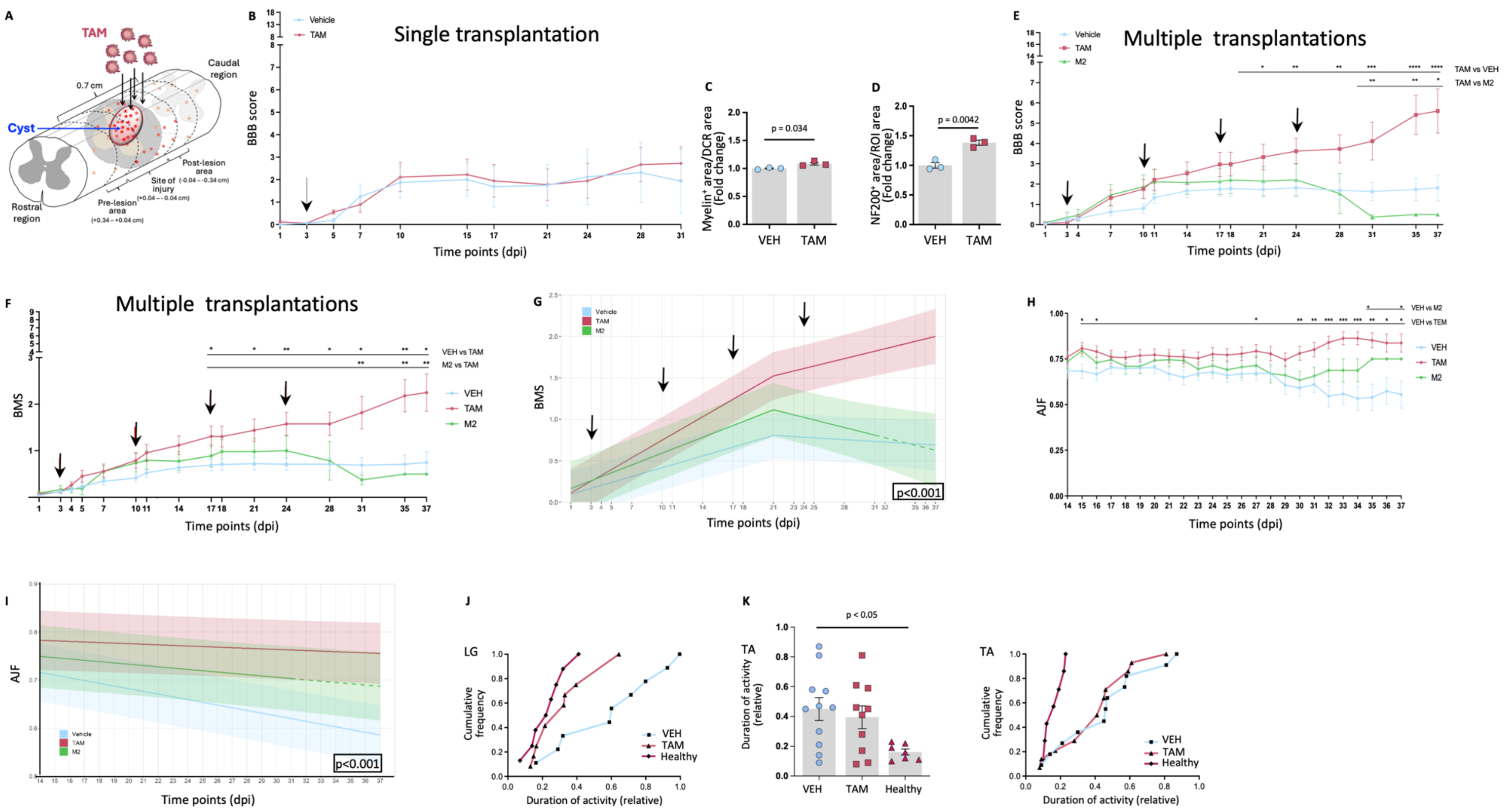
A
TAM
Cyst
0.7 cm
Caudal region
Rostral region
Post-lesion area
Site of injury (-0.04 – -0.34 cm)
Pre-lesion area (+0.34 – +0.04 cm)
B
Single transplantation
Vehicle
TAM
BBB score
Time points (dpi)
C
Myelin+ area/DCR area (Fold change)
p = 0.034
VEH
TAM
D
NF200+ area/ROI area (Fold change)
p = 0.0042
VEH
TAM
E
Multiple transplantations
Vehicle
TAM
M2
TAM vs VEH
TAM vs M2
BBB score
Time points (dpi)
F
Multiple transplantations
VEH vs TAM
M2 vs TAM
VEH
TAM
M2
BMS
Time points (dpi)
G
Vehicle
TAM
M2
BMS
p<0.001
Time points (dpi)
H
VEH vs M2
VEH vs TEM
VEH
TAM
M2
AJF
Time points (dpi)
I
AJF
Vehicle
TAM
M2
p<0.001
Time points (dpi)
J
LG
Cumulative frequency
Duration of activity (relative)
VEH
TAM
Healthy
K
TA
Duration of activity (relative)
p < 0.05
VEH
TAM
Healthy
TA
Cumulative frequency
Duration of activity (relative)
VEH
TAM
Healthy

FIGURE S5

**Legend to Supplementary Figure 5: TAM adoptive transfer promotes partial locomotor recovery in scSCI.**

**A)** Schematic drawing illustrating the experimental design of TAM transplantation in T11 severe spinal cord injury (SCI) model. A severe SCI was applied at T11 vertebral level to injure all descending pathways. TAM/VEH were transplanted/injected in 4 different points (indicated by the black arrows) close to the lesion site. The circled red area shows the cyst, and red asterisks represent transplanted TAM spread into the spinal cord parenchyma. **B)** Line plot of the mean BBB score of VEH- and TAM-treated scSCI mice at 31 dpi which underwent a single transplant showing a similar locomotor recovery (VEH: n=7; TAM: n=9). **C)** Percentage, expressed as fold change respect to VEH-treated scSCI mice, of the myelin$^{+}$ area normalized for the total area of the DCR in VEH- and TAM-treated scSCI mice that underwent only one injection/transplantation. After one transplant, TAM-treated scSCI mice had an increased myelin$^{+}$ area content compared to VEH scSCI mice (n=3/condition). **D)** Percentage, expressed as fold change respect to VEH-treated scSCI mice, of the NF200$^{+}$ area normalized for the total area of the ROI considered in VEH- and TAM-treated scSCI mice that underwent only one injection/transplantation. After one transplant, TAM-treated scSCI mice had an increased NF200$^{+}$ area content compared to VEH scSCI mice (n=3/condition). **E)** Line plot of the mean BBB score of VEH-, TAM-, and M2-treated scSCI mice at 37 dpi that underwent multiple injections/transplantations. TAM-treated scSCI mice show a significant increased locomotor recovery (VEH: n=16; TAM: n=10; M2: n=2). **F)** Line plot of the mean BMS score of VEH-, TAM-, and M2-treated scSCI mice at 37 dpi that underwent multiple injections/transplantations. TAM-treated scSCI mice show a significant increased locomotor recovery (VEH: n=16; TAM: n=10; M2: n=2). Black arrows indicate the days of the injection/transplantation. **G)** Mixed-effect regression model of the BMS response (estimated marginal means) with confidence bands in VEH- (light blue), TAM- (red), and M2-treated (green) scSCI mice. The three lines showed significantly different slopes indicating TAM role in locomotor recovery of scSCI mice (VEH: n=36; TAM: n=33; M2: n=27; see supplementary information for additional statistical information). Black arrows indicate the days of injection/transplantation. **H)** Line plot of the mean Ankle Joint Flexibility (AJF) score of VEH-, TAM- and M2-treated scSCI mice at 37 dpi that underwent multiple injections/transplantations. TAM-treated scSCI mice had a better condition of the ankle joint compared to VEH- and M2-treated mice (VEH: n=16; TAM: n=10; M2: n=2). **I)** Mixed-effect regression model of the AJF response (estimated marginal means) with confidence bands in VEH-, TAM- and M2-treated scSCI mice. The three lines showed significantly different slopes (VEH: n=36; TAM: n=33; M2: n=27; see supplementary information for additional statistical information); **J)** Cumulative frequency analysis of *Lateral Gastrocnemius* (LG) muscle. Values from TAM-treated scSCI mice and from healthy mice are similarly distributed, compared to VEH-treated scSCI mice that instead show longer durations of activity. **K)** (Left) Graph showing the Duration of *Tibialis Anterior* (TA) muscle activity relative to the total duration of recording in VEH- and TAM-treated scSCI mice and in healthy mice (VEH: n=11; TAM: n=10; Healthy: n=7). A value of 1.0 corresponds to a muscle constantly active throughout the recording session. (Right) Graph showing the cumulative frequency analysis. Treatment of scSCI mice with TAM did not restore the duration of muscle activity to normal (Healthy mice). Data are expressed as mean±SEM. Statistical differences were evaluated by unpaired Student t-test and 1-way ANOVA.

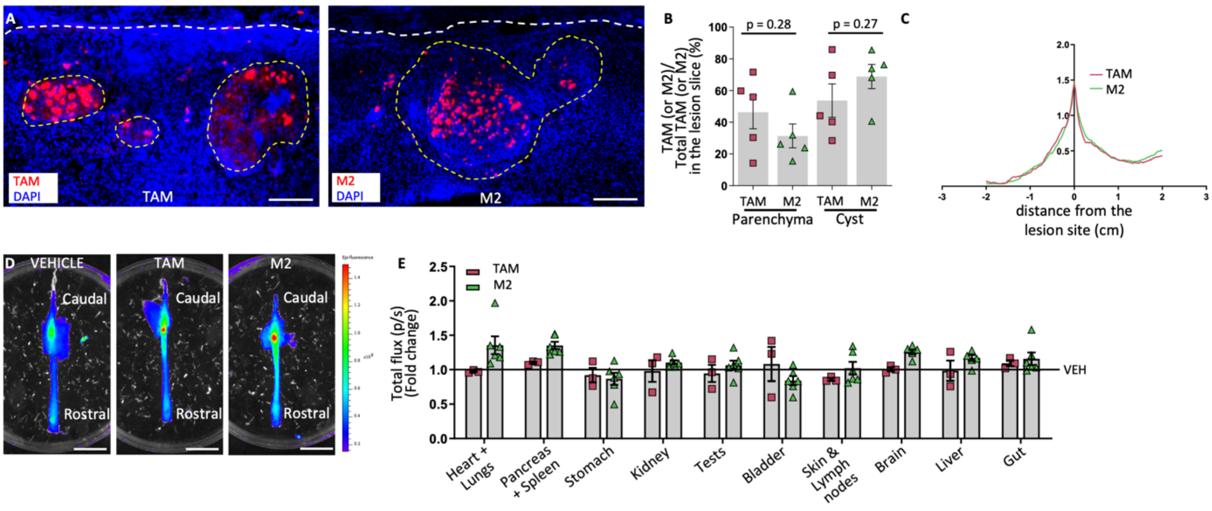
A
TAM
DAPI
TAM
M2
DAPI
M2
B
p = 0.28
p = 0.27
TAM (or M2)/
Total TAM (or M2)
in the lesion slice (%)
100
80
60
40
20
0
TAM
M2
TAM
M2
Parenchyma
Cyst
C
2.0
1.5
1.0
0.5
TAM
M2
-3
-2
-1
0
1
2
3
distance from the
lesion site (cm)
D
VEHICLE
TAM
M2
Caudal
Caudal
Caudal
Rostral
Rostral
Rostral
E
TAM
M2
Total flux (p/s)
(Fold change)
2.5
2.0
1.5
1.0
0.5
0.0
VEH
Heart +
Lungs
Pancreas
+ Spleen
Stomach
Kidney
Tests
Bladder
Skin &
Lymph
nodes
Brain
Liver
Gut

FIGURE S6

**Legend to Supplementary Figure 6: After multiple transplantation TAM and M2 show a similar distribution in the spinal cord injured parenchyma and in the visceral organs.**

**A)** Magnification from an immunofluorescence image of longitudinal spinal cord sections of TAM- and a M2-treated scSCI mouse stained for tdTomato (red) and DAPI (blue). The dotted white line delineates the cyst area. TAM and M2 are shown in red. Scale bar=200µm. **B)** Percentage of the distribution of TAM and M2 in the transversal lesioned spinal cord sections of TAM- and a M2-treated scSCI mouse at 37 dpi normalized for the total number of TAM (or M2, respectively) in the selected spinal cord sections. TAM and M2 present a similar distribution into the cyst area (Cyst) and the surrounding parenchyma (Parenchyma) (TAM_Parenchyma: n=5; M2_Parenchyma: n=3; TAM_Cyst: n=5; M2_Cyst: n=3). **C)** Mean tdTomato TAM (red) and tsTomato M2 (green) caudo-rostral fluorescence distribution along the spinal cord analyzed by IVIS Spectrum. Data are expressed as values/$10^8$. tdTomato TAM and tdTomato M2 fluorescence showed a similar distribution which is higher at the lesion site and reached a lesser extent in the rostral and causal region (TAM n=3; M2 n=6). **D)** Representative IVIS Spectrum images of VEH-, TAM- and M2-treated scSCI spinal cords at 37 dpi showing tdTomato TAM and tdTomato M2 fluorescence at the lesion site (site of TAM and M2 repeated transplantation) and less fluorescence in VEH-treated SCI spinal cord. Scale bar=1cm. **E)** Quantitative analysis, expressed as fold change relative to VEH, of the emitted fluorescence in the visceral organs of tdTomato TAM- and tdTomato M2-treated scSCI mice analyzed by IVIS Spectrum. TAM- and M2 in respectively TAM and M2-treated scSCI mice, showed a similar distribution in the visceral organs. Data are expressed as mean±SEM. Statistical differences were evaluated by unpaired Student t test.

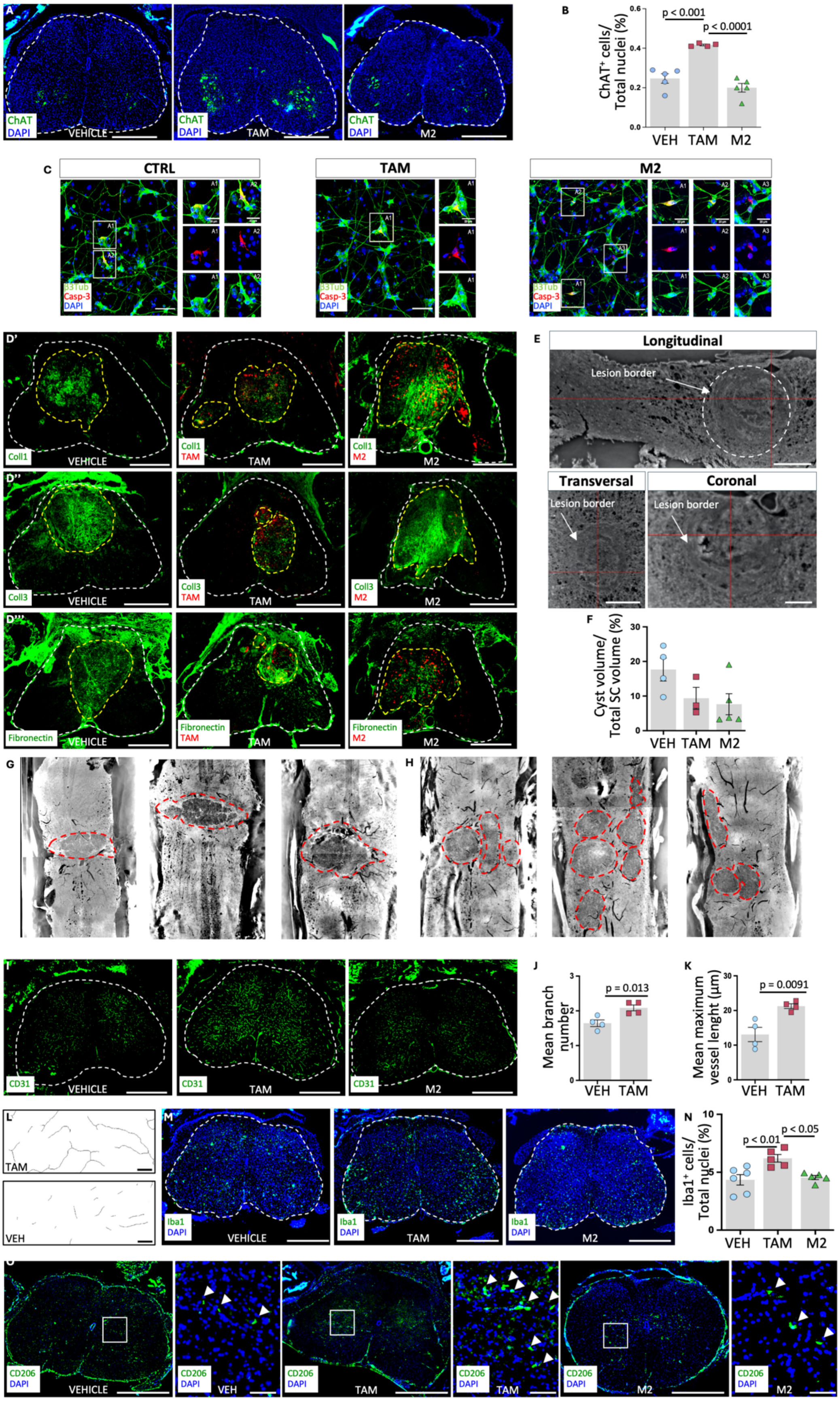
A
ChAT
DAPI
VEHICLE
TAM
M2
B
p < 0.001
p < 0.0001
ChAT+ cells/
Total nuclei (%)
VEH
TAM
M2
C
CTRL
TAM
M2
β3Tub
Casp-3
DAPI
D'
Coll1
VEHICLE
TAM
M2
D''
Coll3
D'''
Fibronectin
E
Longitudinal
Lesion border
Transversal
Coronal
F
Cyst volume/
Total SC volume (%)
G
H
I
CD31
J
p = 0.013
Mean branch
number
K
p = 0.0091
Mean maximum
vessel lenght (μm)
L
M
Iba1
N
p < 0.01
p < 0.05
Iba1+ cells/
Total nuclei (%)
O
CD206
VEH

FIGURE S7

**Legend to Supplementary Figure 7: TAM adoptive transfer modulate the SCI-induced hostile microenvironment.**

**A)** Immunofluorescence images of transversal spinal cord sections of VEH-, TAM- and M2-treated scSCI mice stained with ChAT (green) and DAPI (blue). The dotted white line delineates the section area. Scale bar=500µm. **B)** Percentage of ChAT$^+$ cells normalized for the total nuclei of the section in VEH-, TAM- and M2-treated scSCI mice, showing an increased ChAT$^+$ cells content in TAM-treated scSCI mice compared to all the other groups (VEH: n=5; TAM: n=4; M2: n=6). **C)** Immunofluorescence images of human iPSC-derived MNs (CTRL), iPSC-derived MNs +TAM, and iPSC-derived MNs +M2, stained with β3Tub (green), Caspase-3 (red), and DAPI (blue). Scale bar=10µm. **D')** Immunofluorescence images of transversal spinal cord sections of VEH-, TAM- and M2-treated scSCI mice stained with Coll1 (green). The dotted white line delineates the section area. The dotted yellow line delineates the cyst area. TAM and M2 are shown in red. Scale bar=500µm. **D'')** Immunofluorescence images of transversal spinal cord sections of VEH-, TAM- and M2-treated scSCI mice stained with Coll3 (green). The dotted white line delineates the section area. The dotted yellow line delineates the cyst area. TAM and M2 are shown in red. Scale bar=500µm. **D''')** Immunofluorescence images of transversal spinal cord sections of VEH-, TAM- and M2-treated scSCI mice stained with Fibronectin (green). The dotted white line delineates the section area. The dotted yellow line delineates the cyst area. TAM and M2 are shown in red. Scale bar=500µm. **E)** Representative magnification of longitudinal, transversal, and coronal sections of SCI tissue obtained by X-ray tomography, showing the lesion border. Scale bar=10µm. **F)** Mean cyst volume (%), normalized to the total volume of the spinal cord parenchyma analyzed, in VEH-, TAM- and M2-treated scSCI mice, showing cyst volume is not significantly different in the 3 groups (VEH: n=4; TAM: n=3; M2: n=5). **G)** Representative images of longitudinal sections of VEH scSCI mice obtained by X-ray tomography. VEH mice showed a cyst as a single structure. Red dotted line delineates the cyst. **H)** Representative images of longitudinal sections of TAM-treated scSCI mice were obtained by X-ray tomography. TAM-treated mice showed cysts formed by smaller multilobes. Red dotted line delineates the cyst. **I)** Immunofluorescence images of transversal spinal cord sections of VEH-, TAM-, and M2- treated scSCI mice stained with CD31 (green). The dotted white line delineates the section area. Scale bar=500µm. **J)** Mean branch number of vessels in VEH- and TAM-treated scSCI mice, showing higher mean branch length in TAM-treated scSCI mice compared to VEH-treated scSCI mice (n=4/condition). **K)** Mean maximum vessel length in VEH- and TAM-treated scSCI mice, showing higher maximum vessel length in TAM-treated scSCI mice compared to VEH-treated scSCI mice (n=4/condition). **L)** Vessel skeletonization in VEH and TAM-treated SCI mice was obtained using the vessel morphometric analysis of the angiogenic pattern and it was used to quantify branch number and lenght. Scale bar=20µm. **M)** Immunofluorescence images of transversal spinal cord sections of VEH-, TAM- and M2-treated scSCI mice stained with Iba1 (green) and DAPI (blue). The dotted white line delineates the section area. Scale bar=500µm. **N)** Percentage of Iba1$^+$ cells normalized for the total nuclei of the section in VEH-, TAM- and M2-treated scSCI mice. TAM-treated group showed increased $Iba1^+$ cell content compared to all the other group (VEH: n=6; TAM: n=5; M2: n=5). **O)** (Left) Immunofluorescence images and (right) relative magnifications of transversal spinal cord sections of VEH-, TAM- and M2-treated scSCI mice stained with CD206 (green) and DAPI (blue). Scale bar=500µm. (Right) Magnifications of the white square. White arrows indicate $CD206^+$ cells-sc.White arrows indicate $CD206^+$ cells. Scale bar=50µm. Data are expressed as mean±SEM. Statistical differences were evaluated by unpaired Student t-test and 1-way ANOVA.

A

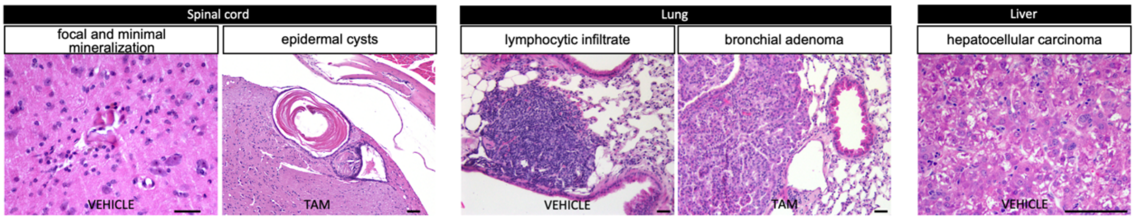


B

Non-tumoral lesions

| **Organ: lesion** | **Vehicle** | **TAM injected** |
|---|---|---|
| Lungs: inflammatory cell infiltrates | 11/12 | 9/10 |
| Lungs: acidophilic macrophage pneumonia | 1/12 | 0/10 |
| Liver: steatosis | 1/12 | 1/10 |
| Liver: inflammatory cell infiltrates | 7/12 | 8/10 |
| Spleen: white pulp hyperplasia | 10/12 | 8/10 |
| Kidneys: inflammatory cell infiltrates | 9/12 | 7/10 |
| Kidneys: glomerulopathy | 0/12 | 2/10 |
| Pancreas: inflammatory cell infiltrates | 4/12 | 6/10 |

C

Tumoral lesions

| **Organ: tumor** | **Vehicle** | **TAM injected** |
|---|---|---|
| Lungs: bronchial adenoma | 1/12 | 1/10 |
| Liver: hepatocellular adenoma | 0/12 | 1/10 |
| Liver: hepatocellular carcinoma | 0/12 | 1/10 |
| Lymph nodes: lymphoma | 1/12 | 0/10 |

FIGURE S8

**Legend to Supplementary Figure 8: Long term pre-clinical safety assessment of TAM adoptive transfer.**

**A)** Representative magnification of hematoxylin & eosin (H&E) images of VEH- and TAM-treated scSCI mice visceral organs, showing non-tumoral and tumoral lesions. Scale bars=40µm. **B)** Summary of non-tumoral lesions observed in VEH- and TAM-treated mice (no lesion) enrolled in the long-term study (1 year). **C)** Summary of tumoral lesions observed in VEH- and TAM-treated mice (no lesion) enrolled in the long-term study (1 year).

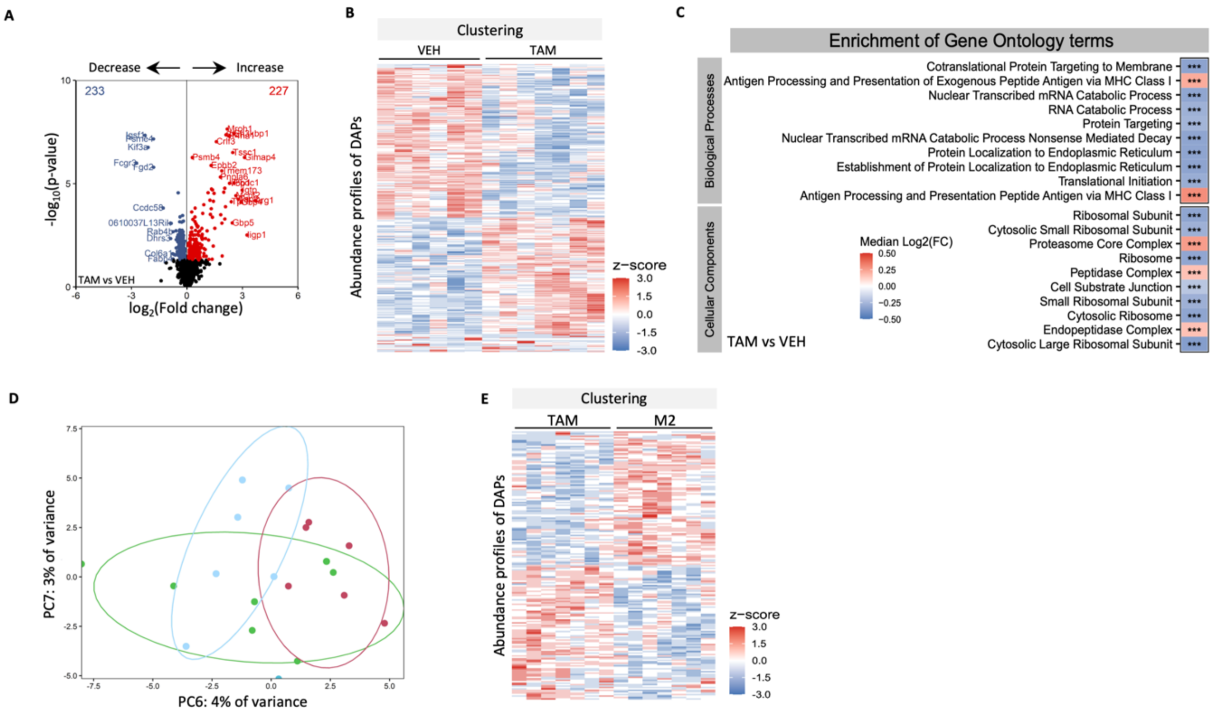

A
Decrease
Increase
233
227
-log10(p-value)
log2(Fold change)
TAM vs VEH
B
Clustering
VEH
TAM
Abundance profiles of DAPs
z-score
3.0
1.5
0.0
-1.5
-3.0
C
Enrichment of Gene Ontology terms
Biological Processes
Cotranslational Protein Targeting to Membrane
Antigen Processing and Presentation of Exogenous Peptide Antigen via MHC Class I
Nuclear Transcribed mRNA Catabolic Process
RNA Catabolic Process
Protein Targeting
Nuclear Transcribed mRNA Catabolic Process Nonsense Mediated Decay
Protein Localization to Endoplasmic Reticulum
Establishment of Protein Localization to Endoplasmic Reticulum
Translational Initiation
Antigen Processing and Presentation Peptide Antigen via MHC Class I
Cellular Components
Ribosomal Subunit
Cytosolic Small Ribosomal Subunit
Proteasome Core Complex
Ribosome
Peptidase Complex
Cell Substrate Junction
Small Ribosomal Subunit
Cytosolic Ribosome
Endopeptidase Complex
Cytosolic Large Ribosomal Subunit
Median Log2(FC)
TAM vs VEH
D
PC7: 3% of variance
PC6: 4% of variance
E
Clustering
TAM
M2
Abundance profiles of DAPs
z-score


### FIGURE S9

**Legend to Supplementary Figure 9: Proteomic analysis reveals differences in TAM-treated scSCI parenchyma compared to VEH- and M2-treated scSCI parenchyma.**

**A)** Volcano plot showing differential protein abundance in the protein extracts from TAM- and VEH-treated scSCI spinal cords. Proteins are separated according to their log2-transformed intensity fold-change (TAM/VEH; x-axis) and their -log10-transformed p-value (y-axis), (TAM: n=7; VEH: n=6). Protein extracts from TAM-treated scSCI spinal cords showed 227 up-regulated proteins and 233 down-regulated proteins compared to protein extracts from VEH-treated scSCI spinal cords. **B)** Clustering of the log2(Abundances) of DAPs between TAM and VEH-treated scSCI mice. **C)** Affected biological processes and cellular components of the Gene Ontology database between VEH- and TAM-treated scSCI mice. **D)** PCA on protein extracts from VEH-, TAM- and M2-treated scSCI spinal cords (VEH: n=6; TAM: n=7; M2: n=7). Each dot represents a sample. **E)** Clustering of the $log_2$(Abundances) of the DAPs between TAM- and M2-treated scSCI mice.

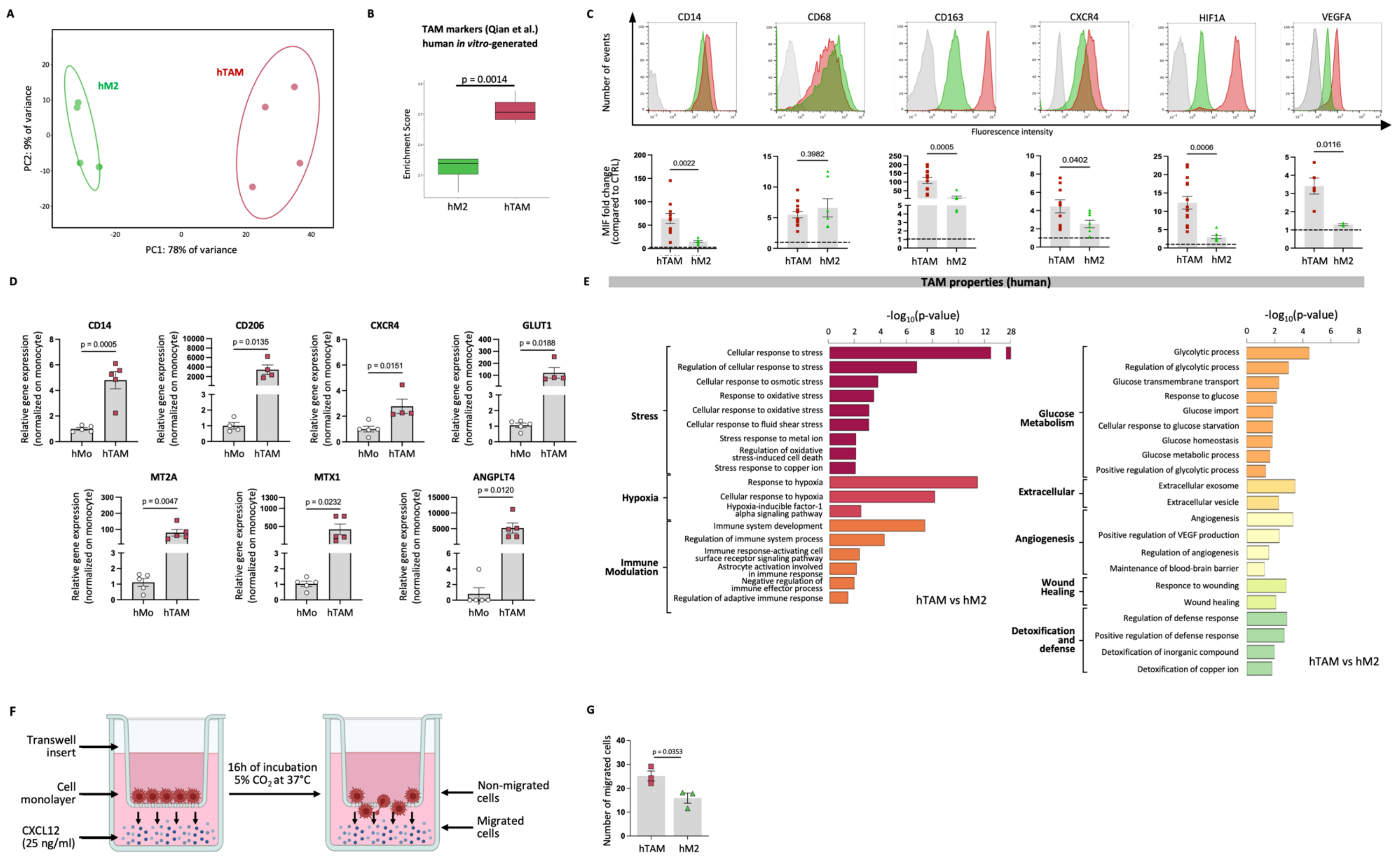
A
hM2
hTAM
PC2: 9% of variance
PC1: 78% of variance
B
TAM markers (Qian et al.)
human in vitro-generated
p = 0.0014
Enrichment Score
hM2
hTAM
C
CD14
CD68
CD163
CXCR4
HIF1A
VEGFA
Number of events
Fluorescence intensity
MIF fold change
(compared to CTRL)
0.0022
0.3982
0.0005
0.0402
0.0006
0.0116
hTAM hM2
D
CD14
p = 0.0005
CD206
p = 0.0135
CXCR4
p = 0.0151
GLUT1
p = 0.0188
MT2A
p = 0.0047
MTX1
p = 0.0232
ANGPLT4
p = 0.0120
Relative gene expression
(normalized on monocyte)
hMo hTAM
E
TAM properties (human)
-log10(p-value)
Stress
Cellular response to stress
Regulation of cellular response to stress
Cellular response to osmotic stress
Response to oxidative stress
Cellular response to oxidative stress
Cellular response to fluid shear stress
Stress response to metal ion
Regulation of oxidative stress-induced cell death
Stress response to copper ion
Hypoxia
Response to hypoxia
Cellular response to hypoxia
Hypoxia-inducible factor-1 alpha signaling pathway
Immune Modulation
Immune system development
Regulation of immune system process
Immune response-activating cell surface receptor signaling pathway
Astrocyte activation involved in immune response
Negative regulation of immune effector process
Regulation of adaptive immune response
hTAM vs hM2
Glucose Metabolism
Glycolytic process
Regulation of glycolytic process
Glucose transmembrane transport
Response to glucose
Glucose import
Cellular response to glucose starvation
Glucose homeostasis
Glucose metabolic process
Positive regulation of glycolytic process
Extracellular
Extracellular exosome
Extracellular vesicle
Angiogenesis
Angiogenesis
Positive regulation of VEGF production
Regulation of angiogenesis
Maintenance of blood-brain barrier
Wound Healing
Responce to wounding
Wound healing
Detoxification and defense
Regulation of defense response
Positive regulation of defense response
Detoxification of inorganic compound
Detoxification of copper ion
hTAM vs hM2
F
Transwell insert
Cell monolayer
CXCL12 (25 ng/ml)
16h of incubation
5% CO2 at 37°C
Non-migrated cells
Migrated cells
G
Number of migrated cells
p = 0.0353
hTAM hM2

FIGURE S10

**Legend to Figure Supplementary 10: Human *in vitro*-generated TAM display *bona fide* TAM signature.**

**A)** PCA on *in vitro*-generated human M2 (hM2, green) and TAM (hTAM, red) (n=4/condition). Each dot represents a sample. **B)** Single sample gene set enrichment analysis of TAM hallmarks defined by Qian et al. in *in vitro*-generated hM2 and hTAM, showing enriched expression in TAM group. **C)** (Upper row) Representative hTAM and hM2 flow cytometry analysis of CD14, CD68, CD163, CXCR4, HIF-1α, and VEGFA markers. (Bottom row) Barplots, expressed as fold-change to an unstained control sample, of the mean fluorescence intensity (MIF) of CD14, CD163, CXCR4, HIF-1α, and VEGFA markers in hTAM and hM2. hTAM showed higher expression of CD14, CD163, CXCR4, HIF-1α, and VEGFA markers compared to hM2 and a similar expression of CD68 marker. **D)** Barplots, expressed as fold-change respect to human monocytes, of the relative gene expression of CD14, CD206, CXCR4, GLUT1, MT2A, MTX1, and ANGPLT4 in human monocytes and hTAM. hTAM showed higher expression of all selected genes compared to human monocytes. **E)** Barplots showing enriched upregulated gene ontologies in hTAM compared to hM2. Barplot length is the $-\log_{10}$(p-value) returned from the enrichment analysis. **F)** Schematic representation of the transwell migration assay. (Left) A transwell insert was placed in the well of a multiwell plate forming a migration chamber composed by an upper and lower compartment separated by a microporous membrane. (Right) In presence of CXCL12 in the culture medium, cells actively migrate from the upper to the lower compartment. **G)** Number of hTAM and hM2 migrated cells in presence of 25 ng/ml CXCL12 in the culture medium (n=3/condition). hTAM showed higher motility compared to hM2. Data are expressed as mean ± SEM. Statistical analysis was performed via Student t-test.

#### Supplementary table

**Table S1**

**"Neural growth" gene list**

| | |
|---|---|
| APLP2 | NAP1L1 |
| ASAH1 | PFN1 |
| CD63 | PGK1 |
| CDC42 | RPS6 |
| CLIC1 | S100A6 |
| GRN | SDCBP |
| HMOX1 | SOD2 |
| ITMB2 | SPP1 |
| LGALS1 | UBE2B |
| MCL1 | VASP |
| MEF2C | VCAN |

**Legend to Table S1:**

Genes included in the "neural growth" gene list. Genes are reported with official symbol names.

**Table S2a**

**Basso, Beattie and Bresnahan locomotor rating scale (BBB):**

| | | **Dpi < 17** | **Dpi ≥ 17** |
|---|---|---|---|
| | Intercept | Slope | Slope |
| **VEH** | -0.107 (-0.307, 0.094)<br>p = 0.297 | 0.113 (0.066, 0.160)<br>p < 0.001 | -0.003 (-0.029, 0.023)<br>p = 0.832 |
| | | **Dpi < 8** | **Dpi ≥ 8** |
| | Intercept | Slope | Slope |
| **M2** | -0.503 (-0.866, -0.140)<br>p = 0.007 | 0.317 (0.141, 0.494)<br>p < 0.001 | -0.008 (-0.043, 0.027)<br>p = 0.667 |
| | | **Dpi < 13** | **Dpi ≥ 13** |
| | Intercept | Slope | Slope |
| **TAM** | -0.363 (-0.603, -0.124)<br>p = 0.003 | 0.220 (0.118, 0.322)<br>p < 0.001 | 0.102 (0.054, 0.150)<br>p < 0.001 |
| | **21 dpi** | **37 dpi** | **57 dpi** |
| **VEH** | 1.81 (1.08 – 2.54) | 1.76 (0.90 – 2.62) | 1.71 (0.49 – 2.93) |
| **M2** | 1.93 (0.70 – 3.17) | 1.81 (0.30 – 3.32) | 1.66 (-0.37 – 3.68) |
| **TAM** | 3.31 (2.13 – 4.49) | 4.94 (3.40 – 6.48) | 6.98 (4.68 – 9.27) |

**Table S2a legend:** Our findings suggested that the nonlinear relationship between BBB and dpi, spanning from 1 to 37 dpi across the three treatment groups (Vehicle, M2, and TAM), could be effectively modelled using linear splines with a single knot. In other terms, we approximated this relationship using a mixed-effect linear model that incorporated a single change point.

For DPIs before the change point, the average BBB grew linearly in all three treatment groups, with rates of 0.113, 0.317, and 0.220 for Vehicle, M2, and TAM, respectively (all $p<0.001$). The differences between these three slopes were statistically significant ($p<0.001$).

For DPIs after the change point, the average BBB continued to grow linearly, but only in the TAM group, with a slope of 0.102 ($p<0.001$). Conversely, the M2 group and the Vehicle group showed trends that did not significantly differ from zero ($p=0.667$ and $p=0.832$). The slopes of these groups were significantly different from that of the TAM group ($p<0.001$). dpi: days post injury; p: p-value of the Wald test for the coefficient in the linear regression model; VEH: vehicle; TAM: tumor associated macrophages.

**Table S2b**

**Basso Mouse Scale (BMS):**

| | | **Dpi < 21** | **Dpi ≥ 21** |
|---|---|---|---|
| | Intercept | Slope | Slope |
| **VEH** | 0.062 (-0.015, 0.138)<br>p = 0.113 | 0.035 (0.022, 0.049)<br>p < 0.001 | -0.007 (-0.019, 0.004)<br>p = 0.221 |
| **TAM** | 0.043 (-0.061, 0.146)<br>p = 0.420 | 0.071 (0.047, 0.094)<br>p < 0.001 | 0.030 (0.002, 0.058)<br>p = 0.035 |
| **M2** | 0.118 (-0.023, 0.259)<br>p = 0.101 | 0.047 (0.020, 0.075)<br>p = 0.001 | -0.031 (-0.057, -0.004)<br>p = 0.022 |

dpi: Days post injury; p: p-value of the Wald test for the coefficient in the linear regression model; VEH: Vehicle; TAM: Tumor associated macrophages.

**Table S2b legend:** Our findings suggested that the nonlinear relationship between the Basso mouse scale (BMS) score and dpi, spanning from 1 to 37, across the three treatment groups (VEH, TAM, and M2), could be effectively modeled using linear splines with a single knot at 21 dpi. In other terms, we approximated this relationship using a mixed-effect linear model that incorporated a single change point.

Before the $21^{st}$ dpi, the average BMS score grew linearly in all groups, with rates of 0.035, 0.071, and 0.047 for VEH-, TAM-, and M2- treated spinal cord injured (SCI) mice, respectively (all $p\leq 0.001$). The differences between these three slopes were statistically significant ($p<0.001$).

This growth led to BMS score of 0.81 (95% Confidence interval (CI) 0.54-1.07) for the VEH-, 1.52 (95%CI 1.01-2.03) for the TAM-, and 1.11 (95%CI 0.52-1.71) for the M2-treated SCI mice, at 21 dpi ($p<0.001$).

After the $21^{st}$ dpi, the average BMS score continued to grow linearly only in the TAM group, with a slope of 0.030 ($p=0.035$), culminating in a BMS score of 2.00 (95%CI 1.43-2.57) at 37 dpi. Conversely, the M2 group showed a statistically significant negative trend with a slope of -0.031 ($p=0.022$), while the slope for the VEH group did not significantly differ from zero (p=0.22). The slopes of VEH and M2 groups were significantly different from that of the TAM group (p<0.001).

At 37 dpi the mean value of BMS score was 0.62 (95%CI 0.16-1.09) and 0.69 (95%CI 0.40-0.98) in the M2 and VEH groups, respectively.

**Table S2c**

**Ankle Joint Flexibility (AJF):**

| | **Dpi ≥ 14** | |
|---|---|---|
| | Intercept | Slope |
| **VEH** | 0.72 (0.65, 0.78)<br>p < 0.001 | -0.006 (-0.009, -0.002)<br>p = 0.004 |
| **TAM** | 0.78 (0.72, 0.84)<br>p < 0.001 | -0.001 (-0.003, 0.0008)<br>p = 0.244 |
| **M2** | 0.75 (0.69, 0.81)<br>p < 0.001 | -0.003 (-0.006, 0.0004)<br>p = 0.096 |

**Table S2c legend:** The trend over time of the ankle joint flexibility (AJF), although showing weak elements of nonlinearity, is mostly characterized by a linear component. Therefore, we have opted to model AJF score using a mixed-effect linear regression model (with no splines).

#### Supplementary legends to movies

**Legend to SMov. 1:**

Video showing open-field of a VEH-treated scSCI mouse at 37 days following severe contusive injury and repeated vehicle injections. VEH-treated mouse showed no movement of the hindlimb paws.

**Legend to SMov. 2:**

Video showing open-field of a TAM-treated scSCI mouse at 37 days following severe contusive injury and repeated TAM transplantations. TAM-treated mouse showed consistent plantar stepping.

**Legend to SMov. 3:**

Video showing 3D reconstruction from X-ray phase-contrast micro computed tomography images of the structure of the cyst from a VEH-treated mouse.

**Legend to SMov. 4:**

Video showing 3D reconstruction from X-ray phase-contrast micro-computed tomography images of the structure of the cysts from a TAM-treated mouse.

**Legend to SMov. 5:**

Video showing a 3D reconstruction of Z-maximum projection of a spinal cord X-ray phase-contrast micro computed tomography showing the vasculature tree in a VEH-treated spinal cord.

**Legend to SMov 6:**

Video showing a3D reconstruction of Z-maximum projection of the a spinal cord X-ray phase-contrast micro computed tomography showing the vasculature tree in a TAM-treated spinal cord.
